# An effect size-based screening approach to identify actigraphy markers of the submarine environment

**DOI:** 10.1101/2025.06.16.659930

**Authors:** Daniel Marques, Carina Fernandes, Nuno L. Barbosa-Morais, Cátia Reis

## Abstract

Identifying behavioral markers of disease risk associated with circadian disruption remains challenging due to reproducibility concerns, as human studies are often under-powered to detect the effects of interest. Submarines provide an extreme example of circadian disruption, depriving individuals of natural light-dark cycles and limiting opportunities for physical activity. To address these challenges, we developed an effect size-based method to robustly screen potential actigraphy-derived markers of the submarine environment. Actigraphy data was collected across the pre-mission, mission and post-mission periods. During the mission, crew members followed a 6-hours-on/6-hours-off split shift schedule, except for submariners with on-call duty or extended hours. Forty-one actigraphy variables were screened for their ability to differentiate between study periods and shifts. A Monte Carlo permutation test was used for this initial screening, followed by receiver operating characteristic (ROC) curve analysis for pairwise comparisons. Light-related features, the slopes of regression lines fitted to the daily predicted dim light melatonin onset (DLMO) trajectories, and Pearson correlation coefficients between each recording day and its corresponding predicted DLMO demonstrated the greatest potential to distinguish between study periods. Most high-performing features discriminated between split shift work schedules and on-call or extended-hours schedules. Only the relative amplitude (*RA*) and the average light intensity during the five least-illuminated hours of the day (*L5*) reliably distinguished between the shift performing the bulk of nighttime work and its counterpart. Although submarine settings offer an opportunity to study a healthy population living under confined movement and dim light, detecting biological signals through actigraphy in this environment is inherently challenging. Demonstrating robust effects under these conditions supports the use of actigraphy in dimly-lit, isolated, confined, and extreme (DICE) environments, but also its applicability to study other operational or clinical populations with reduced activity levels.

## Introduction

The need to ensure the reproducibility of scientific findings has been recognized as a major concern across biology, medicine, and psychology (Kong & Francks, 2020). Poor reproducibility has been attributed to several factors, including the underreporting of effect sizes and insufficient statistical power (Maher et al., 2013). The emphasis on statistical significance and *p*-values has the unintended consequence of inflating effect sizes estimates (Ioannidis, 2008). Moreover, inadequate statistical power reduces the likelihood that a statistically significant result actually represents a true effect (Maher et al., 2013). This does not imply that studies with small sample sizes should be dismissed out-right (Quinlan, 2013). When a significant result is obtained from a small sample, provided that appropriate correction for the false positive rate (FPR) is used, it indicates that the estimated effect size is larger than the equivalent effect obtained using a larger sample (Friston, 2012). Thus, the exclusion of studies solely on the basis of small sample size constitutes what has been described as the fallacy of classical inference (Friston, 2012). Additionally, the presence of outliers in studies with small samples does not necessarily compromise their usefulness, as outliers tend to reduce the probability of incorrectly rejecting the null hypothesis (Type I errors) in parametric tests (Friston, 2012). This occurs because outliers influence the sample variance more strongly than the sample mean (Friston, 2012).

A practical step toward improving reproducibility is to focus on effect size quantification, which captures the magnitude of relationships or treatment effects (Maher et al., 2013; Sullivan & Feinn, 2012). To illustrate the potential of an effect size-based strategy to extract biological information from human behavioral data, we analyzed an actigraphy dataset from a military submarine mission using an actigraphy feature-screening approach based on effect size. Actigraphy, which continuously monitors movement, light and external temperature data from a device worn on the wrist, is used to assess circadian rhythms, near 24-hour endogenous oscillations that control behavior, physiology, and metabolism, allowing the organism to optimally manage bodily needs in response to predictable changes in the environment (Boivin & Boudreau, 2014; Cajochen & Schmidt, 2025). Submarines constitute a challenging environment for actigraphy data collection due to the restricted range of movement and low light levels. Our effect feature-screening approach identifies actigraphy-derived features, such as time spent above a given light threshold and average activity levels, that may serve as potential behavioral markers of submarine and submarine-like conditions.

In humans and other primates, light is the main entrainment cue (*zeitgeber),* since it entrains the phase of the suprachiasmatic nucleus (SCN) with the natural light-dark cycle (Cajochen & Schmidt, 2025). Non-photic *zeitgebers*, such as physical activity and meals, are also able to entrain both the SCN and peripheral circadian clocks (Peters et al., 2024). Mistimed and irregular exposure to photic and non-photic zeitgebers, such as in the submarine milieu, leads to circadian disruption (Meléndez-Fernández et al., 2023). Military submariners are required to operate in a dimly lit, isolated, confined and extreme (DICE) environment, for periods that can span several months (Nieuwenhuys et al., 2021; Zivi et al., 2020). To keep continuous operations, crews perform skilled and demanding tasks during the biological night, when the endogenous circadian system promotes sleep (Marando et al., 2023; McHill et al., 2014). In addition, light intensity levels are potentially insufficient to promote effective entrainment, and opportunities for consolidated sleep are slim, with rest periods often diverted to meals, personal care and recreational activities (Nieuwenhuys et al., 2021). The prolonged social isolation and reduced opportunities for physical activity may further contribute to worse physical and mental health outcomes and well-being (Nieuwenhuys et al., 2021).

Shift work in DICE environments accentuates many characteristics of night shift work (NSW), and it may also exacerbate its long-term health effects (Marando et al., 2023). NSW has been associated with multiple health conditions, including cancer and cardiometabolic disorders (Ansu Baidoo & Knutson, 2023; Erren et al., 2019). Proposed mechanisms to explain this association point to circadian desynchrony, including circadian misalignment, chronic partial sleep deprivation and melatonin suppression by nocturnal light exposure (Smith & Eastman, 2012). NSW uncouples *zeitgeber* exposure from internal time, inducing a state of internal desynchrony among circadian clocks in different tissues and organs (Heyde & Oster, 2022). Compared to NSW, the health effects of sub-marine shift work remain poorly characterized, limiting the ability to assess whether any alterations persist beyond the mission (Marando et al., 2023).

Given that similarity and considering that submariners constitute a healthy population with restricted movement, the submarine environment provides an ideal setting to introduce our novel effect size-based strategy for identifying actigraphy-derived markers of shift work and to validate the use of actigraphy in conditions of low light and limited mobility.

## Materials and methods

### Participants and actigraphy data collection

Actigraphy data from a group of twenty-nine Portuguese military submariners (mean age 35.6 ± 4.9 years) was collected to test the feature-screening methodology proposed in this study. Participants were mostly male (26/29), and most were quite experienced both in active duty (15.5 ± 5.1 years) and in the submarine fleet (6.0 ± 5.6 years). Data collection was conducted as part of a broader research project approved by the Ethics Committee of the NOVA National Public Health School, NOVA University Lisbon, Portugal (CE-ENSP n°12/2022). All participants provided informed consent prior to enrollment. Psychological variables from the same cohort have been previously published elsewhere (Fernandes et al., 2025). Inclusion was limited to individuals actively serving as crew members during the mission, and those with a previous diagnosis of a sleep disorder or taking chronic medication affecting sleep or wakefulness were excluded.

Data acquisition window spanned three consecutive periods: pre-mission (P1), mission (P2), and post-mission (P3) [**Figure 1**]. Throughout the eight-week acquisition window, submariners wore an ActTrust2 actigraphy monitoring device (Condor Instru-ments™, Brazil) on their non-dominant wrist. Movement (Proportional Integral Mode (PIM)), skin temperature (°C), and white light (lx) were recorded at a sampling rate of sixty seconds. Spurious periods of activity were manually removed at the start and end of the actigraphy recordings using the ActStudio® software (version 2.2.2). Off-wrist periods of the actigraphy trace were identified through visual inspection and marked as miss-ing values.

**Figure 1.**
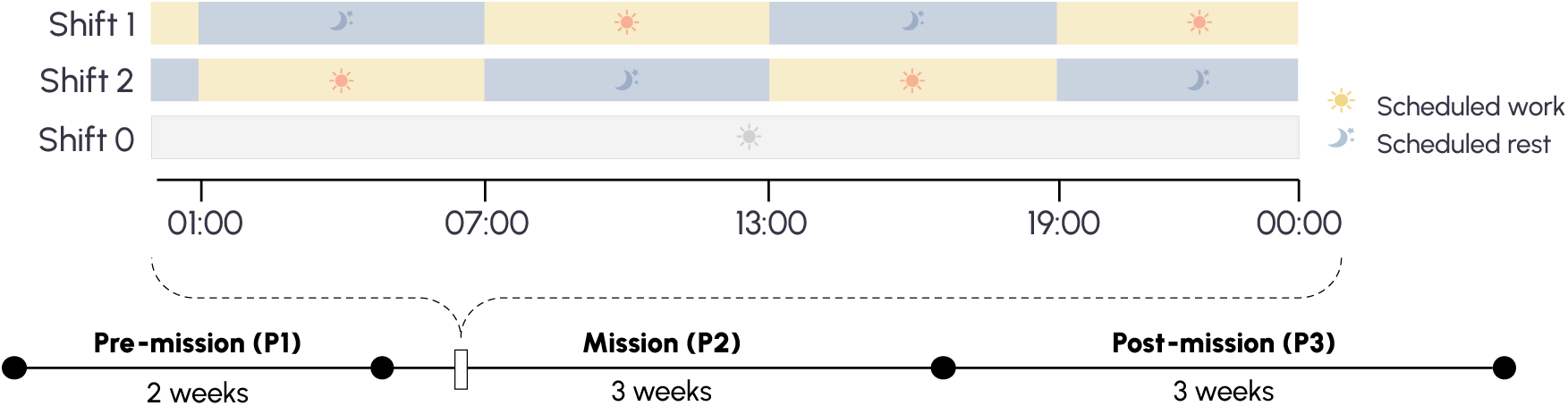
Study periods (P1, P2, and P3) and 6-hour-on/6-hour-off shift schedules (Shift 0, Shift 1, and Shift 2) during a typical submarine mission day. Submariners were organized into complementary shifts that remained fixed throughout the mission. Shift 1 rested from 01:00 to 07:00 and 13:00 to 19:00, and worked from 07:00 to 13:00 and from 19:00 to 01:00. In contrast, Shift 2 rested from 07:00 to 13:00 and from 19:00 to 01:00, and worked from 01:00 to 07:00 and from 13:00 to 19:00. Shift 0, shown in grey, represents personnel assigned to on-call or extended-hour duties. Prior to the pre-mission period (P1), submariners had no previous deployments, while the post-mission period (P3) refers to the 2–3 weeks following the mission, during which submariners resumed their normal work routines without subsequent deployments.

Pre-mission refers to the period of two weeks before the submarine mission and serves as a baseline condition, as no operational deployments occurred in the preceding months. During this period, participants kept their usual routines, including standard working hours at the naval base. Mission refers to a three-week submarine mission, in which submariners adhered to a 6-hours on/6-hours off split work schedule. To ensure continuous operations during the mission, submariners were assigned to three different shifts. Shift 1 (n = 13) operated from 07:00 to 13:00 and from 19:00 to 01:00, while shift 2 (n = 12) worked from 01:00 to 07:00, and from 13:00 to 19:00. Shift 0 (n = 5) included personnel with on-call duty or extended working hours. Each participant had the same shift assignment throughout the entire mission duration. The submarine involved in this mission is a diesel-electric type (Tridente-class), which is quite compact, and compartments for sleep, meals, leisure and work are adjacent and very small. Submarine lighting was predominantly yellow (≈560–590 nm) and consistently dim, with crew members exposed to light levels above 100 lux (*tat_100lux*) for an average of only 11 minutes during the entire mission. Finally, post-mission corresponds to a three-week recovery period, in which submariners returned to a standard working schedule and/or off-duty days.

### An actigraphy feature-screening approach based on effect size

*Constructing a panel of actigraphy features.* A panel of forty-one actigraphy features, detailed in **Table S1**, was calculated with *circStudio* (v1.0-*alpha*.1), a custom package for actigraphy data analysis built on top of *pyActigraphy* (Hammad et al., 2021, 2024). This custom version was modified for compatibility with Python 3.12+ and extended with a module for analyzing light recordings using mathematical models of circadian rhythms. Specifically, twelve features were derived from two families of light-informed mathematical models of circadian rhythms (Forger et al., 1999; Hannay et al., 2019): Kronauer-based (Forger, Jewett) and Hannay (Single Population [HannaySP], Two Populations [HannayTP]). Prior to numerical integration, missing values from off-wrist periods of the actigraphy trace were imputed by averaging the light intensity values recorded at the same time on different days, after which the signal was resampled at a 10-minute rate. For a given light intensity trajectory in *lux*, *L*(*t*), these models were integrated using the SciPy’s *odeint* function, which calls the LSODA integrator from the FORTRAN library ODEPACK.

Dim light melatonin onset (DLMO), a marker of circadian phase, was estimated by applying a seven-hour offset to the time of the core body temperature minimum estimates from the four models, consistent with prior work (Lim et al., 2025; Murray et al., 2021; Revell et al., 2006). Since previous work showed that model predictions contained larger systematic errors for individuals with circadian disruption (Huang et al., 2021), we did not rely on the nominal DLMO values predicted by the models. Instead, we focused on the predicted trajectory of DLMO values; using *scipy.stats.linregress*, we calculated the slope of the regression line across P1, P2, and P3, which reflects the kinetics of DLMO changes, and Pearson’s r, which quantifies the correlation between predicted DLMO trajectory and time. Correlations between model predictions across study periods (P1, P2, and P3) were assessed using Spearman’s rank correlation coefficient (*ρ*) as implemented in *scipy.stats.spearmanr*.

*Screening actigraphy-derived features.* The main axes of variance were identified via principal component analysis (PCA) on z-scored data (*sklearn.preprocessing.scale*) using *sklearn.decomposition.PCA*. Next, to evaluate if any actigraphy features from the panel could discriminate between at least one study period or shift, a null distribution of F-values was generated using a Monte Carlo permutation test (10^!^permutations). These F-values were subsequently converted to *Cohen’s f*, a standardized effect size metric. From this distribution, a false positive rate (FPR), also known as empirical p-value, was estimated and controlled for the false discovery rate (FDR) associated with multiple hypothesis testing via the Benjamini-Hochberg procedure. Study period labels (P1, P2, P3) were randomly permuted within each subject using the *pandas sample* method, preserving the repeated-measures structure of this factor. Shift labels (S0, S1, S2) were also randomly permuted within each study period block. To select the most informative features, a dual filtering criterion was applied based on statistical significance (FDR < 0.05) and effect size *(Cohen’s f* > 0.63). This effect size threshold corresponds to the minimum detectable effect size for a repeated measures design with two factors, given our total sample size (n = 29) and approximately 83% power, as estimated using *G*Power* (version 3.1.97).

To assess the ability of the previously selected features to discriminate across pair-wise comparisons of period and shift levels, receiver operating characteristic (ROC) curves were drawn using the *roc_curve* function, and the area under the curve (AUC) was computed with the *auc* function, both from the *scikit-learn* package. In this analysis, each actigraphy-derived feature was treated as a univariate classifier, and the AUC was used to quantify its discriminative performance across factor levels. To determine the statistical significance of the AUC observed, an AUC null distribution was constructed using a

Monte Carlo permutation test. Plots were generated using the *seaborn* and *matplotlib* libraries. All analyses were performed in Python 3.12 using the PyCharm 2024.3.3 (Professional Edition) integrated development environment (IDE). Complete code and documentation can be found in the **Supplementary Notebook** and in [*circStudio* GitHub re-pository](https://github.com/djmarques/circStudio). The overall workflow of the feature-screening approach base on effect size quan-tification is illustrated in **Figure 2**.

**Figure 2.**
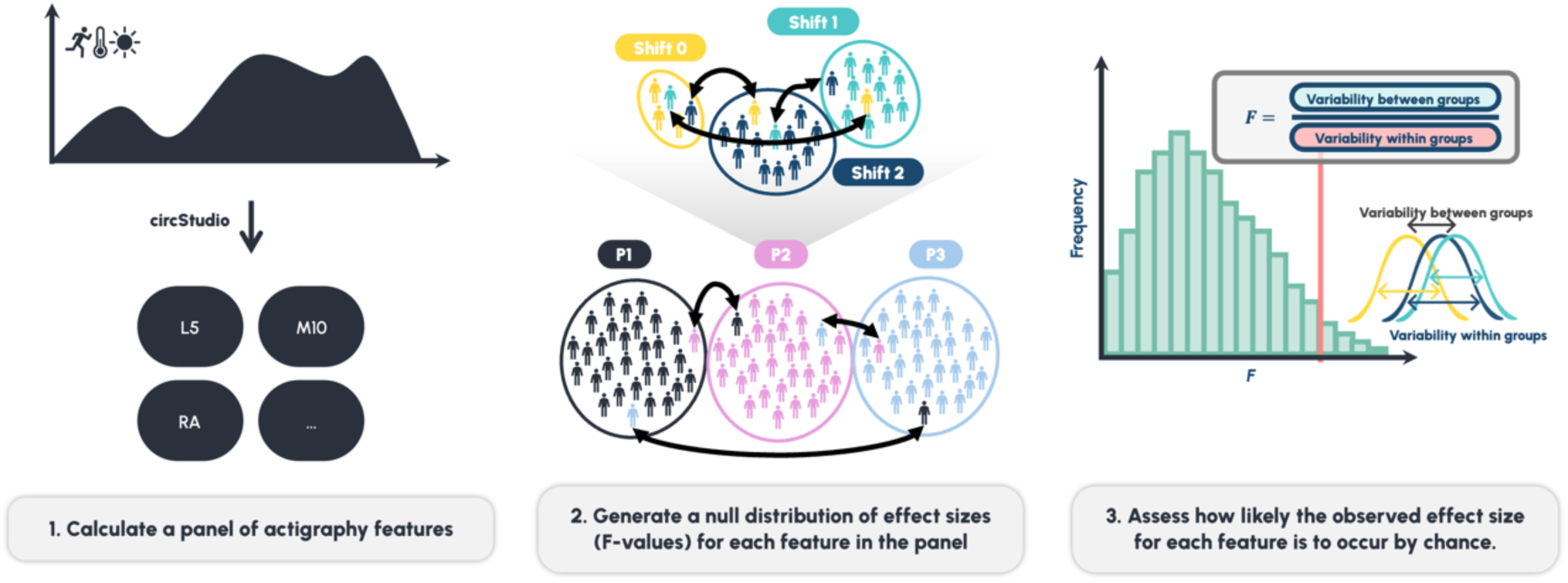
Illustration of the actigraphy feature-screening approach based on effect size quantification. First, a panel of forty-one actigraphy-derived features was computed from movement, light and skin temperature data. Next, the observed effect size was estimated using the F-value, which can be converted to a standardized effect size metric (*Cohen’s f*) by accounting for the degrees of freedom. The F-value represents the ratio of between-group to within-group (error) variability. To determine the statistical significance of the observed F-value, a null distribution of effect sizes was generated by randomly permuting shift (Shift 0, Shift 1, Shift 2) and study period (P1, P2, P3) labels across subjects. For each permutation, the F-value was recalculated, and this process was repeated multiple times to yield a distribution of null F-values. This distribution was then used to establish the probability of observing an F-value as extreme, or more extreme, than the observed value under random chance.

## Results

### Identifying candidate actigraphy submarine mission markers

To identify the dominant sources of variance in the dataset, we performed principal component analysis (PCA). Over a third of the total variance observed was concentrated within the first principal component (PC1), which distinctly separated the mission period (P2) from the pre- and post-mission periods (P1 and P3), respectively [**Figures 3A, S1]**. The two first principal components also differentiated between shifts during P2, but not in P1 nor P3 [**Figure 3B**]. The top ten contributors to PC1, ranked by loading coefficient, included the Pearson’s *r* values of all four mathematical models (*forger_r, jewett_r, hannaytp_r, hannaysp_r*), the slope of the Jewett model (*jewett_slope*), the time spent per day above 100 and 500 lux (*tat_100lux, tat_500lux*), the light intensity during the ten most illuminated hours of the day (*M10_light*), and the relative amplitude (*RA*), a metric that measures the contrast between periods of highest (*M10*) and lowest activity (*L5*) hours **[Supplementary Notebook 1]**. Overall, this seems to indicate that the primary axis of variation in the dataset reflects both the kinetics (captured by the slope) and the strength and direction of temporal shifts in the predicted daily DLMO (captured by Pearson’s *r*), alongside with light intensity metrics and *RA*.

**Figure 3.**
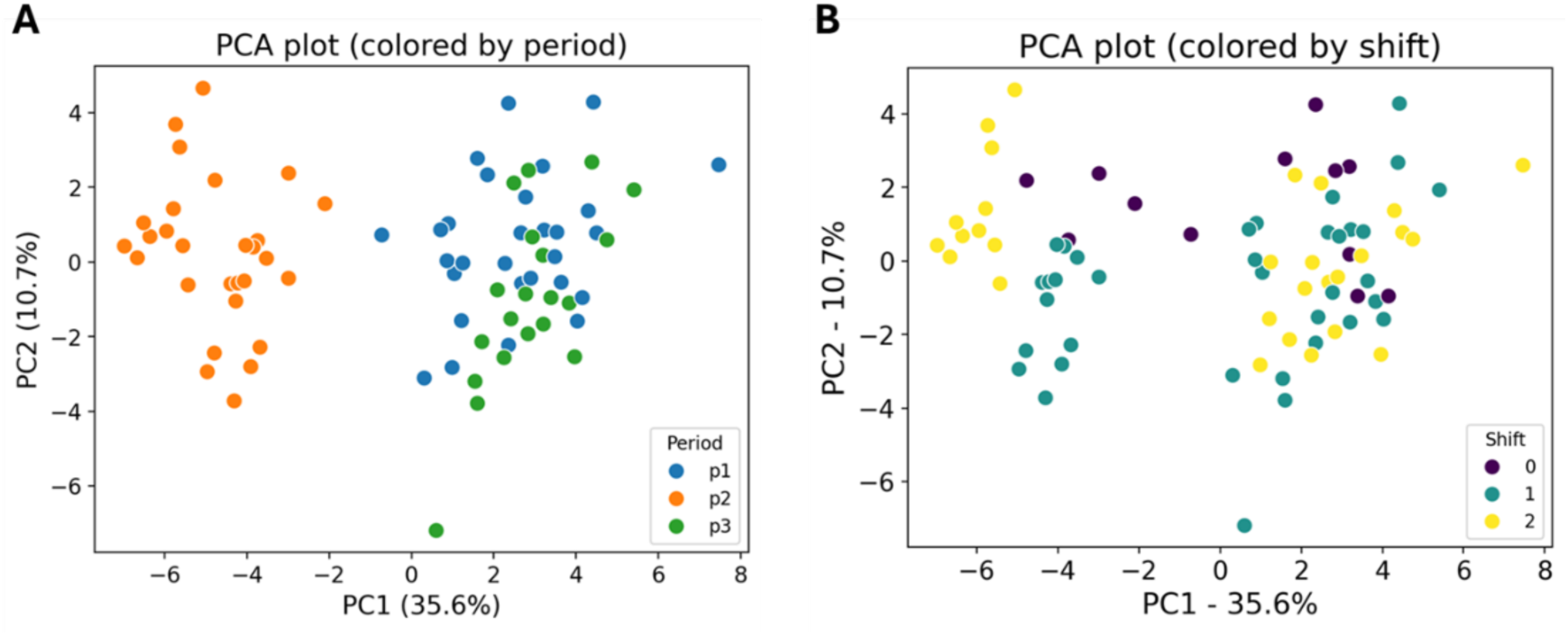
Principal component analysis of actigraphy features. (A) PCA plot displaying the first two principal components (PC1 and PC2), with samples colored by study period. PC1 effectively separates P2 (the mission period) from P1 and P3 (pre- and post-mission periods, respectively), indicating that study period is a major source of variance. (B) The same PCA projection with samples colored by shift, showing a clear separation of shifts in P2, but not in P1 and P3.

Given these observations, we sought to identify which features provided the greatest discriminatory ability across study periods. Features were initially filtered based on a Monte Carlo permutation approach to estimate the *Cohen’s f*, as an effect size metric, and the FPR, which was subsequently adjusted for multiple comparisons via Benjamini–Hochberg FDR procedure. Among the five highest-ranking features by effect size were several variables mentioned before, with *tat_100lux* having the highest effect size (*Cohen’s f* =1.91, FDR ≤ 1.64× 10^-4^), followed by *forger_r* (*Cohen’s f* = 1.69, FDR ≤ 1.64 × 10^-4^), *tat_500lux* (*Cohen’s f* = 1.66, FDR ≤ 1.64 × 10^-4^), *jewett_r* (Cohen’s f = 1.62, FDR ≤ 1.64 × 10^-4^), and *hannaytp_r* (Cohen’s f = 1.38, FDR ≤ 1.64 × 10^-4^) [**Figure 4A, Table S2]**. Similarly, *hannaysp_r* (Cohen’s f = 1.12, FDR ≤ 1.64 × 10^-4^) exhibited comparable discriminative power across at least one study period. Taken together, these findings point to a high degree of agreement among the four mathematical models, supporting the notion that the stability and/or temporal dynamics of the predicted DLMO trajectory significantly differed in at least one study period.

**Figure 4.**
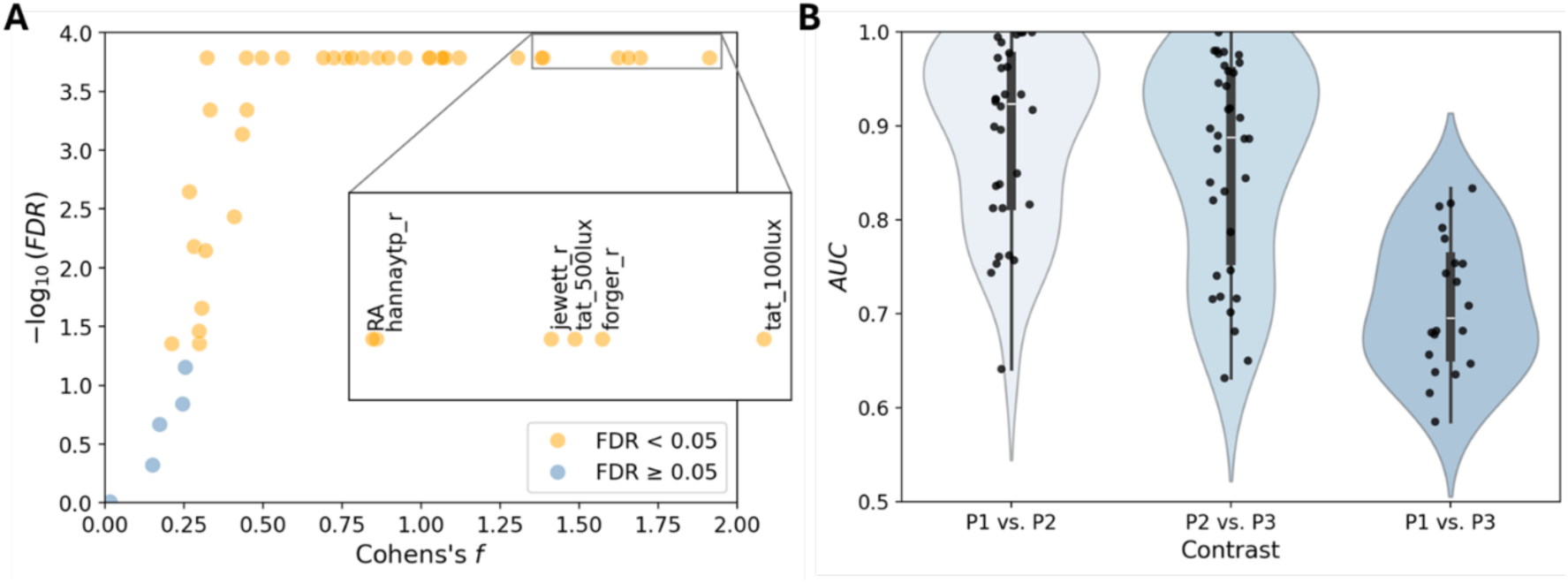
Ability of features to discriminate across study periods. (A) Scatter plot of module of effect size (*Cohen’s f* – X-axis) and significance (−log_10_(FDR), Y-axis) of features with potential to distinguish between study periods. The five features with the largest effect sizes are shown in the inset. Light-related variables ranked the highest, alongside with the Pearson’s *r* for several mathematical models. (B) Jittered violin plots of AUC values across features for each pairwise contrast. Discrimination was strongest when comparing the mission period (P2) to either pre-(P1) or post-mission (P3), while pre-(P1) vs. post-mission (P3) periods were markedly less distinguishable, indicating a transient effect of submarine confinement.

To determine whether specific study period contrasts were associated with greater discriminability across features, we examined the AUC distributions derived from ROC curves for all pairwise comparisons (P1 vs. P2, P2 vs. P3, and P1 vs. P3) [**Figure 4B**]. The AUC distribution for the pre-vs. post-mission contrast (P1 vs. P3) differed significantly from the pre-vs. mission (P1 vs. P2*; p = 7.53 ×10^-7^, AUC=0.85*) and mission vs. post-mission (P2 vs. P3; *p = 4.98 ×10^-5^, AUC = 0.7*) contrasts. Yet, no significant difference was observed between the P1 vs. P2 and P2 vs. P3 distributions (*p = 0.22, AUC=0.58*). These results point to greater similarity between the pre- and post-mission periods, with a higher concentration of discriminative features in comparisons involving the mission period.

Building on these findings, we used the AUC to select features with the strongest ability to discriminate between pairs of study periods **[Table S3]**. Features meeting the criteria of FDR < 0.05 and AUC ≥ 0.99 in at least one contrast were deemed highly discriminative and selected; their mean and standard deviation are shown in **Table 1** and include both light-related metrics and the Pearson’s *r* for the Forger and Jewett models [**Figure 5A**]. Summary statistics for additional features are provided in **Table S4**. Illustrative ROC curves and violin plots for *jewett_r* are shown in **Figure 5B,C**, with the remaining plots available in **Figures S3 and S5**.

**Figure 5.**
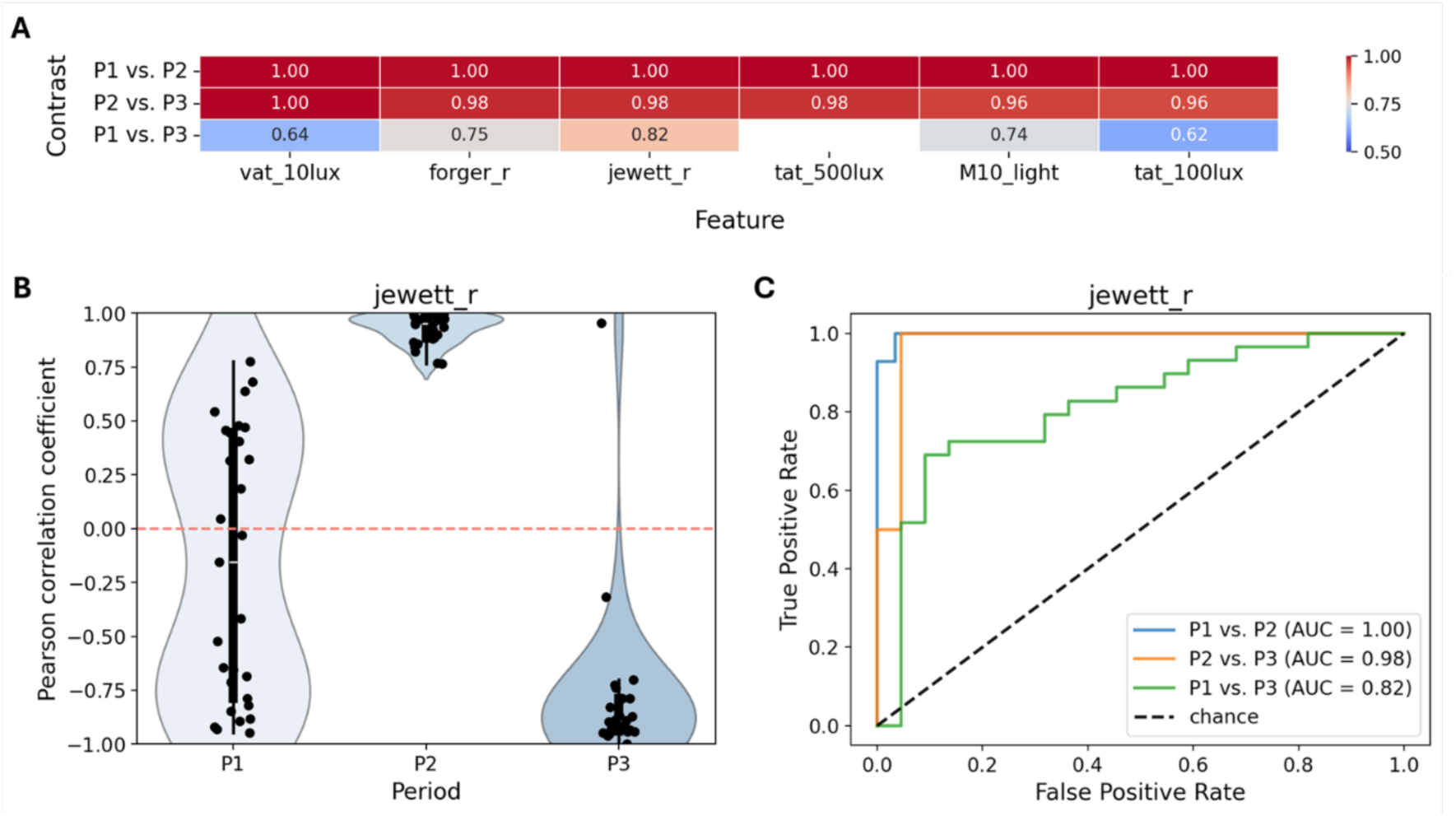
Pairwise feature discrimination across study periods. (A) Heatmap showing the performance of features in pairwise period comparisons, ranked by average AUC. (B) Illustrative jittered violin plots and (C) ROC curves for *jewett_r* (Pearson’s correlation coefficient between each day in the recording and the respective DLMO predicted by the Jewett model), showing a very strong ability to distinguish P2 from P1 and P3, but not as strong between P1 and P3. Each point in panel B represents an individual submariner. Each point in B represents an individual submariner.

**Table 1.** Mean and standard deviation in each study period of features with FPR < 0.05 AUC ≥ 0.99 in at least one pairwise comparison between study periods. Variable descriptions: *vat_10lux* – median daily light exposure above 10 lux; *forger r, jewett_r* – Pearson’s correlation coefficient between each day in the recording and the corre-sponding DLMO predicted by the Forger and Jewett model, respectively; *tat_500lux, tat_100lux* – median daily light exposure time above 500 and 100 lux, respectively; *M10_light* – mean light intensity during the ten brightest hours of the day.

| Feature | P1 (mean ± SD) | P2 (mean ± SD) | P3 (mean ± SD) |
| --- | --- | --- | --- |
| *vat_10lux* (lux) | 115.17 ± 58.52 | 27.17 ± 4.01 | 90.32 ± 26.25 |
| *forger_r* | -0.36 ± 0.64 | 0.95 ± 0.04 | -0.81 ± 0.41 |
| *jewett_r* | -0.18 ± 0.62 | 0.93 ± 0.07 | -0.77 ± 0.41 |
| *tat_500lux* (minutes) | 156.1 ± 51.96 | 0.57 ± 1.93 | 133.54 ± 55.28 |
| *M10_light* (lux) | 1600.58 ± 490.09 | 264.44 ± 177.83 | 1219.72 ± 605.44 |
| *tat_100lux* (minutes) | 284.17 ± 76.28 | 11.11 ± 14.33 | 241.17 ± 85.01 |

During the submarine mission, light-related metrics providing the greatest discrimination between shifts consistently exhibited low intensity values, and reduced time, per day, above 100 and 500 lux (*tat_100lux*: 11.11 ± 14.33 min; *tat_500lux*: 0.57 ± 1.93), with low light intensity values during the ten brightest hours of the day (*M10_light*: 264.44 ± 177.83). Moreover, in P1, *forger_r* and *jewett_r* showed minimal correlation of the predicted daily DLMO with time (−0.36 ± 0.64 and −0.18 ± 0.62, respectively). However, during P2, both coefficients increased markedly (0.95 ± 0.04 and 0.93 ± 0.07), indicating a strong positive association consistent with a phase delay. In contrast, P3 was characterized by strongly negative correlations (−0.81 ± 0.41 and −0.77 ± 0.41), suggestive of a phase advance. A similar pattern was followed by the daily DLMO predicted by HannaySP model (P1: −0.24 ± 0.60; P2: 0.80 ± 0.46; P3: −0.56 ± 0.50) and HannayTP model (P1: −0.24 ± 0.61; P2: 0.90 ± 0.37; P3: −0.70 ± 0.45) [**Figures 6, S7–9]**. Taken together, all four models seem to consistently predict a pronounced phase delay in response to the dim-light conditions of the submarine, followed by a phase advance in the post-mission period.

**Figure 6.**
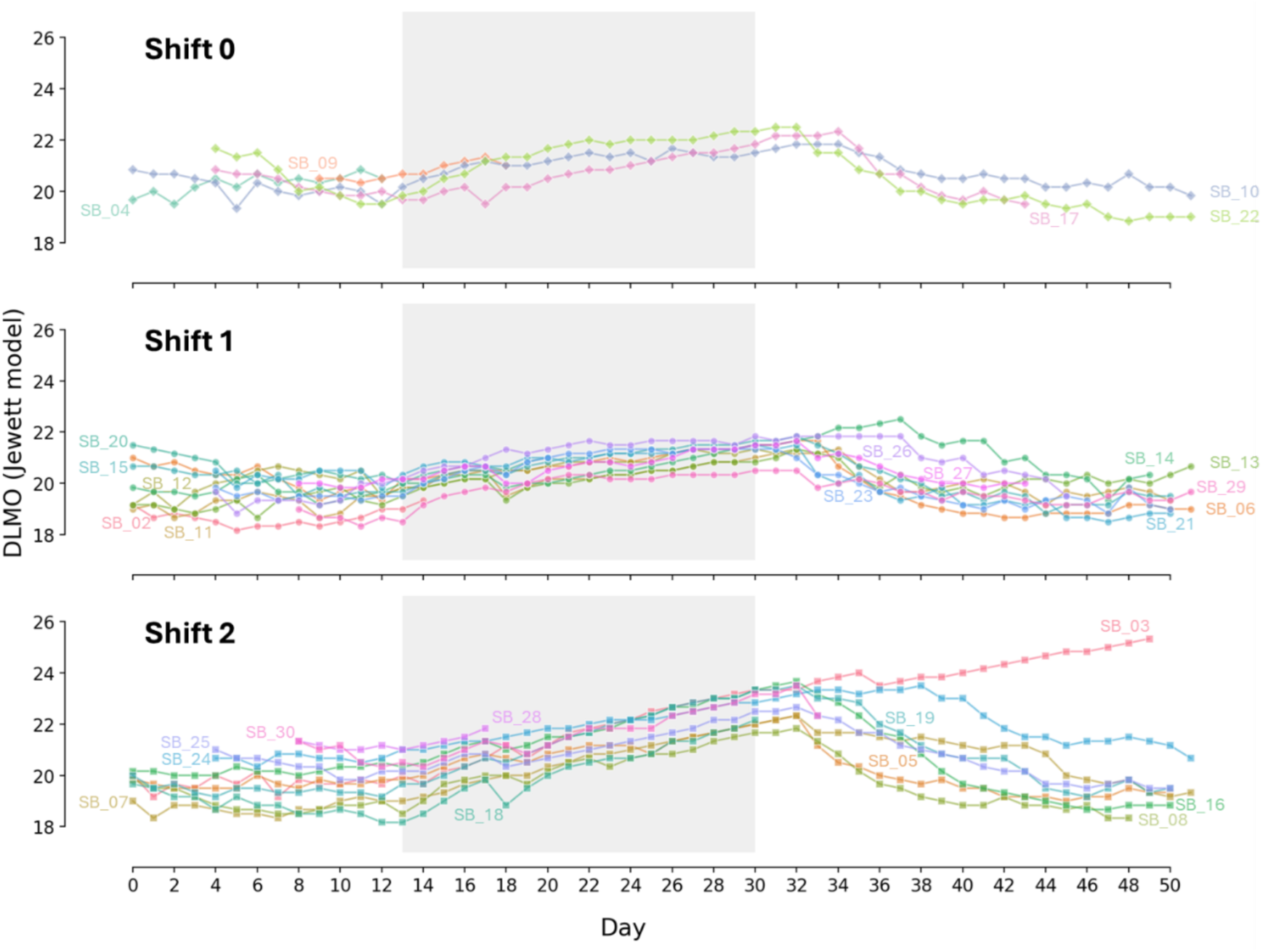
Predicted daily circadian phase trajectory based on the Jewett model. Predicted dim light melatonin onset (DLMO) for each individual across the whole study, separated by shifts. The submarine mission period (P2) is shaded in light grey.

We also examined whether individuals with earlier or later predicted DLMOs retained this pattern during the mission and whether recovery depended on the predicted pre-mission DLMOs. A strong, statistically significant correlation was observed between predicted DLMOs in P1 and P2, indicating that individuals with later predicted DLMOs before the mission also exhibited later predicted DLMOs during the submarine mission *Spearman’s ρ =0.678, p < 1.00 × 10^-4^*) [**Figure 7**]. However, no significant correlations were found between predicted DLMOs in P2 and P3 (*Spearman’s ρ =0.181, p = 0.420*) or between P1 and P3 (*Spearman’s ρ =0.398, p < 0.0665*) [**Figure 7**], indicating that recovery was not dependent on the predicted pre-mission DLMOs and that the pattern observed in P1 was not fully restored during P3.

**Figure 7.**
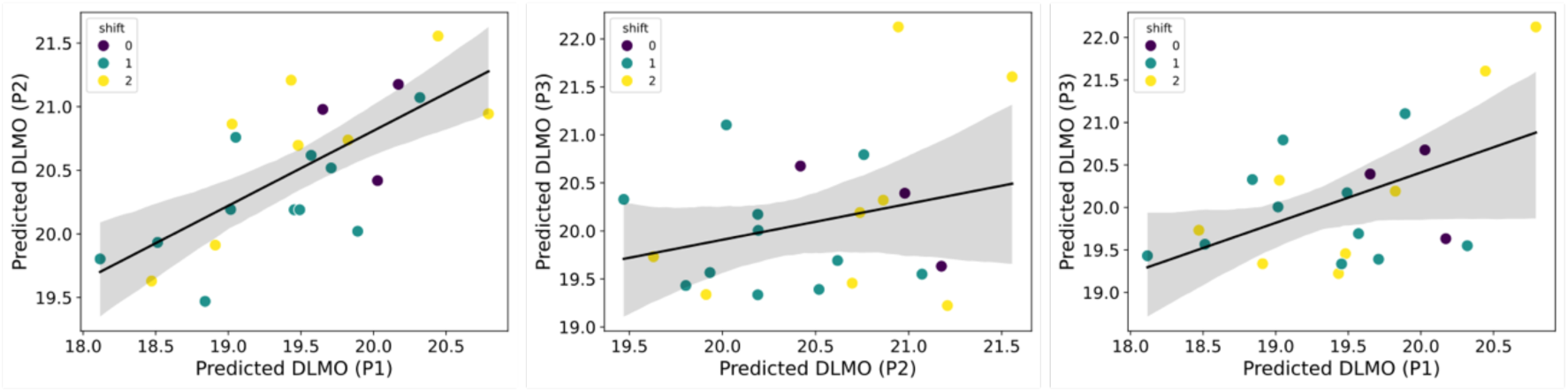
Correlation between predicted DLMOs obtained from four mathematical models across study periods. A statistically significant correlation was observed between predicted DLMOs in the pre-mission and mission periods (P1 vs. P2: Spearman’s *ρ =0.678, p <1.00 × 10^-4^*), but not between the mission and post-mission (P2 vs. P3: Spearman’s *ρ =0.181, p = 0.420*) or pre-mission and post-mission (P1 vs. P3: Spearman’s *ρ =0.398, p < 0.0665*) periods.

### Identifying shift-specific actigraphy markers during the submarine mission

While submarine missions represent a strong circadian perturbation, we investi-gated whether their physiological impact might also vary depending on shift assignment. If so, we would expect distinct actigraphy-based signatures to emerge across shifts.

Although the distribution of effect sizes was significantly lower when grouping by shift rather than by study period (*Mann Whitney’s U = 3747.00, p = 3.00×10^-6^*) [**Figure 8A**], we wondered if it would be possible to identify features capable of distinguishing between shifts. In this context, P1 served as a negative control, as participants had not yet been assigned to shifts, while P3 provided insights into post-mission recovery upon return to baseline conditions. No features differentiated shifts at a 5% significance level in either P1 or P3. However, during the mission period (P2), seventeen features met the threshold for statistical significance and were ranked by effect size (*Cohen’s f*) [**Figure 8B**]. The top five features were related to sleep, namely the *main_sleep_bout* (*Cohen’s f* = 1.59, FDR ≤ 8.19 *×* 10^-4^), *sleep_midpoint* (*Cohen’s f* =1.41, FDR ≤ 8.19 *×* 10^-4^) and the sleep regularity index (*SRI*) (*Cohen’s f* = 1.27, FDR ≤ 8.19 *×* 10^-4^). The highest effect size was observed for the *RA* (*Cohen’s f* = 1.68, FDR ≤ 8.19 *×* 10^-4^), and the lowest among the top 5 was observed in *exp_level* (*Cohen’s f* = 1.06; FDR = 0.0023).

**Figure 8.**
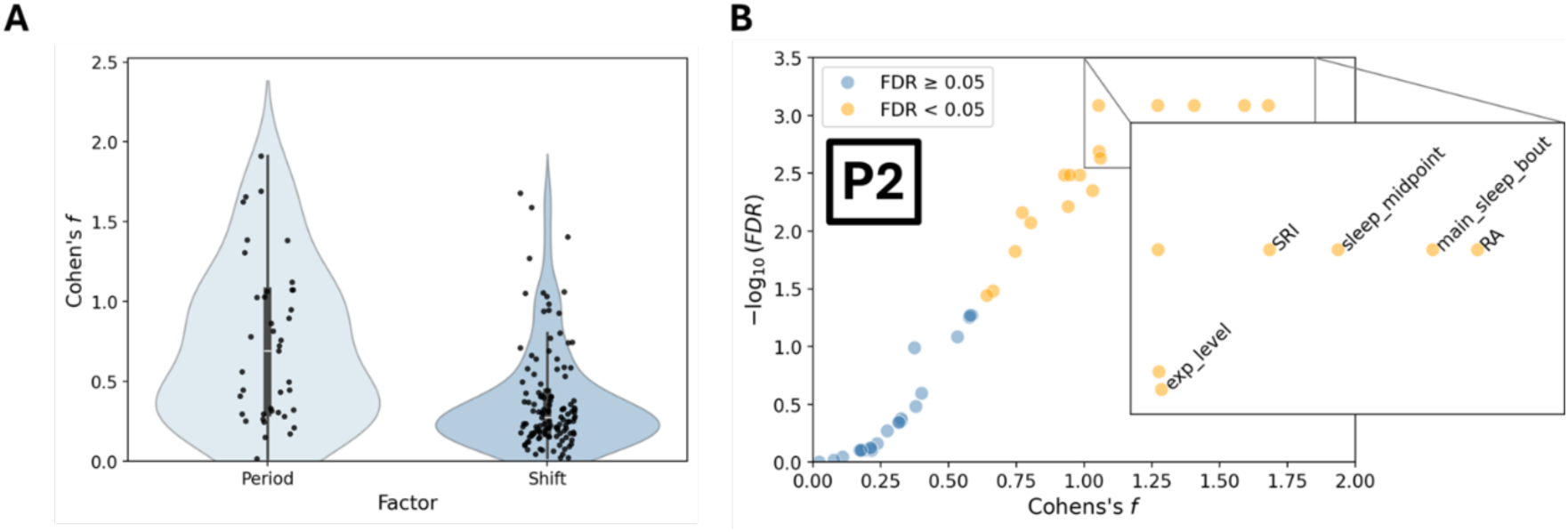
Discriminative power of features across shifts during the mission period (P2) (A) Jittered violin plots of *Cohen’s f* distributions across features, by period and shift, showing that larger effect sizes are mainly associated with distinctions between study periods, while shift comparisons within P2 are characterized by smaller, yet non-negligible, effect sizes. (B) Scatter plot of module of effect size (*Cohen’s f* – X axis) and significance (−log10(FDR), Y-axis) of features that significantly distinguish at least one shift in P2. The inset on the right highlights the top five features ranked by effect size (*Cohen’s f*).

Variables derived from mathematical models of circadian rhythms were absent from the set of features that differentiated between at least one shift. While *forger_slope* yielded a statistically significant result (*Cohen’s f* = 0.93, FDR = 0.0033), its effect size was comparatively modest and it was not replicated across other models (*jewett_slope*:

*Cohen’s f* = 0.58, FDR = 0.054, *hannaysp_slope*: *Cohen’s f* = 0.17, FDR = 0.79; *han-naytp_slope*: *Cohen’s f* = 0.20, FDR = 0.53). Similarly, a small yet significant effect was observed for *jewett_r* (*Cohen’s f* = 0.75, FDR = 0.015), but not for the Pearson’s *r* of the other models (*forger_slope*: *Cohen’s f* = 0.53, FDR = 0.082; *hannaysp_r*: *Cohen’s f* = 0.18, FDR = 0.79; *hannaytp_r*: *Cohen’s f* = 0.21, FDR = 0.79). Thus, although all models consistently predicted a phase delay during P2 when contrasting study periods, the absence of cross-model agreement when grouping by shift leads us to conclude that there is insufficient evidence that either the linear dependence of the predicted DLMO on time or the rate of change in predicted DLMO with respect to time varied across shifts.

The discriminative capacity of the seventeen features was further evaluated across pairwise shift comparisons in P2 (S0 vs. S1, S0 vs. S2, S1 vs. S2) **[Table S3]**. As previously, discrimination ability was measured using ROC curves, with the AUC serving as a performance metric. Shift-discriminating features meeting the criteria of an FDR < 0.05 and AUC ≥ 0.99 in at least one comparison are presented in **Figure 9**, with the mean and standard deviation shown in **Table 2**, and the remaining descriptives on **Table S5**. Illustrative ROC curves and violin plots are presented for *SRI* in **Figure9B,C**, with the other violin plots and ROC curves shown in **Figures S4 and S6**. Notably, two sleep features, *SRI* and *main_sleep_bout*, were significant across all shift comparisons in P2. Interestingly, *L5* and *RA* were found to exclusively differentiate between shift 1 .and 2, with a lower *RA* and higher *L5* in shift 2 (*L5*: 1159.07 ± 428.97 [PIM]; *RA*: 45% ± 4%) in comparison with shift 1 (*L5*: 472.22 ± 116.59 [PIM]; *RA*: 71% ± 6%), suggestive of increased nighttime movement during the five hours of lower activity.

**Figure 9.**
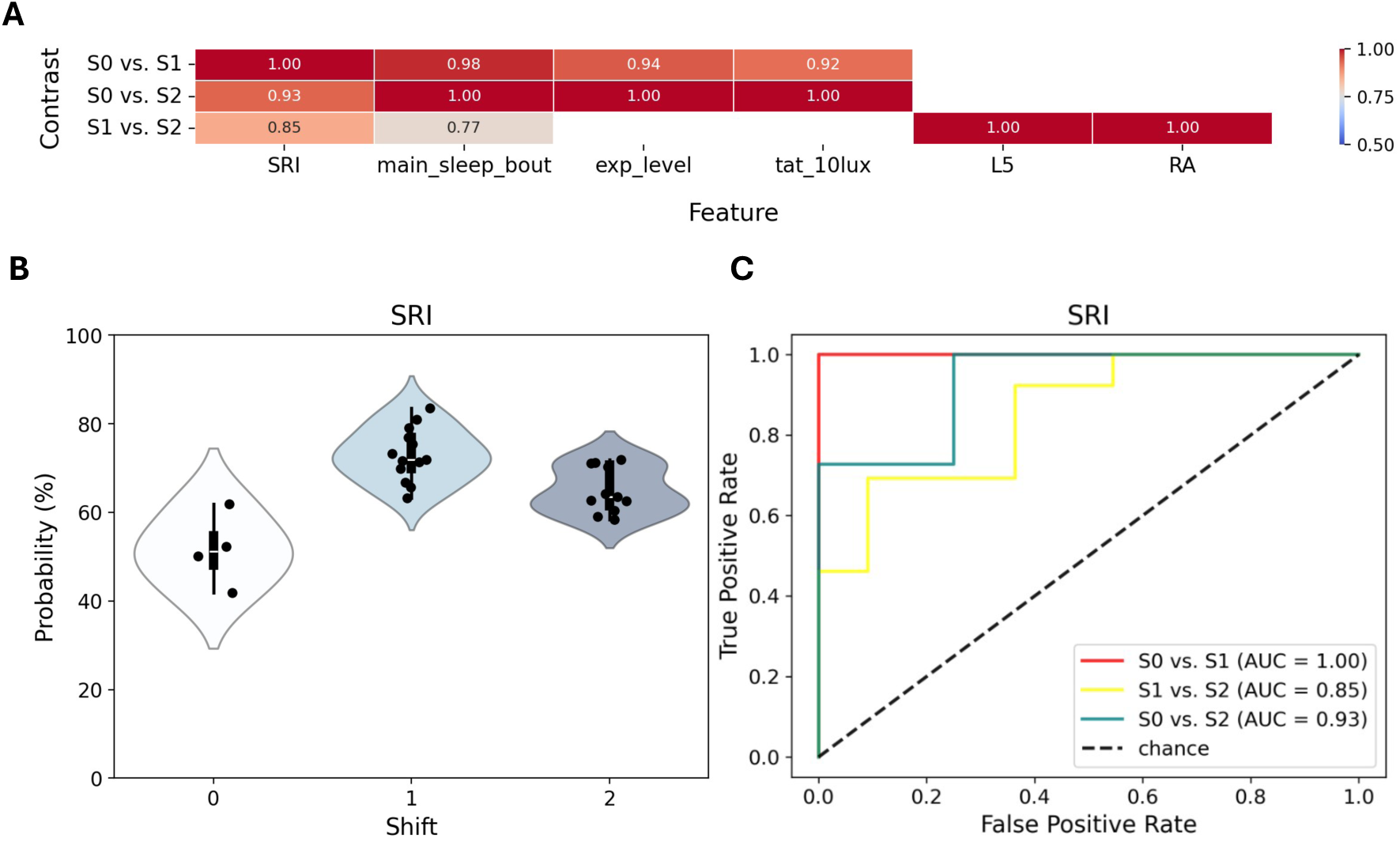
Pairwise feature discrimination across shifts. (A) Heatmap of pairwise shift comparisons in P2, with fea-tures ordered by average AUC across the three shift-pair contrasts. Empty cells denote non-significant (FDR > 0.05) results. (B) Jittered violin plots and (C) ROC curves for *SRI* (sleep regularity index). Most features exhibiting near-perfect (AUC > 0.99) discriminative performance differentiated between the split shift schedules (shifts 1 and 2) and the on-call schedule (shift 0), with *RA* and *L5* being notable exceptions. Each point in B represents an individual sub-mariner.

**Table 2.** Mean and standard deviation in each shift of features with FDR < 0.05 and AUC ≥ 0.99 in at least one pairwise comparison between shifts. Variable descriptions: *SRI* – probability that any two time points exactly 24 hours apart are in the same state (either sleep or wake), averaged across all days; *main_sleep_bout* – duration of the primary sleep episode each day, defined as the longest consolidated period of sleep within a 24-hour interval; *exp_level* – median light exposure level; *L5* – mean light intensity during the five least active hours of the day; *RA* – relative amplitude between the ten most active hours (M10) and the five least active hours of the day (L5).

| Feature | Shift 0 (mean ± SD) | Shift 1 (mean ± SD) | Shift 2 (mean ± SD) |
| --- | --- | --- | --- |
| SRI (%) | 51.55 ± 8.26 | 73.03 ± 6.03 | 65.02 ± 5.16 |
| main_sleep_bout (minutes) | 377.62 ± 52.23 | 259.15 ± 29.99 | 299.91 ± 25.63 |
| exp_level (lux) | 4.13 ± 3.27 | 0.70 ± 0.63 | 0.53 ± 0.32 |
| tat_10lux (minutes) | 527.75 ± 108.62 | 309.15 ± 93.77 | 292.91 ± 74.81 |
| L5 (PIM) | 846.84 ± 420.88 | 472.22 ± 116.59 | 1159.07 ± 428.97 |
| RA (%) | 60 ± 17 | 71 ± 6 | 45 ± 4 |

## Discussion

This study aimed to identify actigraphy signatures associated with living aboard a submarine. The rationale for comparing signatures between different shifts stemmed from a considerable body of literature linking night shift work with a wide spectrum of detrimental health outcomes (Dutheil et al., 2020; Hansen & Pedersen, 2024; Jahn et al., 2024; Manohar et al., 2017; Moon et al., 2024; Wang et al., 2014). However, some meta-analyses emphasized the need for well-designed and adequately powered studies using standardized methods to facilitate comparisons (Jahn et al., 2024; Viramgami et al., 2025). The dual filtering criterion adopted in this study addresses this concern by retaining only those actigraphy features with an effect size above *Cohen’s f* = 0.63 **[see Methods]**. By moving beyond *p*-value-based analyses, our results enable future submarine studies to estimate the sample sizes needed to detect meaningful effects. One surprising finding in our study was that the detected effects were large. This may be explained by the submarine environment itself, which constitutes an exceptionally strong and uniform perturbation. The combination of its disruptive nature and the homogeneity of conditions likely amplify the detectability of biological effects, allowing them to be observed even with a relatively small sample size (n = 29).

As expected, light-related variables emerged as predominant sources of variance in our dataset. Interestingly, the Pearson’s *r*, which measured the correlation between the predicted DLMO trajectory and time, was effective in distinguishing the mission period (P2) from the pre- and post-mission periods (P1 and P3, respectively). Remarkably, no conclusive evidence was found of a shift-specific impact on either the Pearson’s *r* or the slope of the regression line fitted to the DLMO model predictions, seemingly contradicting a previous study on a nuclear submarine in which an adaptation to the watchkeeping schedule seems to occur (Chabal et al., 2024). Nonetheless, the absence of shift-related differences in the predicted DLMO is consistent with results from a prior study involving submariners also on a 6-hours-on/6-hours-off routine in a diesel-electric submarine, where melatonin secretion failed to adjust to the biphasic work-sleep cycle (Van Puyvelde et al., 2022).

Remarkably, model-predicted DLMOs showed a significant correlation between the pre-mission (P1) and mission (P2) periods, suggesting that adaptation to the submarine environment may have followed a phase-dependent (or chronotype-specific) pattern not maintained upon return to baseline conditions in the post-mission (P3) period. Notably, no correlation was observed between P1 and P3, indicating that submariners did not revert to the circadian phase pattern seen before the mission. This lack of correlation may reflect either residual circadian disruption – where the post-mission period (P3) was insufficient for complete resynchronization – or interindividual variability in re-entrainment, with some participants adjusting more rapidly to external cues such as natural light exposure and work routines than others.

Another interesting pattern observed in our study was that individuals assigned to split shift schedules differed consistently from those assigned to on-call/extended hours, that is, increased similarity between shifts 1 and 2, and a higher dissimilarity between these two and shift 0. A noteworthy exception to this pattern was the reduction in *RA* and an elevation in *L5* observed in shift 2, with a working bout entirely during the biological night. This signature likely reflects increased nocturnal activity and, thus, points to impaired sleep quality and/or increased circadian disruption. Future studies in diesel-electric submariners should attempt to include some modality of portable polysomnography, as it would give further insights into the differential impact of being assigned to a shift during the biological night, in the absence of a robust photic *zeitgeber*.

As noted in the **Methods** section, we did not rely on nominal DLMO values, since previous research has reported larger systematic errors in DLMO predictions by light-informed models in individuals with circadian disruption (Huang et al., 2021). Instead, we focused on the temporal trajectory of the predicted DLMO. This leads us to introduce the notion of *metaphorical model*, *M*(*t*), a mathematical model whose predictions may not exactly match empirical measurements (e.g., empirical DLMO values) due to systematic error, but that can be useful insofar as it reliably differentiates between experimental conditions or factor levels. Conceptually, this can be expressed as *M*(*t*) = *f*(*t*) + *e*(*t*), where *f*(*t*) represents the true underlying biological process and *e*(*t*) denotes a systematic error term. If the error is approximately constant (*e*(*t*) = *c*), then the derivative of *M*(*t*) simplifies to the derivative of *f*(*t*), since the derivative of a constant term evaluates to zero: *M*’(t) = *f*’(t). In our case, across all four light-informed models, a consistent pattern of variation was predicted, namely a robust phase delay during the mission, followed by a phase advance upon return to baseline conditions in the post-mission (P3) period [**Figures 6, S7–9]**. Thus, by analyzing the pattern of variation (i.e., the derivative of the predicted DLMO) rather than the absolute DLMO values, we can circumvent the limitations introduced by larger systematic errors in model predictions and hypothesize that inferential insights can be derived even when empirical validation of the model is not feasible.

Ultimately, these findings are consistent with established literature showing that night shift workers experience poor sleep due to misalignment between the sleep-wake cycle and circadian phase (Costa, 2015). Remarkably, this disruptive pattern was detect-able in shift 2, the one most closely resembling a night shift, despite all submariners being exposed to the same environmental conditions. These results indicate that, even in the absence of natural light-dark cues, working during the biological night may contribute to sleep impairment or, more likely, that sleeping during the biological day leads to a more disrupted sleep, as observed in nurses (Martin et al., 2006). Given the parallels between submarine and night shift work, with the latter being linked to increased cardiometabolic risk and certain cancers (Su et al., 2021), it would be relevant to investigate whether mo-lecular alterations are observed in submariners and if we can observe an overlap with those documented in night shift workers.

A major limitation of our data was the small representation of female participants, reflecting the reality in most submarine crews, but leaving open the question of whether the impact of submarine and submarine-like environments would be comparable in women, as several studies point to sex-related differences in circadian rhythms (Walton et al., 2022). This study was also limited by the fact that only data from a single submarine mission was available. Since the post-mission recovery period has scarcely been evalu-ated and the importance of ensuring adequate recuperation between deployments, future studies should attempt to evaluate the long-term physiological effects of repeated subma-rine missions. Similarly, it would be valuable to investigate the differential impacts of shorter versus longer deployments, as well as the influence of vessel type, particularly in relation to the degree of spatial confinement.

While some results may align with prior expectations, they demonstrate that, even under technically challenging conditions for actigraphy data collection, the combination of restricted movement and dim light constitutes a robust enough stimulus to be reliably detected through effect size quantification. Indeed, despite the technical challenges of collecting actigraphy data under dim light and restricted movement, these very conditions may enhance detectability due to the strength and consistency of the stimulus, yielding measurable effect sizes. The ability to extract biologically meaningful signals from such constrained settings highlights the potential applicability of this methodology to other operational or clinical contexts characterized by limited mobility or light exposure, such as in bedridden individuals, older adults, and patients with dementia (Tolea et al., 2016) or depression (Elkjær et al., 2022).

## Supporting information

Supplementary Notebook

## Notes

### Competing Interest Statement

The authors have declared no competing interest.

### Summary of Updates

We revised the manuscript to emphasize the methodological contribution introduced here, namely the application of an actigraphy feature-screening approach based on effect size quantification.

### Funding

- Ministery of Defense, Portugal: P° 55/2022(17) N° 2468 / CG

