## Supplementary Figures for "An effect size-based screening approach to identify actigraphy markers of the submarine environment"

### Circadian disruption beneath the surface: a screening approach to identify actigraphy-based markers of split-shift work in submariners

Daniel Marques, Carina Fernandes, Nuno L. Barbosa-Morais, Cátia Reis

**Figure S4 Jittered violin plots for features that differentiated at least one shift during the submarine mission (P2).** Feature selection was initially based on a dual filtering criterion: statistical significance (FPR < 0.05) and effect size (Cohen's F > 0.63), the latter corresponding to the estimated minimum detectable effect size given the sample size used in this study (n = 29). Detailed descriptions of each feature are provided in Table S1. ... 6

**Figure S5 ROC curves for features that differentiated at least one study period.** Pairwise discriminative performance was assessed via AUC, with significance determined through Monte Carlo permutation testing. This post-hoc evaluation was conducted only for features passing the initial screening criteria (FPR < 0.05 and Cohen's F > 0.63). Detailed descriptions of each feature are provided in Table S1. ..................... 7

**Figure S6 ROC curves for features that differentiated at least one shift during the submarine mission (P2).** Pairwise discriminative performance was assessed via AUC, with significance determined through Monte Carlo permutation testing. This post-hoc evaluation was conducted only for features passing the initial screening criteria (FPR <

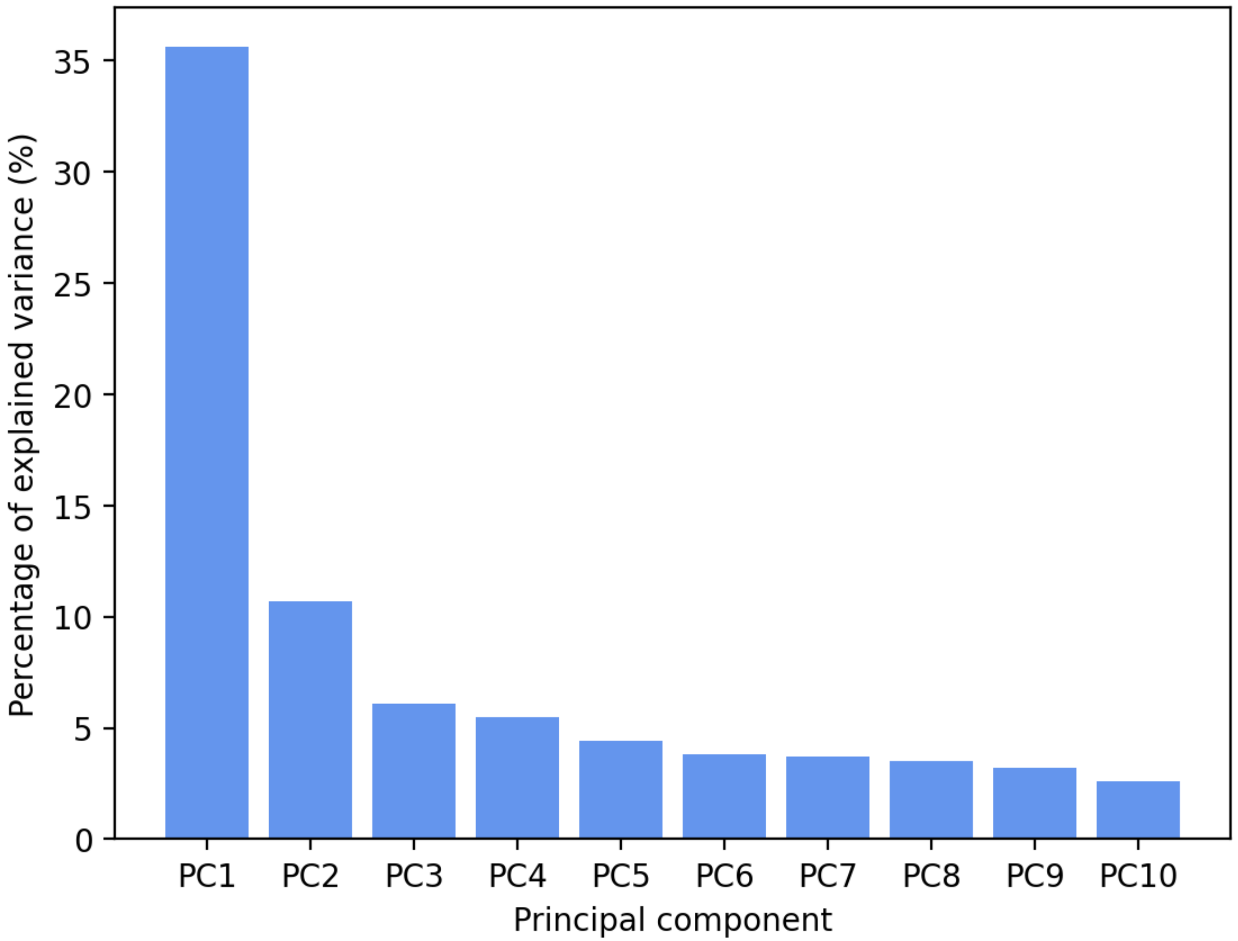

**Figure S1 Scree plot displaying the percentage of variance explained by the first ten principal components.** An important proportion of variance in the submarine dataset was captured by PC1 (35.6%) and PC2 (10.7%), with substantially lower contributions from subsequent components.

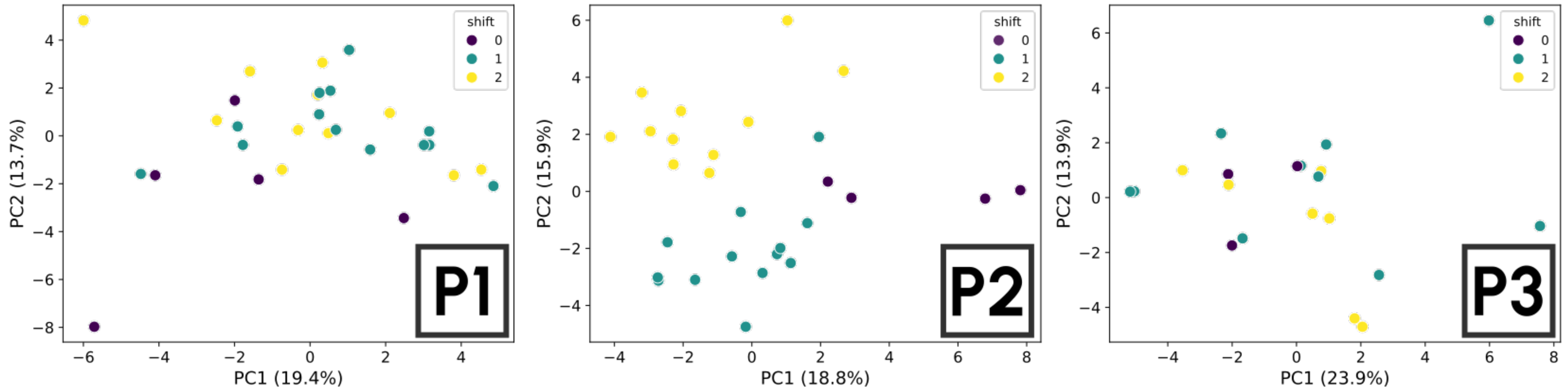

**Figure S2 Principal component analysis (PCA) stratifying by study period.** PC1 and PC2 clearly separate shifts during the submarine mission (P2), but not during the pre- (P1) and post-mission (P3) periods, indicating that any mission-related effects associated with specific shift assignments dissipated during the post-mission recovery period.

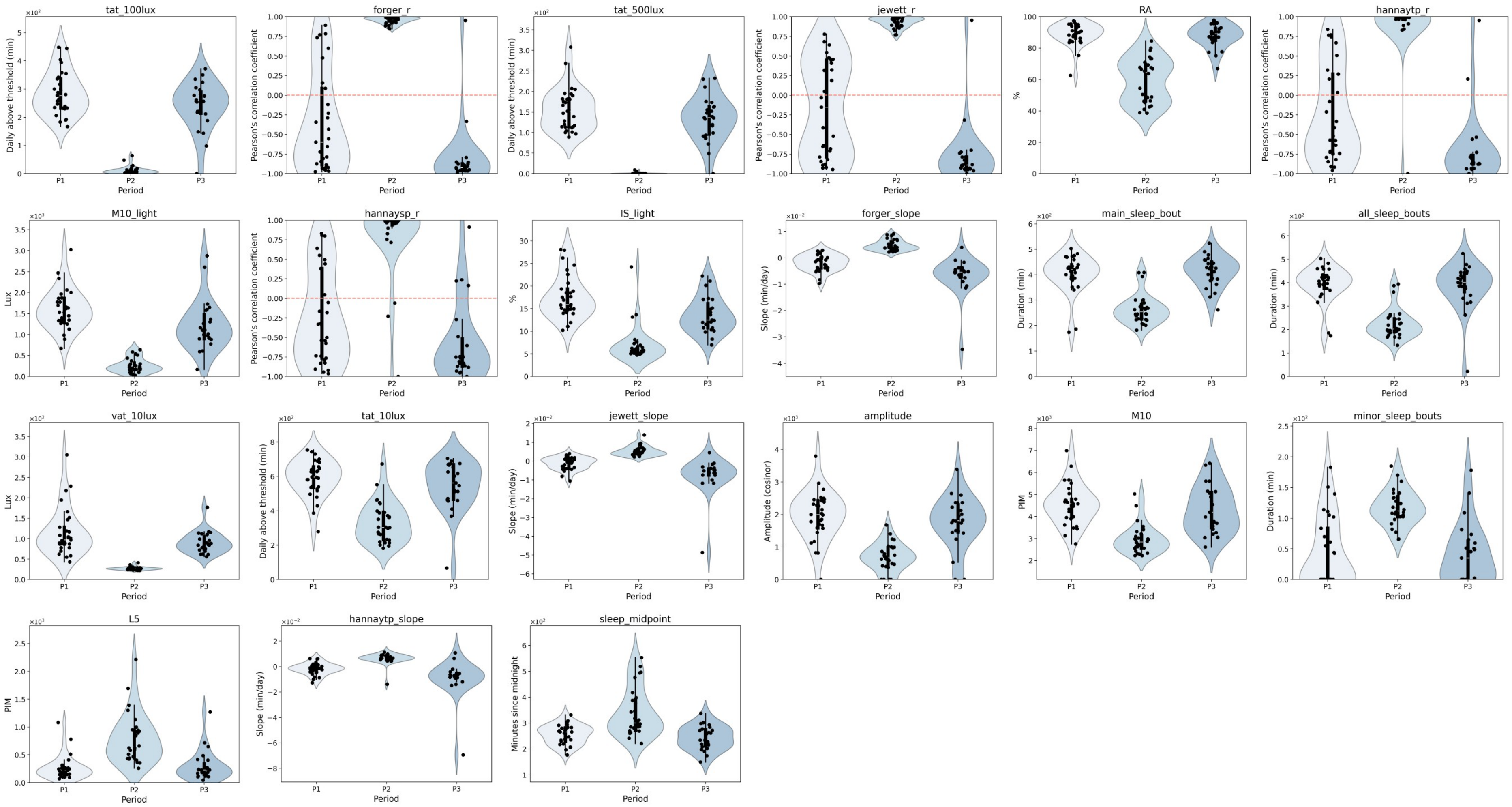

**Figure S3 Jittered violin plots for features that differentiated at least one study period.** Feature selection was initially based on a dual filtering criterion: statistical significance (FPR < 0.05) and effect size (*Cohen's f* > 0.63), the latter corresponding to the estimated minimum detectable effect size given the sample size used in this study (n = 29). Detailed descriptions of each feature are provided in Table S1.

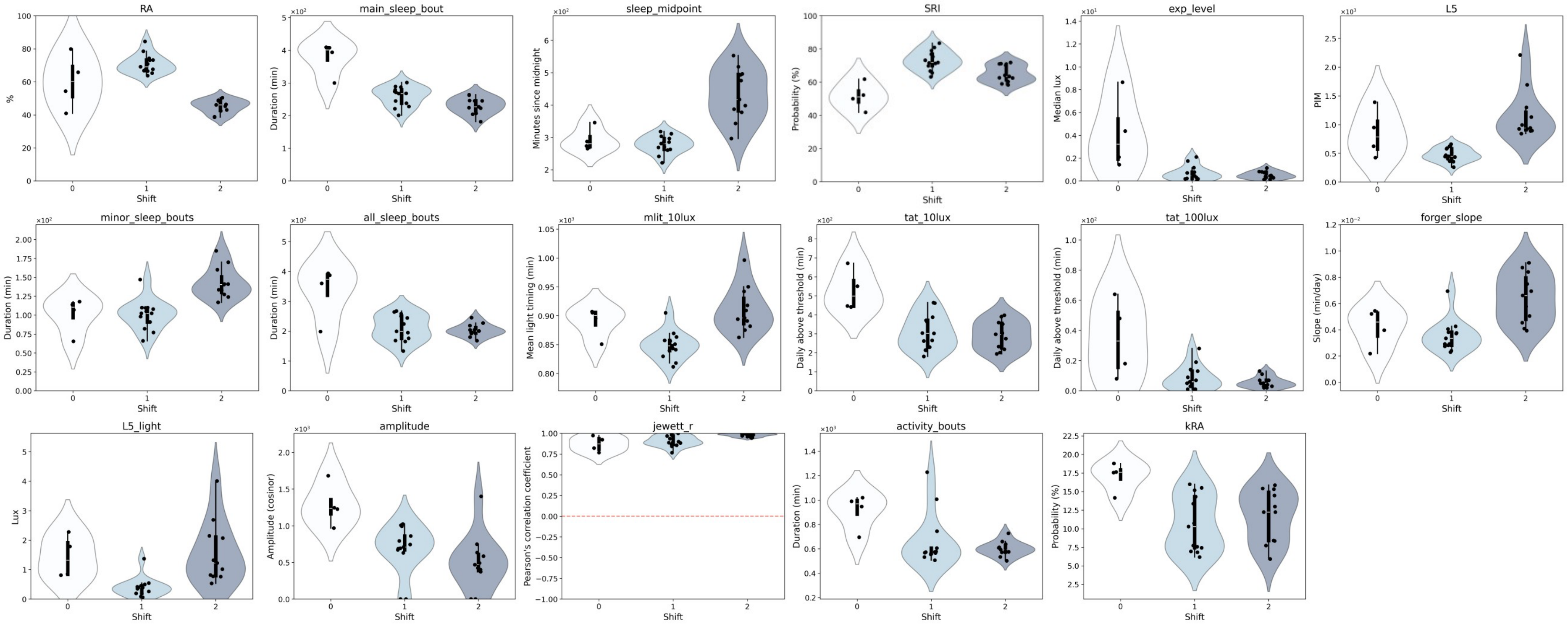

**Figure S4 Jittered violin plots for features that differentiated at least one shift during the submarine mission (P2).** Feature selection was initially based on a dual filtering criterion: statistical significance (FPR < 0.05) and effect size (*Cohen's f* > 0.63), the latter corresponding to the estimated minimum detectable effect size given the sample size used in this study (n = 29). Detailed descriptions of each feature are provided in Table S1.

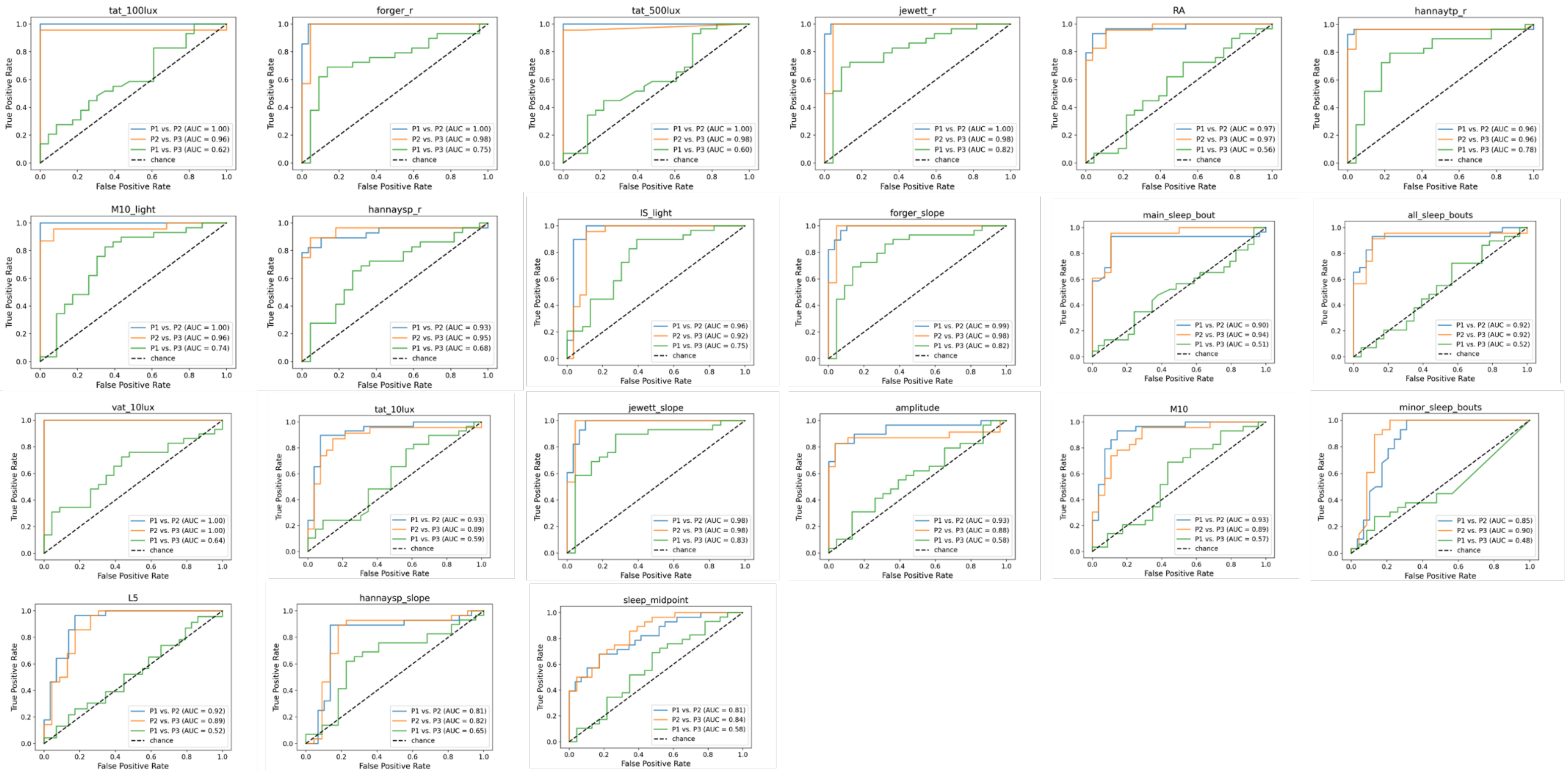

**Figure S5 ROC curves for features that differentiated at least one study period.** Pairwise discriminative performance was assessed via AUC, with significance determined through Monte Carlo permutation testing. This post-hoc evaluation was conducted only for features passing the initial screening criteria (FPR < 0.05 and *Cohen's f* > 0.63). Detailed descriptions of each feature are provided in Table S1.

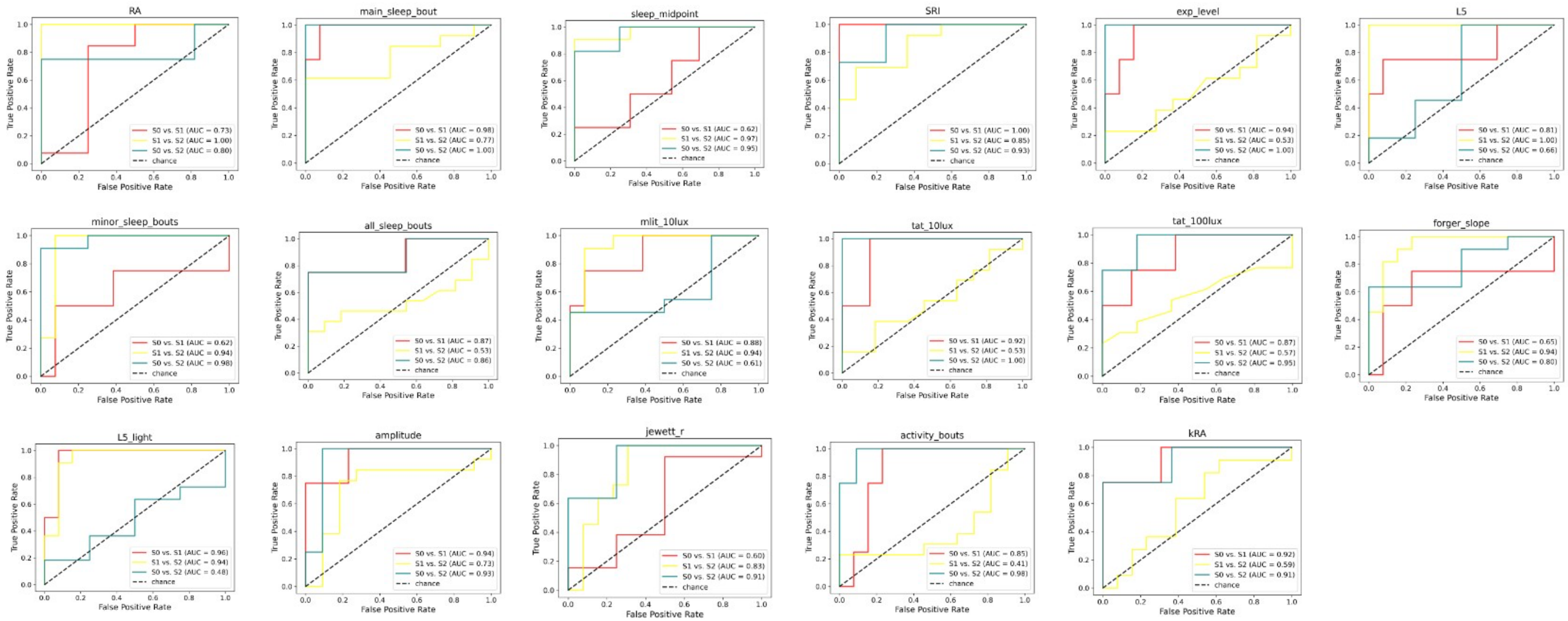

**Figure S6 ROC curves for features that differentiated at least one shift during the submarine mission (P2).** Pairwise discriminative performance was assessed via AUC, with significance determined through Monte Carlo permutation testing. This post-hoc evaluation was conducted only for features passing the initial screening criteria (FPR < 0.05 and *Cohen's f* > 0.63). Detailed descriptions of each feature are provided in Table S1.

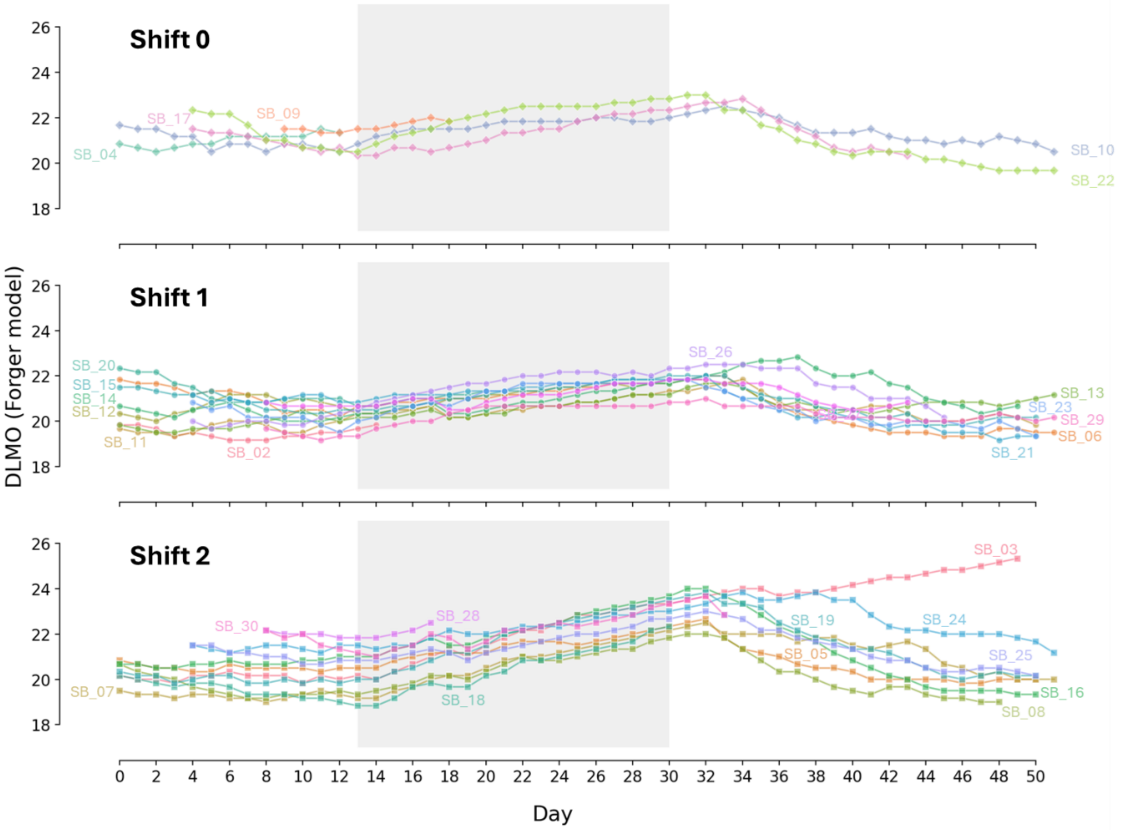

**Figure S7 Predicted daily circadian phase trajectory based on the Forger model.** Predicted dim light melatonin onset (DLMO) for each individual across the whole study, separated by shifts. The submarine mission period (P2) is shaded in light grey.

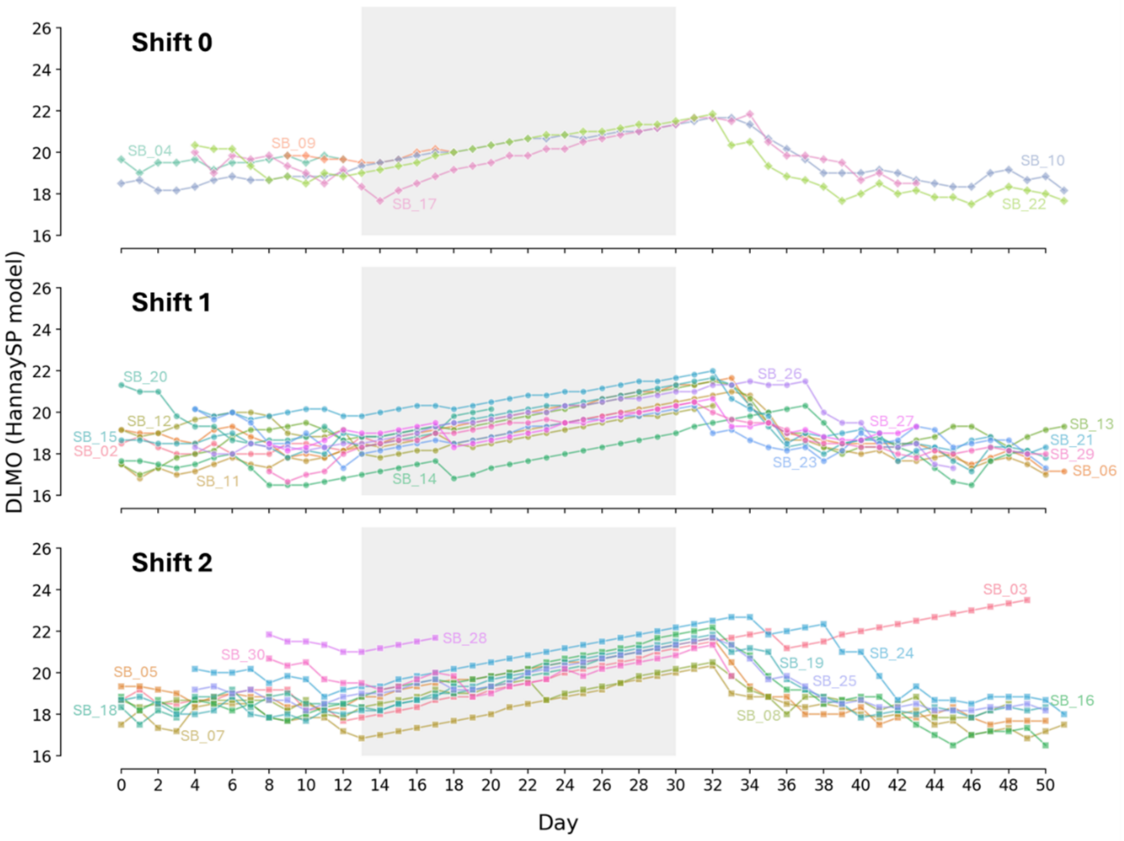

**Figure S8 Predicted daily circadian phase trajectory based on the HannaySP model.** Predicted dim light melatonin onset (DLMO) for each individual across the whole study, separated by shifts. The submarine mission period (P2) is shaded in light grey.

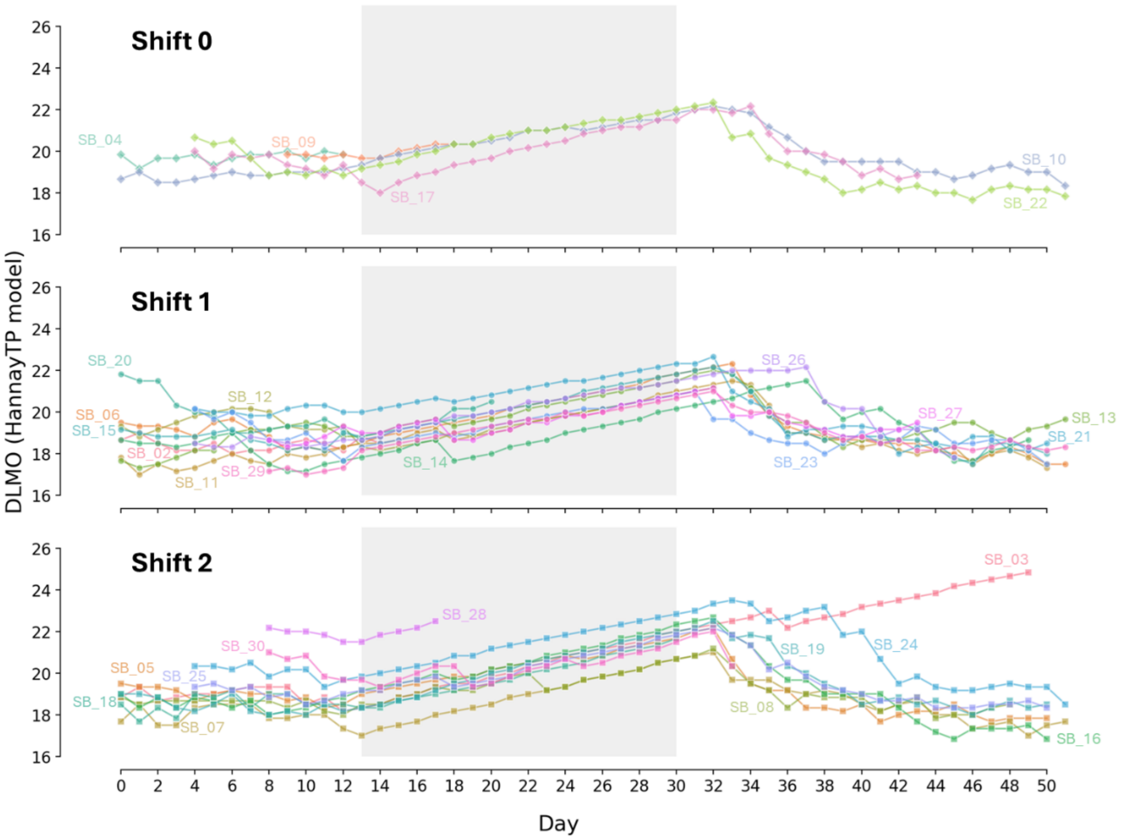

**Figure S9 Predicted daily circadian phase trajectory based on the HannayTP model**. Predicted dim light melatonin onset (DLMO) for each individual across the whole study, separated by shifts. The submarine mission period (P2) is shaded in light grey.
