## Supplementary Notebook for "An effect size-based screening approach to identify actigraphy markers of the submarine environment"

### submarine_analysis

June 16, 2025

### 1 Step-by-Step Analysis of the Submarine Pipeline

#### 1.1 Introduction

This Jupyter Notebook provides a step-by-step walkthrough of the data processing and analysis briefly described in the *Materials and Methods* section of the paper "Circadian disruption beneath the surface: a screening approach to identify actigraphy-based markers of split-shift work in submariners". It is intended to promote reproducibility and transparency of the analyses, while respecting the confidentiality of the raw data.

The notebook begins by importing the core Python libraries used thoughout the entire analysis pipeline for data wrangling, numerical computations, and data visualization. Additional libraries will be imported as needed in specific sections. Specifically, we import:

- `NumPy` for handling array data and performing matrix operations,
- `Pandas`for structured data manipulation using DataFrames,
- `scipy.stats` and `Pingouin` for scientific computing and statistical functions,
- `maplotlib.pyplot`, `seaborn`, and `plotly` for static and interactive data visualization,
- `os` for file navigation and handling,
- `re` for regular expression matching operations,
- `multiprocessing`, `jobliband` `tqdm` for parallel processing and monitoring execution time and progress,

[1]:
```
# Import core libraries common to most sections in the notebook
import numpy as np
import pandas as pd
from scipy import stats
import pingouin as pg
import matplotlib.pyplot as plt
import seaborn as sns
import re
import os
import time
import multiprocessing
from joblib import Parallel, delayed
from tqdm import tqdm
import warnings
```

##### 1.2 Computing the panel of actigraphy features using `circStudio`

**Import circStudio** `circStudio` is a version of `pyActigraphy` (v1.2.1) adapted for compatibility with Python 3.12 and extended with a module for mathematical modelling of circadian rhythms. This customized package encompasses all the necessary classes and functions to compute the full panel of actigraphy features described in the main manuscript. Future releases will offer both a programmatic interface and a GUI in the form a web-based application or dashboard.

We import `circStudio` and the `Cosinor` class from `circStudio.analysis`:

[2]:
```
import circStudio
from circStudio.analysis import Cosinor
```

**Calculate actigraphy features (rest-activity pattern, light-dark cycle, and sleep)** We process actigraphy files to compute features related to rest-activity patterns, the light-dark cycle, and sleep. briefly, for each actigraphy file located within the directory corresponding to a given study period, a panel of variables is computed and stored in a dataframe. Additional actigraphy-derived features, based on mathematical models of circadian rhythms, are computed in the following section.

**Note on `FutureWarning`:** A FutureWarning is raised regarding the behavior of DataFrame concatenation with empty or all-NA entries in `Pandas`. However, since the dataframes being concatenated do not contain any NaN entries, this warning can be safely ignored.

[ ]:
```
# Filter FutureWarning during the calculation of actigraphy features
warnings.filterwarnings("ignore", category=FutureWarning)

# Create a list of actigraphy-derived features to be computed
feature_set = ['sb_id','period','shift', 'ADAT','IS',␣
↪'IV','L5','M10','RA','exp_level',
                'tat_10lux','tat_100lux','tat_500lux', 'vat_10lux',␣
↪'vat_100lux', 'vat_500lux',
                'mlit_10lux', 'mlit_100lux','mlit_500lux', 'L5_light_onset',␣
↪'L5_light', 'M10_light_onset',
                'M10_light','IS_light','IV_light','SRI','sleep_midpoint',␣
↪'main_sleep_bout', 'minor_sleep_bouts',
                'all_sleep_bouts', 'activity_bouts', 'kRA', 'kAR', 'mesor',␣
↪'amplitude', 'acrophase']

# Create a new empty dataframe to store actigraphy-derived features
df = pd.DataFrame(columns=feature_set)

periods = ['p1', 'p2','p3']
for period in tqdm(periods, desc='Calculating actigraphy variables.'):
    # Create the directory path for actigraphy data of a given period
    source = os.path.join('..','data','processed','txt_divided', period)
```

```
    # Iterate over all the raw ATR files in the aforementioned directory
    for filename in os.listdir(source):
        # Generate a file path
        fpath = os.path.join(source, filename)

        # Check if that file path exists and if it has the correct extension (.
↪txt)
        if os.path.isfile(fpath) and filename.endswith('.txt'):
            # Load the ATR file, pre-trimmed in Condor ActStudio (v. 2.1.2) to␣
↪remove invalid start/end sequences
            raw = circStudio.io.read_raw_atr(fpath)

            # Mask spurious periods of inactivity
            ## 1) Reset any existing inactivity mask
            raw.inactivity_length = None

            ## 2) Mask periods of inactivity larger than or equal to 48 hours
            raw.create_inactivity_mask(duration='48h00min')

            ## 3) Add mask periods based on the visual inspection of spurious␣
↪periods of inactivity
            mask_name = os.path.splitext(os.path.basename(fpath))[0]
            raw.add_mask_periods(os.path.join('..', 'data',␣
↪'processed','missings',period,'csv', f'{mask_name}.csv'))

            ## 4) Activate the inactivity mask created in the previous steps
            raw.mask_inactivity= True

            # Retrieve the current submariner's shift from a shift lookup table
            identificator = pd.read_csv(os.path.join('..','data', 'raw',␣
↪'database','sb_shifts.csv'),index_col='sb_id')
            shift = identificator.loc[mask_name]['shift']

            # Calculate actigraphy-derived features
            ## Activity-related features (based on the non-binarized PIM signal)
            ADAT = raw.ADAT(binarize=False) # total average daily activity
            IS = raw.IS(freq='1h', binarize=False) # interdaily stability
            IV = raw.IV(freq='1h', binarize=False) # intradaily variability
            L5 = raw.L5(binarize=False) # Mean activity during the 5 least␣
↪active hours of the day
            M10 = raw.M10(binarize=False) # Mean activity during the 10 most␣
↪active hours of the day
            RA = raw.RA(binarize=False) # Relative amplitude

            ## Light-related features; pyActigraphy was customized to skip␣
↪log (+1) transformation
```

```
            ### Light interdaily stability
            light_is = raw.light.IS()
            light_is = light_is.loc[light_is['channel'] == 'LIGHT']['IS'].
↪values[0]

            ### Light intradaily variability
            light_iv = raw.light.IV()
            light_iv = light_iv.loc[light_iv['channel'] == 'LIGHT']['IV'].
↪values[0]

            ### Onset and mean light exposure levels during the 10 most bright␣
↪hours of the day
            m10_light = raw.light.LMX(length='10h',lowest=False)
            m10_light_onset = m10_light.loc[m10_light['channel'] ==␣
↪'LIGHT']['index'].values[0]
            m10_light_onset = m10_light_onset.total_seconds()/60 # Convert from␣
↪seconds to minutes
            m10_light = m10_light.loc[m10_light['channel'] == 'LIGHT']['value'].
↪values[0]

            ### Onset and mean light exposure levels during the 5 least bright␣
↪hours of the day
            l5_light = raw.light.LMX(length='5h', lowest=True)
            l5_light_onset = l5_light.loc[l5_light['channel'] ==␣
↪'LIGHT']['index'].values[0]
            l5_light_onset = l5_light_onset.total_seconds()/60 # Convert from␣
↪seconds to minutes
            l5_light = l5_light.loc[l5_light['channel'] == 'LIGHT']['value'].
↪values[0]

            ### Mean light timing above threshold (10, 100 and 500 lux)
            mlit_10lux = raw.light.MLiT(threshold=10).loc['LIGHT']
            mlit_100lux = raw.light.MLiT(threshold=100).loc['LIGHT']
            mlit_500lux = raw.light.MLiT(threshold=500).loc['LIGHT']

            ### Mean daily light exposure time above threshold (10, 100 and 500␣
↪lux)
            tat_10lux = raw.light.TATp(threshold=10, oformat='minute')['LIGHT'].
↪median()
            tat_100lux = raw.light.TATp(threshold=100,␣
↪oformat='minute')['LIGHT'].median()
            tat_500lux = raw.light.TATp(threshold=500,␣
↪oformat='minute')['LIGHT'].median()

            ### Mean light intensity value above threshold (10, 100 and 500 lux)
            vat_10lux = raw.light.VAT(threshold=10)['LIGHT'].median()
```

```python
            vat_100lux = raw.light.VAT(threshold=100)['LIGHT'].median()
            vat_500lux = raw.light.VAT(threshold=500)['LIGHT'].median()

            ### Median light exposure level per epoch
            exp_level = raw.light.light_exposure_level(agg='median').
↪loc['LIGHT']

            # Sleep-related features
            ## Sleep Regularity Index (using the Roenneberg algorithm)
            sri = raw.SleepRegularityIndex(freq='10min', algo='Roenneberg')

            ## Sleep Midpoint (using the Roenneberg algorithm)
            sleep_midpoint = raw.SleepMidPoint(freq='10min', to_td=False,␣
↪algo='Roenneberg')

            ## Median duration and IQR of major and minor sleep bouts
            ### 1) Create empty dataframe to store sleep bouts
            sleep_bouts = pd.DataFrame()

            ### 2) Calculate activity onset and offset
            activity_onset, activity_offset = raw.Roenneberg_AoT()

            #### 2.1) Assuming sleep_onset = activity_offset & sleep_offset =␣
↪activity_onset
            sleep_bouts['date'] = activity_onset.date
            sleep_bouts['start_time'] = activity_offset
            sleep_bouts['stop_time'] = activity_onset

            ### 3) Calculate the duration of the sleep/rest episode
            sleep_bouts['duration'] = sleep_bouts['stop_time'] -␣
↪sleep_bouts['start_time']

            ### 4) From the list of sleep bouts grouped by date, identify the␣
↪major sleep bout
            major_sleep_bouts = sleep_bouts.loc[sleep_bouts.
↪groupby('date')['duration'].idxmax()]

            #### 4.1) Remove major sleep bouts from the sleep bout set to get a␣
↪set of minor sleep bouts
            minor_sleep_bouts = sleep_bouts.drop(major_sleep_bouts.index)

            ### 5) Median duration for major sleep bout (minutes)
            average_duration_major = major_sleep_bouts['duration'].median().
↪total_seconds()/60

            ### 6) Median duration for minor sleep bouts (minutes)
```

```
            average_duration_minor = minor_sleep_bouts['duration'].median().
↪total_seconds()/60

            # Assign 0 if no minor sleep bouts are detected (biologically␣
↪meaningful: zero duration  missing data)
            if np.isnan(average_duration_minor):
                average_duration_minor = 0

            ## Median sleep bout duration across all sleep bouts (minutes)
            td = pd.Series(raw.sleep_durations(duration_min='10min',␣
↪algo='Roenneberg'))
            sleep_duration = td/pd.Timedelta('1min') # From Timedelta to minutes
            sleep_bout_median = sleep_duration.median() # Median

            ## Median activity duration (minutes)
            td_act = pd.Series(raw.active_durations(duration_min='10min',␣
↪algo='Roenneberg'))
            minutes_activity = td_act/pd.Timedelta('1min')
            activity_median = minutes_activity.median() # Median

            ## Probability of transitioning from rest to activity after␣
↪activity offset
            kra = raw.kRA(4, start='AoffT', freq='10min')

            ## Probability of transitioning from activity to rest after␣
↪activity onset
            kar = raw.kAR(4, start='AonT', freq='10min')

            ## Cosinor analysis
            ### 1) Initiate a cosinor object
            cosinor = Cosinor()

            ### 2) Disable inactivity mask (since Cosinor does not tolerate NaN␣
↪values)
            raw.mask_inactivity= False

            ### 3) Set and fix the period to 24-h (1440 minutes for a 60-second␣
↪sampling rate)
            cosinor.fit_initial_params['Period'].value = 1440
            cosinor.fit_initial_params['Period'].vary = False

            ### 4) Store the results from the cosinor fit and extract them
            results = cosinor.fit(raw.data, verbose=False)
            mesor = results.params['Mesor'].value
            amplitude = results.params['Amplitude'].value
            acrophase = results.params['Acrophase'].value
```

```
            ### Collect entry and add it to the dataframe
            df = pd.concat([df, pd.DataFrame({'sb_id': [mask_name], 'period':␣
↪[period], 'shift': [shift],
                                        'ADAT':[ADAT], 'IS': [IS], 'IV':␣
↪[IV], 'L5': [L5], 'M10': [M10],
                                        'RA': [RA], 'exp_level':␣
↪[exp_level], 'tat_10lux': [tat_10lux],
                                        'tat_100lux': [tat_100lux],␣
↪'tat_500lux': [tat_500lux], 'vat_10lux': [vat_10lux],
                                        'vat_100lux':␣
↪[vat_100lux],'vat_500lux': [vat_500lux],'mlit_10lux': [mlit_10lux],
                                        'mlit_10lux': [mlit_10lux],␣
↪'mlit_100lux': [mlit_100lux], 'mlit_500lux': [mlit_500lux],
                                        'L5_light_onset':␣
↪[l5_light_onset], 'L5_light': [l5_light],
                                        'M10_light_onset':␣
↪[m10_light_onset], 'M10_light': [m10_light], 'IS_light': [light_is],
                                        'IV_light': [light_iv], 'SRI':
↪[sri], 'sleep_midpoint':[sleep_midpoint],
                                        'main_sleep_bout':␣
↪[average_duration_major], 'minor_sleep_bouts': [average_duration_minor],
                                        'all_sleep_bouts':␣
↪[sleep_bout_median], 'activity_bouts': [activity_median], 'kRA': [kra],
                                        'kAR': [kar], 'mesor': [mesor],␣
↪'amplitude': [amplitude], 'acrophase': [acrophase]})], ignore_index=True)

# Save intermediate dataframe with actigraphy-derived features computed using␣
↪pyActigraphy
df.to_csv(os.path.join('..', 'data', 'processed', 'raw_act_features.csv'),␣
↪index=False)

# Reset the warning filter
warnings.resetwarnings()
```

**Predict the daily *DLMO* trajectory using light-informed models of circadian rhythms** The light intensity time series from the actigraphy recording is used approximate the dim light melatonin onset (*DLMO*) based on predictions from four mathematical models of circadian rhythms — two based on the Kronauer formalism (`Forger` and `Jewett`), and two on the Hannay formalism (`HannaySP` and `HannayTP`). Specifically, *DLMO* is estimated by applying a fixed 7-hour offset to the predicted core body temperature minimum ($CBT_{min}$ values. Although this method provides only a coarse approximation, it enables direct comparison with prior studies employing the same models, which typically report report *DLMO* rather than $CBT_{min}$ using the same 7-hour offset.

We import classes designed to process the light signal (`circStudio.light_tools`) and to calculate the aforementioned mathematical models (`circStudio.models.math_models`). Additionally, some circular statistics tools are imported (circStudio.models.tools).

[1]:
```
from circStudio.models.light_tools import *
from circStudio.models.math_models import *
from circStudio.models.tools import *
```

**Functions for light-based circadian modelling** Functions for processing submarine light data and calculate mathematical models

1. interpolate_zeros is a helper function designed to interpolate zeros within gaps of specified durations (dt_minand dt_max). This functionality is useful for mitigating the impact of off-wrist periods in the actigraphy recording. impute_nan fills NaN values with the mean of non-NaN at the same hour in different days within the same study period.

[ ]:
```
def interpolate_zeros(df, col_name, datetime_col, dt_min=5,dt_max=20):
    # Construct Boolean mask identifying which elements of the column vector␣
 ↪are zero
    df['is_zero'] = (df[col_name] == 0)

    # To identify group boundaries, compare each row in 'is_zeros' to its␣
 ↪vertically shifted version
    # Compute cumsum to count True values, corresponding to boundaries of␣
 ↪consecutive sequences of zeros
    # Each count of the lower bound of each consecutive group of zeros serves␣
 ↪as a zero group identifier
    # Multiply by df['is_zero'] to label nonzero values with zero (notice that␣
 ↪False * value = 0)
    df['zero_groups'] = ((df['is_zero'] != df['is_zero'].shift())
                         .cumsum()) * df['is_zero']

    # Iterate over each zero group to check if it is a valid duration
    for group in df['zero_groups'].unique():
        if group != 0:
            # Find the indices of zero group elements
            group_indices = df[df['zero_groups'] == group].index

            # Calculate length/duration of consecutive sequence of zeros
            duration = df.loc[group_indices[-1], datetime_col] - df.
 ↪loc[group_indices[0], datetime_col]

            if pd.Timedelta(minutes=dt_min - 1) <= duration <= pd.
 ↪Timedelta(minutes=dt_max+1):
                # Label any zero value in the light column with NaN
                df.loc[group_indices, col_name] = np.nan

                # Interpolate those NaN
                df[col_name] = df[col_name].interpolate()
```

```
    # Replace any remaining NaN with 0
    df.loc[df[col_name].isna(), col_name] = 0

    # Return the dataframe
    return df
```

[ ]:
```
def impute_nan(df, col_name, datetime_col, mask_address):
    # Open mask periods as a dataframe
    df_missing = pd.read_csv(mask_address)

    # Convert mask start and stop times to datetime
    df_missing['Start_time'] = pd.to_datetime(df_missing['Start_time'])
    df_missing['Stop_time'] = pd.to_datetime(df_missing['Stop_time'])

    for _, row in df_missing.iterrows():
        mask = (df[datetime_col] >= row['Start_time']) & (df[datetime_col] <=␣
↪row['Stop_time'])
        df.loc[mask, col_name] = np.nan

    # Fill NaN with the mean on the given time (math models do not allow for␣
↪gaps in the time series)
    df[col_name] = df[col_name].fillna(df[col_name].groupby(df[datetime_col].dt.
↪time).transform("mean"))

    # Return the original dataframe with NaN gaps imputed
    return df
```

2. The `light_pipeline` function streamlines the processing of light time series data. It trims incomplete days at the borders, resamples the data into 10-minute intervals, and converts the original datetime objects into hours since midnight, starting from the first day. The auxiliar function `smooth_light` can be used to apply smoothing filters. By default, we use a `median_filter`.

[ ]:
```
def smooth_light(light, window_size=10):
    return median_filter(light,window_size)
```

[ ]:
```
def light_pipeline(df, mask_path):
    # Convert 'DATE/TIME' type from str to datetime
    df['DATE/TIME'] = pd.to_datetime(df['DATE/TIME'], format="%d/%m/%Y %H:%M:
↪%S")

    # Linearly interpolate consecutive sequences of zeros between 5 and 45␣
↪minutes
    #df = interpolate_zeros(df, col_name='LIGHT', datetime_col='DATE/
↪TIME',dt_min=5, dt_max=45)
```

```
    df = impute_nan(df, col_name='LIGHT', datetime_col='DATE/TIME',␣
↪mask_address=mask_path)

    # Set DATE/TIME as the index for resampling
    df.set_index('DATE/TIME', inplace=True)

    # Resample to 10-minute intervals, taking the mean of numerical values
    df = df.resample('10min').mean()

    # Reset the index
    df.reset_index(inplace=True)

    # Create a new column for the date without time
    df['DATE'] = df['DATE/TIME'].dt.date

    # Find the first and last date
    first_day = df['DATE'].min()
    last_day = df['DATE'].max()

    # Exclude rows from the first and last days
    df = df[(df['DATE'] != first_day) & (df['DATE'] != last_day)]

    # Calculate hours elapsed from first datetime value
    filtered_df = df[(df['DATE'] != first_day) & (df['DATE'] != last_day)]
    filtered_min_time = filtered_df['DATE/TIME'].min()
    first_value = df['DATE/TIME'].min()
    df['HOURS'] = (df['DATE/TIME'] - filtered_min_time).dt.total_seconds() /␣
↪3600

    # Smooth light
    df['LIGHT'] = smooth_light(light = df['LIGHT'], window_size = 50)

    # Reset dataframe index
    df = df.reset_index()

    # Return the clean df
    return df
```

3. The submarine_pipeline function encapsulates the entire process of cleaning and analyzing submarine light intensity time series. By default, it finds initial conditions by looping over the first two complete days of actigraphy until the system reaches equilibrium - where successive iterations no longer produce changes in the results.

[ ]:
```
def submarine_pipeline(directory, save_address):
    def get_path(directory, shift):
        return os.path.join(directory,f'shift_{shift}')
```

```python
def compute_models(fpath, mask_path):
    def create_light(dataframe, mask_path):
        # Create a dataframe with 'DATE/TIME' and 'LIGHT' columns
        df_light = dataframe[['DATE/TIME', 'LIGHT']].copy()

        # Apply the light processing pipeline
        treated_df = light_pipeline(df_light, mask_path)

        # Retrieve the first two days of actigraphy data to calculate␣
↪initial conditions
        first_two_days = treated_df['DATE/TIME'].dt.date.drop_duplicates()[:
↪2]

        # Filter rows from the first two days
        ics_df = treated_df[treated_df['DATE/TIME'].dt.date.
↪isin(first_two_days)]

        # Create a return the light schedule
        schedule = Light(time_vector=treated_df.HOURS,␣
↪light_vector=treated_df.LIGHT)
        ics_schedule = Light(time_vector=ics_df.HOURS, light_vector=ics_df.
↪LIGHT)
        return schedule, ics_schedule, treated_df

    def create_model(time, light, time_ics, light_ics):
        # Model class list
        models = [Forger, Jewett, HannaySP, HannayTP]

        # Dictionary to store instances of each model
        model_objs = {}

        # Create instances of each model and populate model_instances
        for model in models:
            # Create model to compute initial conditions
            ics_model = model(time=time_ics, inputs=light_ics)
            ics_model.initial_condition = ics_model.
↪get_initial_conditions(time_vector=time_ics,
                                                                              ␣
↪light_vector=light_ics,
                                                                              ␣
↪loop_number=50
                                                                              )

            # Create model to compute the trajectory based on initial␣
↪conditions
            model_obj = model(time=time,inputs=light)
```

```
                model_obj.model_states = model_obj.integrate(time_vector=time,
                                                             light_vector=light,
                                                             ␣
↪initial_condition=ics_model.initial_condition)

                # Store the result in model collector dictionary
                model_name = model.__name__.lower()
                model_objs[model_name] = model_obj

            # Return the created model instances
            return model_objs

        # Open dataframe containing actigraphy data
        data = pd.read_csv(fpath)

        # Create light schedule
        schedule, ics_schedule, data = create_light(data, mask_path)

        # Create models
        models = create_model(time=np.asarray(schedule.time_vector),
                              light=np.asarray(schedule.light_vector),
                              time_ics=np.asarray(ics_schedule.time_vector),
                              light_ics=np.asarray(ics_schedule.light_vector)
                             )

        # Collect model outputs in a dictionary
        model_outputs = {
            "datetime": np.asarray(data['DATE/TIME']),
            "hours": schedule.time_vector,
            "light": schedule.light_vector
        }

        # Define model and model variable names
        model_var_dict = {
            'forger': ['x_forger', 'xc_forger', 'n_forger'],
            'jewett': ['x_jewett', 'xc_jewett', 'n_jewett'],
            'hannaysp': ['r_sp', 'phi_sp', 'n_sp'],
            'hannaytp': ['r1_tp', 'r2_tp', 'phi1_tp', 'phi2_tp', 'n_tp']
        }

        # Extract outputs from each model
        for model_name, output_vars in model_var_dict.items():
            model = models[model_name]
            for i, var_name in enumerate(output_vars):
                model_outputs[var_name] = model.model_states[:, i]

        # Return DataFrame with collected outputs
```

```
        return pd.DataFrame(model_outputs)

    for shift in range(3):
        source = get_path(directory, shift)
        for filename in os.listdir(source):
            fpath = os.path.join(source, filename)

            if os.path.isfile(fpath) and filename.endswith('.csv'):
                print(filename)
                mask_path = os.path.join('..','data','processed',␣
↪'missings_undivided', 'output_upper', f'{filename}')
                output_df = compute_models(fpath, mask_path)
                final_save_address = os.path.join(save_address,␣
↪f"shift_{shift}", f"{filename[:-4]}.csv")
                output_df.to_csv(final_save_address, index=False)
```

4. Finally, run the entire mathematical model pipeline to get a time series of model states!

[ ]:
```
# Source and destination
source = os.path.join('..','data','processed','csv_not_divided')
save_address = os.path.join('..', 'data', 'analyzed', 'entrained')

# Calculate model states based self-convergence
submarine_pipeline(source, save_address)
```

Functions to calculate $CBT_{min}$, $DLMO$, phase, and amplitude from model states

Now that we have calculated a time series of model states, we can extract biologically meaningful quantities, such as the predicted $CBT_{min}$, $DLMO$, phase, and rhythm amplitude.

1. Compute `dlmo` and `cbtmin` vectors from model states stored in `pandas` DataFrames:

[ ]:
```
# Forger and Jewett models (Kronauer-based, Van der Pol (VDP) models)
def cbtmin_vdp(time, x):
    # Calculate time step (dt) between consecutive time points
    dt = np.diff(time)[0]

    # Invert cos(x) to turn the minima into maxima (peaks)
    inverted_x = -1.0 * x

    # Identify the indices where minima occur
    cbtmin_indices, _ = find_peaks(inverted_x, distance=np.ceil(13.0/dt))

    # Use the previous indices to find the cbtmin times
    cbtmin_times = time[cbtmin_indices]

    # To convert to clock time -> cbtmin_times % 24
    return cbtmin_times
```

```
# HannaySP and HannayTP models
def cbtmin_hannay(time,phase):
    # Calculate time step (dt) between consecutive time points
    dt = np.diff(time)[0]

    # Invert cos(x) to turn the minima into maxima (peaks)
    inverted = -1.0 * np.cos(phase)

    # Identify the indices where minima occur
    cbtmin_indices, _ = find_peaks(inverted, distance=np.ceil(13.0/dt))

    # Use the previous indices to find the cbtmin times
    cbtmin_times = time[cbtmin_indices]

    # To convert to clock time -> cbtmin_times % 24
    return cbtmin_times

# Compute dlmos (both VDP and Hannay models)
def dlmos(cbt_vector, cbt_to_dlmo=7):
    return (cbt_vector - cbt_to_dlmo) % 24
```

2. Compute the phase and amplitude based on model states:

[ ]:
```
# Compute amplitude for VDP models (Forger and Jewett)
def amplitude_vdp(x,xc):
    return np.sqrt(x**2, xc**2)

def phase_vdp(x,xc):
    xc = -1.0 * xc
    return np.angle(x + complex(0,1) * xc)

def phase_hannay(phase):
    x = np.cos(phase)
    y = np.sin(phase)
    return np.angle(x + complex(0,1) * y)
```

3. Although the procedures for calculating these metrics across models are similar, we opted to keep them separate for clarity. All resulting dataframes are merged at the end of the pipeline.

[ ]:
```
def forger_params(fpath, sub_id, shift):
    # Create dataframe with model states
    df = pd.read_csv(fpath)

    # Extract the date from the datetime column
    df['datetime'] = pd.to_datetime(df['datetime'])
    df['date'] = df['datetime'].dt.date
```

```
    # Compute parameters using the entire columns
    cbtmin_values = cbtmin_vdp(df.hours, df.x_forger)
    dlmo_values = dlmos(cbtmin_values)

    # Pad shorter columns with NaN
    max_length = max(len(df["date"].unique()), len(cbtmin_values),␣
↪len(dlmo_values))
    dates = np.pad(df["date"].unique(), (0, max_length - len(df["date"].
↪unique())), constant_values=np.nan)
    cbtmin_values = np.pad(cbtmin_values, (0, max_length - len(cbtmin_values)),␣
↪constant_values=np.nan)
    dlmo_values = np.pad(dlmo_values, (0, max_length - len(dlmo_values)),␣
↪constant_values=np.nan)

    forger_df = pd.DataFrame({'date': dates,
                              'sub_id': sub_id,
                              'shift': shift,
                              'model': 'forger',
                              'cbtmin': cbtmin_values,
                              'dlmo': dlmo_values
                             })

    return forger_df
```

```
[ ]: def jewett_params(fpath, sub_id, shift):
    # Create dataframe with model states
    df = pd.read_csv(fpath)

    # Extract the date from the datetime column
    df['datetime'] = pd.to_datetime(df['datetime'])
    df['date'] = df['datetime'].dt.date

    # Compute parameters using the entire columns
    cbtmin_values = cbtmin_vdp(df.hours, df.x_jewett)
    dlmo_values = dlmos(cbtmin_values)

    # Pad shorter columns with NaN
    max_length = max(len(df["date"].unique()), len(cbtmin_values),␣
↪len(dlmo_values))
    dates = np.pad(df["date"].unique(), (0, max_length - len(df["date"].
↪unique())), constant_values=np.nan)
    cbtmin_values = np.pad(cbtmin_values, (0, max_length - len(cbtmin_values)),␣
↪constant_values=np.nan)
    dlmo_values = np.pad(dlmo_values, (0, max_length - len(dlmo_values)),␣
↪constant_values=np.nan)
```

```
    jewett_df = pd.DataFrame({'date': dates,
                              'sub_id': sub_id,
                              'shift': shift,
                              'model': 'jewett',
                              'cbtmin': cbtmin_values,
                              'dlmo': dlmo_values
                             })

    return jewett_df
```

[ ]:
```
def hannaysp_params(fpath, sub_id, shift):
    # Open DataFrame containing model states
    df = pd.read_csv(fpath)

    # Extract the date from the datetime column
    df['datetime'] = pd.to_datetime(df['datetime'])
    df['date'] = df['datetime'].dt.date

    # Compute parameters using the entire columns
    cbtmin_values = cbtmin_hannay(df.hours, df.phi_sp)
    dlmo_values = dlmos(cbtmin_values)

    # Pad shorter columns with NaN
    max_length = max(len(df["date"].unique()), len(cbtmin_values),␣
↪len(dlmo_values))
    dates = np.pad(df["date"].unique(), (0, max_length - len(df["date"].
↪unique())), constant_values=np.nan)
    cbtmin_values = np.pad(cbtmin_values, (0, max_length - len(cbtmin_values)),␣
↪constant_values=np.nan)
    dlmo_values = np.pad(dlmo_values, (0, max_length - len(dlmo_values)),␣
↪constant_values=np.nan)

    hannaysp_df = pd.DataFrame({'date': dates,
                              'sub_id': sub_id,
                              'shift': shift,
                              'model': 'hannaysp',
                              'cbtmin': cbtmin_values,
                              'dlmo': dlmo_values
                             })

    return hannaysp_df
```

[ ]:
```
def hannaytp_params(fpath, sub_id, shift):
    # Open DataFrame containing model states
    df = pd.read_csv(fpath)

    # Extract the date from the datetime column
```

```
    df['datetime'] = pd.to_datetime(df['datetime'])
    df['date'] = df['datetime'].dt.date

    # Compute parameters using the entire columns
    cbtmin_values = cbtmin_hannay(df.hours, df.phi1_tp)
    dlmo_values = dlmos(cbtmin_values)

    # Pad shorter columns with NaN
    max_length = max(len(df["date"].unique()), len(cbtmin_values),␣
 ↪len(dlmo_values))
    dates = np.pad(df["date"].unique(), (0, max_length - len(df["date"].
 ↪unique())), constant_values=np.nan)
    cbtmin_values = np.pad(cbtmin_values, (0, max_length - len(cbtmin_values)),␣
 ↪constant_values=np.nan)
    dlmo_values = np.pad(dlmo_values, (0, max_length - len(dlmo_values)),␣
 ↪constant_values=np.nan)

    hannaytp_df = pd.DataFrame({'date': dates,
                              'sub_id': sub_id,
                              'shift': shift,
                              'model': 'hannaytp',
                              'cbtmin': cbtmin_values,
                              'dlmo': dlmo_values
                             })

    return hannaytp_df
```

4. Collect predicted $DLMO$ and $CBT_{min}$ for different models on a single dataframe.

[ ]:
```
# Source and save directories
source_address = os.path.join('..','data','analyzed','entrained')
save_address = os.path.join('..', 'data', 'analyzed', 'params','dlmo_cbtmin')

# Model function
param_funcs = [forger_params, jewett_params, hannaysp_params, hannaytp_params]
dataframes = []

# Iterave over parameter functions and shifts
for param_func in param_funcs:
    for shift in range(3):
        def get_path(directory, shift_num):
            return os.path.join(directory, f'shift_{shift_num}')
        source = get_path(source_address, shift)

        for filename in os.listdir(source):
            fpath = os.path.join(source, filename)
            if os.path.isfile(fpath) and filename.endswith('.csv'):
```

```
                params = param_func(fpath,sub_id=filename[:-4],shift=shift)
                dataframes.append(params)

# Concatenate all dataframes and save them on an intermediate dataframe
df_params = pd.concat(dataframes,ignore_index=True)

# Save intermediary dataframe containing dlmo and cbtmin values
df_params.to_csv(os.path.join(save_address, "dlmo_cbtmin.csv"), index=False)
```

Fitting a regression line to *DLMO* time series by period

A **regression line** is fitted to each predicted *DLMO* time series. We evaluate the strength and direction of a linear dependence with time - using the Pearson's correlation coefficient ($r$), indicative of phase advances or delays, and the rate of change of the *DLMO* trajectory using the slope of the regression line.

[ ]:
```
# Create a list with start and end dates corresponding to the start and end of␣
↪each study period
start_dates = [pd.to_datetime('yyyy-mm-dd'), pd.to_datetime('yyyy-mm-dd'), pd.
↪to_datetime('yyyy-mm-dd')]
end_dates = [pd.to_datetime('yyyy-mm-dd'), pd.to_datetime('yyyy-mm-dd'), pd.
↪to_datetime('yyyy-mm-dd')]

# Create a list with shift labels
shifts = [0, 1, 2]

# Create a list of models
models = ['forger', 'jewett', 'hannaysp', 'hannaytp']

# Create an empty dataframe to store regression results
df_output = pd.DataFrame()


for shift in shifts:
    for i, (start, end) in enumerate(zip(start_dates, end_dates), start=1):
        reg_dict = {}

        for model in models:
            # Filter input dataframe for a specific model, shift, and period
            df_filtered = df_input[(df_input['model'] == model) &␣
↪(df_input['shift'] == shift) & (df_input['date'].between(start, end))].
↪dropna()
            df_filtered['period'] = f'p{i}'

            # Define t0 (zero hours elapsed since the beginning)
            first_day = df_filtered['date'].min().normalize()
```

```
            # Convert dates to hours since first day
            df_filtered['t'] = (df_filtered['date'] - first_day).dt.
↪total_seconds() / 3600

            # Group by sb_id
            for sb, df_sb in df_filtered.groupby('sb_id'):
                # Perform linear regression
                res = stats.linregress(df_sb['t'], df_sb['cbtmin'] % 24)

                # Store results in dictionary
                if (sb, shift, f'p{i}') not in reg_dict:
                    reg_dict[(sb, shift, f'p{i}')] = {'sb_id': sb.lower(),␣
↪'shift': shift, 'period': f'p{i}'}

                reg_dict[(sb, shift, f'p{i}')].update({
                    f'{model}_r': res.rvalue,
                    f'{model}_slope': res.slope
                })

        if reg_dict:
            df_model_results = pd.DataFrame.from_dict(reg_dict, orient='index')
            df_output = pd.concat([df_output, df_model_results],␣
↪ignore_index=True)

# Save the intermediate dataframe containing regression results
df_output.to_csv(os.path.join('..', 'data', 'processed', 'regression_models.
↪csv'), index=False)
```

Visualizing the predicted *DLMO* daily trajectory

Jewett model:

[ ]:
```
# Set Seaborn color palette
sns.set_palette("Set2")

# Load model-derived circadian phase data (DLMO and CBTmin estimates)
path = os.path.join('..', 'data', 'analyzed', 'params', 'dlmo_cbtmin',␣
↪'dlmo_cbtmin.csv')
df = pd.read_csv(path)

# Convert 'date' series to a pandas datetime object
df["date"] = pd.to_datetime(df["date"])

# Adjust DLMO values occurring after midnight for better visualization
df["dlmo_adjusted"] = df["dlmo"].apply(lambda x: x + 24 if x < 6 else x)

# Remove any rows with missing values
df = df.dropna()
```

```python
# Create a 3-panel figure to display each shift DLMO daily progression
fig, axes = plt.subplots(nrows=3, ncols=1, figsize=(12,9), sharey=True,␣
 ↪sharex='col', dpi=200)
axes = axes.flatten()

# Define shifts and unique marker style by subplot
shifts = [0, 1, 2]
marker_list = ['D', 'o', 's']

for i, shift in enumerate(shifts):
    # Filter data for the current shift group and the Jewett model
    data_filtered = df[(df['model'] == 'jewett') & (df['shift'] == shift)]

    # Highlight mission period (replace with actual start/end dates)
    axes[i].axvspan(pd.to_datetime('yyyy-mm-dd'), pd.to_datetime('yyyy-mm-dd'),␣
 ↪color='grey', alpha=0.12, lw=0)

    # Plot adjusted DLMO time series by submariner
    sns.lineplot(data=data_filtered, x='date', y='dlmo_adjusted', hue='sb_id',
                 marker=marker_list[i], markersize=5, alpha=0.6, ax=axes[i],␣
 ↪linewidth = 1.25)

    # Remove legend (labels are added manually in PowerPoint)
    axes[i].get_legend().remove()

    # Format y-axis label only for the middle subplot
    if i == 1:
        axes[i].set_ylabel('DLMO (Jewett model)', fontsize=16)
    else:
        axes[i].set_ylabel('')

    # Remove x-axis label for all subplots (add one globally later)
    axes[i].set_xlabel('')

    # Set y-axis limits and ticks
    axes[i].set_ylim(17, 27)
    axes[i].set_yticks([18, 20, 22, 24, 26])
    axes[i].tick_params(axis='both', labelsize=12)

    # Reduce x-axis clutter by keeping every other day (0, 2, 4, ...)
    unique_dates = sorted(data_filtered['date'].dt.floor('D').unique())
    custom_dates = unique_dates[::2]
    custom_labels = [f'{i}' for i in range(0, len(unique_dates), 2)]
    axes[i].set_xticks(custom_dates)
    axes[i].set_xticklabels(custom_labels, rotation=0)
```

```
    # Remove top and right plot spines
    sns.despine(offset=10, trim=True)

# Add a shared x-axis title
fig.supxlabel('Day', fontsize=16)

# Adjust layout to prevent overlap
plt.tight_layout()

# Display the figure
plt.show()
```

Forger model:

```
[ ]: # Set Seaborn color palette
sns.set_palette("Set2")

# Load model-derived circadian phase data (DLMO and CBTmin estimates)
path = os.path.join('..', 'data', 'analyzed', 'params', 'dlmo_cbtmin',␣
 ↪'dlmo_cbtmin.csv')
df = pd.read_csv(path)

# Convert 'date' series to a pandas datetime object
df["date"] = pd.to_datetime(df["date"])

# Adjust DLMO values occurring after midnight for better visualization
df["dlmo_adjusted"] = df["dlmo"].apply(lambda x: x + 24 if x < 6 else x)

# Remove any rows with missing values
df = df.dropna()

# Create a 3-panel figure to display each shift DLMO daily progression
fig, axes = plt.subplots(nrows=3, ncols=1, figsize=(12,9), sharey=True,␣
 ↪sharex='col', dpi=200)
axes = axes.flatten()

# Define shifts and unique marker style by subplot
shifts = [0, 1, 2]
marker_list = ['D', 'o', 's']

for i, shift in enumerate(shifts):
    # Filter data for the current shift group and the Forger model
    data_filtered = df[(df['model'] == 'forger') & (df['shift'] == shift)]

    # Highlight mission period (replace with actual start/end dates)
    axes[i].axvspan(pd.to_datetime('yyyy-mm-dd'), pd.to_datetime('yyyy-mm-dd'),␣
 ↪color='grey', alpha=0.12, lw=0)
```

```
    # Plot adjusted DLMO time series by submariner
    sns.lineplot(data=data_filtered, x='date', y='dlmo_adjusted', hue='sb_id',
                 marker=marker_list[i], markersize=5, alpha=0.6, ax=axes[i],␣
 ↪linewidth = 1.25)

    # Remove legend (labels are added manually in PowerPoint)
    axes[i].get_legend().remove()

    # Format y-axis label only for the middle subplot
    if i == 1:
        axes[i].set_ylabel('DLMO (Forger model)', fontsize=16)
    else:
        axes[i].set_ylabel('')

    # Remove x-axis label for all subplots (add one globally later)
    axes[i].set_xlabel('')

    # Set y-axis limits and ticks
    axes[i].set_ylim(17, 27)
    axes[i].set_yticks([18, 20, 22, 24, 26])
    axes[i].tick_params(axis='both', labelsize=12)

    # Reduce x-axis clutter by keeping every other day (0, 2, 4, ...)
    unique_dates = sorted(data_filtered['date'].dt.floor('D').unique())
    custom_dates = unique_dates[::2]
    custom_labels = [f'{i}' for i in range(0, len(unique_dates), 2)]
    axes[i].set_xticks(custom_dates)
    axes[i].set_xticklabels(custom_labels, rotation=0)

    # Remove top and right plot spines
    sns.despine(offset=10, trim=True)

# Add a shared x-axis title
fig.supxlabel('Day', fontsize=16)

# Adjust layout to prevent overlap
plt.tight_layout()

# Display the figure
plt.show()
```

HannaySP:

```
[ ]: # Set Seaborn color palette
     sns.set_palette("Set2")
```

```python
# Load model-derived circadian phase data (DLMO and CBTmin estimates)
path = os.path.join('..', 'data', 'analyzed', 'params', 'dlmo_cbtmin',␣
↪'dlmo_cbtmin.csv')
df = pd.read_csv(path)

# Convert 'date' series to a pandas datetime object
df["date"] = pd.to_datetime(df["date"])

# Adjust DLMO values occurring after midnight for better visualization
df["dlmo_adjusted"] = df["dlmo"].apply(lambda x: x + 24 if x < 6 else x)

# Remove any rows with missing values
df = df.dropna()

# Create a 3-panel figure to display each shift DLMO daily progression
fig, axes = plt.subplots(nrows=3, ncols=1, figsize=(12,9), sharey=True,␣
↪sharex='col', dpi=200)
axes = axes.flatten()

# Define shifts and unique marker style by subplot
shifts = [0, 1, 2]
marker_list = ['D', 'o', 's']

for i, shift in enumerate(shifts):
    # Filter data for the current shift group and the HannaySP model
    data_filtered = df[(df['model'] == 'hannaysp') & (df['shift'] == shift)]

    # Highlight mission period (replace with actual start/end dates)
    axes[i].axvspan(pd.to_datetime('yyyy-mm-dd'), pd.to_datetime('yyyy-mm-dd'),␣
↪color='grey', alpha=0.12, lw=0)

    # Plot adjusted DLMO time series by submariner
    sns.lineplot(data=data_filtered, x='date', y='dlmo_adjusted', hue='sb_id',
                 marker=marker_list[i], markersize=5, alpha=0.6, ax=axes[i],␣
↪linewidth = 1.25)

    # Remove legend (labels are added manually in PowerPoint)
    axes[i].get_legend().remove()

    # Format y-axis label only for the middle subplot
    if i == 1:
        axes[i].set_ylabel('DLMO (HannaySP model)', fontsize=16)
    else:
        axes[i].set_ylabel('')

    # Remove x-axis label for all subplots (add one globally later)
    axes[i].set_xlabel('')
```

```
    # Set y-axis limits and ticks
    axes[i].set_ylim(17, 27)
    axes[i].set_yticks([16, 18, 20, 22, 24, 26])
    axes[i].tick_params(axis='both', labelsize=12)

    # Reduce x-axis clutter by keeping every other day (0, 2, 4, ...)
    unique_dates = sorted(data_filtered['date'].dt.floor('D').unique())
    custom_dates = unique_dates[::2]
    custom_labels = [f'{i}' for i in range(0, len(unique_dates), 2)]
    axes[i].set_xticks(custom_dates)
    axes[i].set_xticklabels(custom_labels, rotation=0)

    # Remove top and right plot spines
    sns.despine(offset=10, trim=True)

# Add a shared x-axis title
fig.supxlabel('Day', fontsize=16)

# Adjust layout to prevent overlap
plt.tight_layout()

# Display the figure
plt.show()
```

HannayTP:

```
[ ]: # Set Seaborn color palette
sns.set_palette("Set2")

# Load model-derived circadian phase data (DLMO and CBTmin estimates)
path = os.path.join('..', 'data', 'analyzed', 'params', 'dlmo_cbtmin',␣
 ↪'dlmo_cbtmin.csv')
df = pd.read_csv(path)

# Convert 'date' series to a pandas datetime object
df["date"] = pd.to_datetime(df["date"])

# Adjust DLMO values occurring after midnight for better visualization
df["dlmo_adjusted"] = df["dlmo"].apply(lambda x: x + 24 if x < 6 else x)

# Remove any rows with missing values
df = df.dropna()

# Create a 3-panel figure to display each shift DLMO daily progression
fig, axes = plt.subplots(nrows=3, ncols=1, figsize=(12,9), sharey=True,␣
 ↪sharex='col', dpi=200)
```

```python
axes = axes.flatten()

# Define shifts and unique marker style by subplot
shifts = [0, 1, 2]
marker_list = ['D', 'o', 's']

for i, shift in enumerate(shifts):
    # Filter data for the current shift group and the HannayTP model
    data_filtered = df[(df['model'] == 'hannaytp') & (df['shift'] == shift)]

    # Highlight mission period (replace with actual start/end dates)
    axes[i].axvspan(pd.to_datetime('yyyy-mm-dd'), pd.to_datetime('yyyy-mm-dd'),␣
 ↪color='grey', alpha=0.12, lw=0)

    # Plot adjusted DLMO time series by submariner
    sns.lineplot(data=data_filtered, x='date', y='dlmo_adjusted', hue='sb_id',
                 marker=marker_list[i], markersize=5, alpha=0.6, ax=axes[i],␣
 ↪linewidth = 1.25)

    # Remove legend (labels are added manually in PowerPoint)
    axes[i].get_legend().remove()

    # Format y-axis label only for the middle subplot
    if i == 1:
        axes[i].set_ylabel('DLMO (HannayTP model)', fontsize=16)
    else:
        axes[i].set_ylabel('')

    # Remove x-axis label for all subplots (add one globally later)
    axes[i].set_xlabel('')

    # Set y-axis limits and ticks
    axes[i].set_ylim(17, 27)
    axes[i].set_yticks([16, 18, 20, 22, 24, 26])
    axes[i].tick_params(axis='both', labelsize=12)

    # Reduce x-axis clutter by keeping every other day (0, 2, 4, ...)
    unique_dates = sorted(data_filtered['date'].dt.floor('D').unique())
    custom_dates = unique_dates[::2]
    custom_labels = [f'{i}' for i in range(0, len(unique_dates), 2)]
    axes[i].set_xticks(custom_dates)
    axes[i].set_xticklabels(custom_labels, rotation=0)

    # Remove top and right plot spines
    sns.despine(offset=10, trim=True)

# Add a shared x-axis title
```

```
fig.supxlabel('Day', fontsize=16)

# Adjust layout to prevent overlap
plt.tight_layout()

# Display the figure
plt.show()
```

**Join all features into a single dataframe** Finally, we need to combine both `regression_models` and `raw_act_features` dataframes into a `combined_df` that we will use for further analyses.

[10]:
```
# Open dataframes to merge
df1 = pd.read_csv(os.path.join('..','data','processed','raw_act_features.csv'))
df2 = pd.read_csv(os.path.join('..','data','processed','regression_models.csv'))

# Merge dataframes based on union logic
combined_df = pd.merge(df1, df2, on=['sb_id', 'period', 'shift'], how='outer')

# Save the newly created combined dataframe
combined_df.to_csv(os.path.join('..', 'data', 'processed','combined_df.csv'),␣
 ↪index=False)
```

##### 1.3 Principal component analysis (PCA)

The main sources of variance in the submarine dataset are identified via principal component analysis (PCA). The following section will present the major steps followed.

1. Load the `combined_df` dataframe obtained in earlier steps, and extract the metadata (i.e., shift and period information) into a new dataframe which will be later used for coloring by shift and period.

[21]:
```
# Open the combined_df.csv file created earlier
raw = pd.read_csv(os.path.join('..', 'data', 'processed', 'combined_df.csv'))

# Fuse the sb_id and period labels into a new id column
raw['id'] = raw['sb_id'].astype(str) + '_' + raw['period']

# Set the resulting 'id' column as index
raw = raw.set_index('id')

# Save period and shift for coloring in the meta dataframe
meta = raw[['period', 'shift']].copy()

# Drop rows containing NaNs
raw = raw.dropna(axis='rows', how='any')
```

```python
# Remove matadata (non-numeric) columns
rm = ['sb_id', 'period', 'shift']
raw = raw.drop(columns=rm)

# Drop rows containing NaNs
raw = raw.dropna(axis='rows', how='any')

# Align meta with remaining rows
meta = meta.loc[raw.index]
```

2. Prior to PCA, it is important perform data normalization. Import the `sklearn.preprocessing` module to center and scale the data. After this transformation, each actigraphy-derived feature will have a mean of 0 and a standard deviatio of 1 ($Z$-scores).

[22]:
```python
from sklearn import preprocessing
```

[23]:
```python
normalized_data = preprocessing.scale(raw)
```

3. Using the PCA class from `sklearn.decomposition`, create a new PCA object and fit it to the normalized data.

[24]:
```python
from sklearn.decomposition import PCA
```

[25]:
```python
# Create a new PCA object
pca = PCA()

# Fit the normalized data to the new PCA object
pca.fit(normalized_data)

# Generate coordinates for the PCA plot, based on the loading scores and␣
↪normalized data
pca_data = pca.transform(normalized_data)
```

4. Now, create a scree plot and PCA plots colored by period and by shift. To make sure that we are using the `matplotlib` original style, reset to defaults using the `seaborn` library.

[26]:
```python
# Reset to matplotlib default palettes
sns.reset_defaults()
```

[27]:
```python
# Calculate the percentage of variation for each principal component (PC)
pca_var = np.round(pca.explained_variance_ratio_ * 100, decimals=1)

# Generate labels
labels = [f'PC{str(x)}' for x in range(1, len(pca_var) + 1)]

# Define the number of top components
```

```
top_n = 10
plt.figure(dpi=100)
# Create a scree plot with the top n PCs
plt.bar(x=range(1, top_n + 1), height=pca_var[:top_n], tick_label=labels[:
 ↪top_n], facecolor='cornflowerblue')

# Label the y axis.
plt.ylabel('Percentage of explained variance (%)') # Label the y axis.
plt.xlabel('Principal component') # Label the x axis.
#plt.title('Scree plot - Top 10 PCs') # Give a title to the scree plot.
plt.show()
```

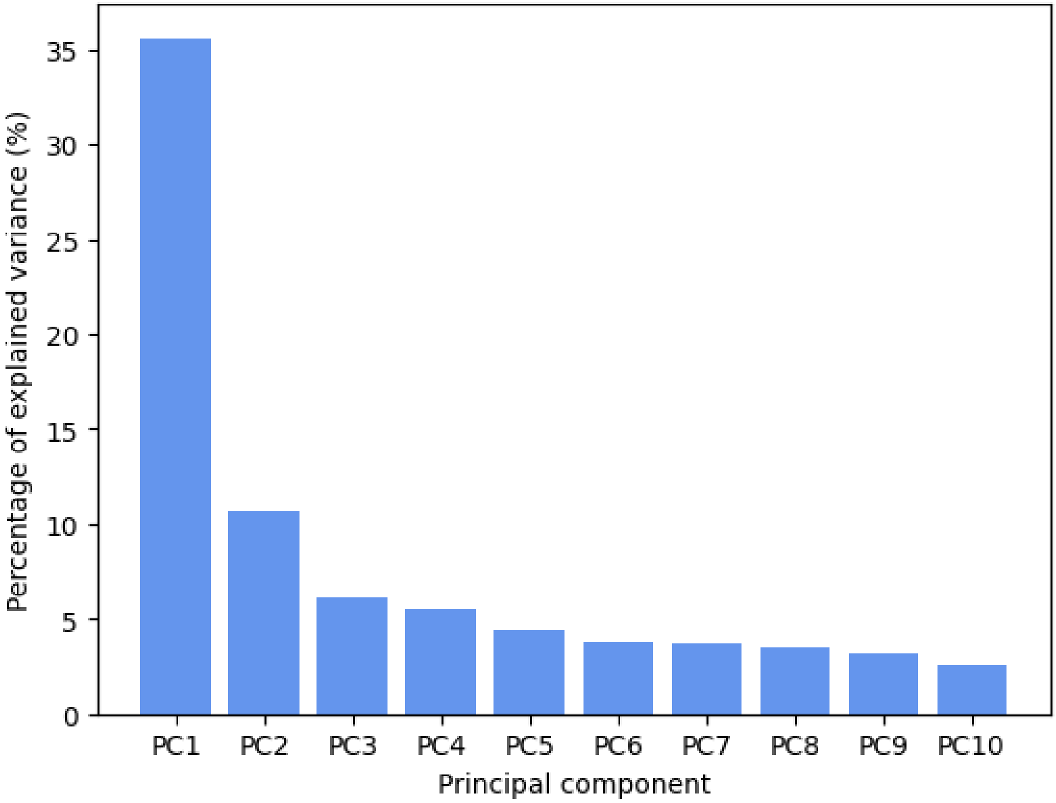


[28]:
```
# Create a pca dataframe to store the PCA data
pca_df = pd.DataFrame(pca_data, index=meta.index, columns=labels)

# Join metadata to the pca dataframe
pca_df = pca_df.join(meta)
```

[43]:
```
# PCA plot colored by period
plt.figure(dpi=100)
```

```
sns.scatterplot(data=pca_df, x = 'PC1', y = 'PC2', hue='period',␣
↪palette='tab10', s=80)
plt.title('PCA plot (colored by period)', fontsize=16) # Title the PCA plot
plt.xlabel(f'PC1 ({pca_var[0]}%)', fontsize=14) # Label the x axis (PC1)
plt.ylabel(f'PC2 ({pca_var[1]}%)', fontsize=14) # Label the y axis (PC2)
plt.xticks(fontsize=12)
plt.yticks(fontsize=12)
plt.legend(loc='lower right', title='Period')
plt.show()
```

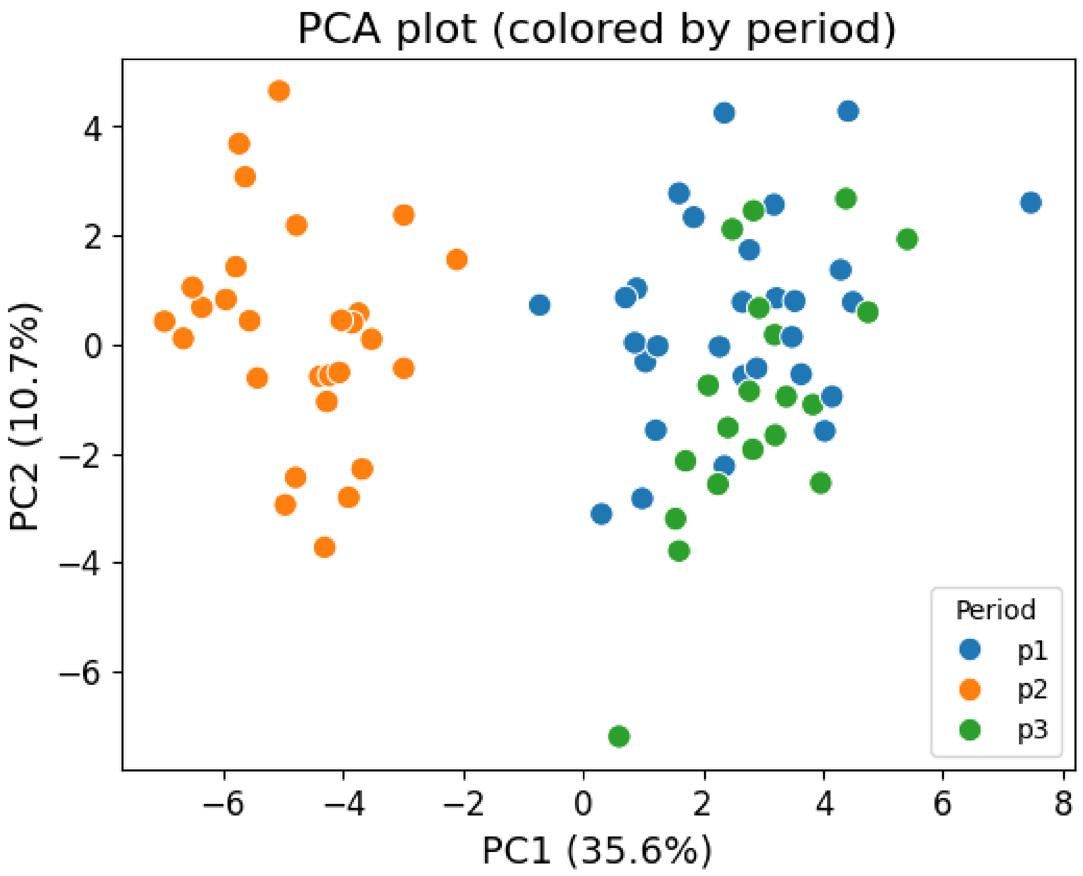


[45]:
```
# Plot colored by shift
plt.figure(dpi=100)
sns.scatterplot(data=pca_df, x = 'PC1', y = 'PC2', hue='shift',␣
↪palette='viridis', s=80)
plt.title('PCA plot (colored by shift)', fontsize=18) # Title the PCA plot
plt.xlabel(f'PC1 - {pca_var[0]}%', fontsize=16) # Label the x axis (PC1)
plt.ylabel(f'PC2 - {pca_var[1]}%', fontsize=16) # Label the y axis (PC2)
plt.xticks(fontsize=14)
plt.yticks(fontsize=14)
```

```
plt.legend(loc='lower right', title='Shift')
plt.show()
```

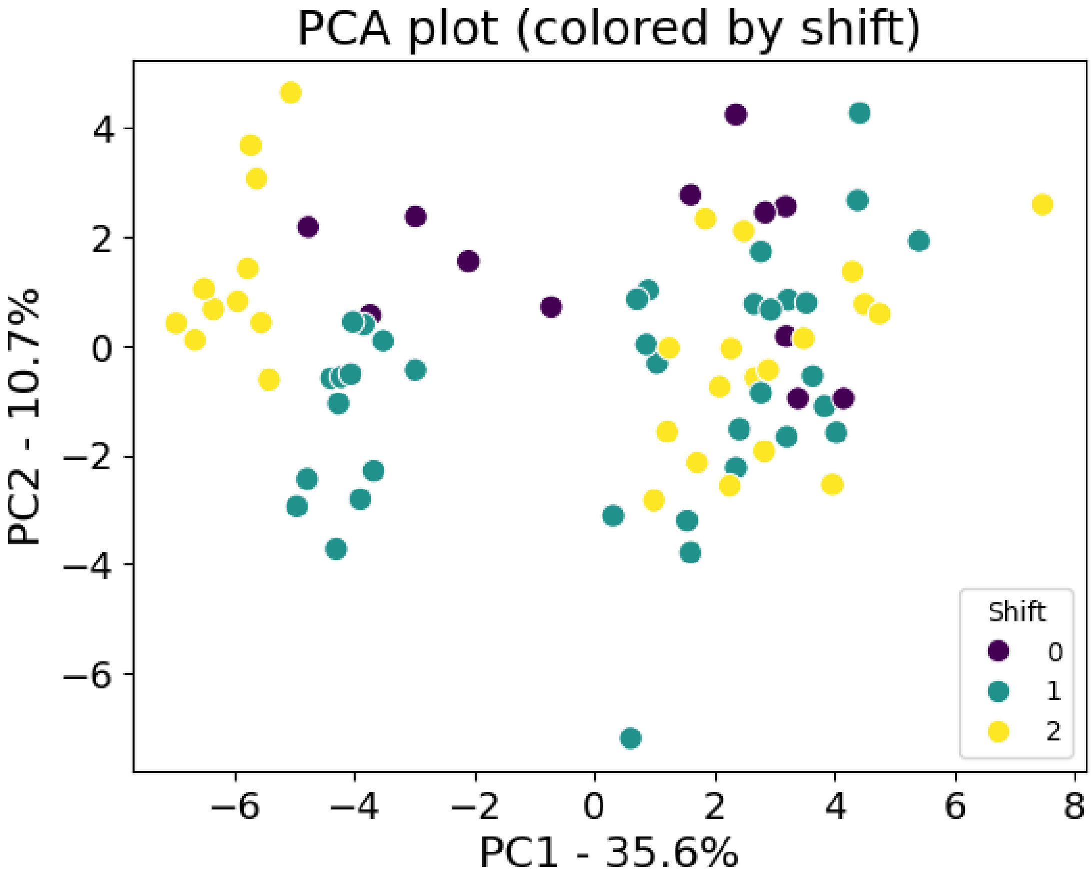


5. Generate a list of the 10 highest loading scores in PC1.

[12]:
```
# Generate a series containing all the loading scores for PC1
loading_scores = pd.Series(pca.components_[0], index=raw.columns)

# Sort loading scores from highest to lowest
sorted_loading_scores = loading_scores.abs().sort_values(ascending=False)

# Select the 10 highest values in the sorted_loading_scores
top_10_features = sorted_loading_scores[0:10].index.values

# Show the top 10
loading_scores[top_10_features]
```

[12]:
```
tat_100lux       0.238978
forger_r        -0.234805
```

```
tat_500lux      0.229523
jewett_r       -0.226964
forger_slope   -0.225058
RA              0.222704
M10_light       0.214474
jewett_slope   -0.213856
hannaytp_r     -0.213772
hannaysp_r     -0.206513
dtype: float64
```

##### 1.4 Monte Carlo Permutation Testing

A permutation test can be performed by randomly shuffling the labels to generate a null distribution. We refer to our approach as a **Monte Carlo permutation test**, since the total number of possible label permutations is far too large to compute exhaustively. Take, for example, the `period` factor, which has three possible labels ($P1$, $P2$ and $P3$). To preserve the repeated-measures design, we permute these labels within each experimental unit (i.e., within each submariner). So, for each single individual, there are:

$$\frac{3!}{(3-3)!} = \frac{3 \times 2 \times 1}{0!} = 6$$

possible permutations. With 30 submariners, this yields $6^{30} \approx 2.21 \times 10^{23}$ total permutations - an impractically large number. Instead, we use a manageable subset ($10^4$) of permutations to approximate the null distribution.

As mentioned, for the `period` factor, we shuffle labels within each submariner, ensuring that the repeated-measures nature of the data is preserved. Regarding the `shift` factor, we performed random permutations within each study period block. Since submariners were not organized in shifts in $P1$ and $P3$, $P1$ serves as a baseline - any statistically significance differences detected here are attributable to random variation -, while $P3$ allows the assessment of the recovery response in the post-mission period.

###### 1.4.1 Computing observed F-values and estimating null distributions

Observed F-values are compared against a null distribution estimated via a Monte Carlo permutation test. This procedure enables the computation of a false positive rate (FPR), which is subsequently adjusted for multiple comparisons using the Benjamini-Hochberg (BH) procedure. The corrected FPR is used in the initial filtering step, in conjunction with the estimated minimum effect size detectable with a sample size of 29 participants (as determined using `G*Power`; see the main manuscript for details).

**By shift**

[63]:

```
def process_feature_shift(df, feature, n_permutations):
    shifts = [group[feature].dropna().values for _, group in df.
 ↪groupby('shift')]

    # Compute observed F statistic and P-value
    f_obs_shifts = stats.f_oneway(*shifts)[0]

    # Create list to store null-distribution F for periods and shifts
    f_perm_shifts = []

    # Permutation test for the "period" factor
    for _ in range(n_permutations):
        # Permutate shift labels
        df_perm = df.copy()
        df_perm['shift'] = df_perm['shift'].sample(frac=1).values

        # Compute F statistics (for shift)
        shifts = [group[feature].dropna().values for _, group in df_perm.
 ↪groupby('shift')]
        f_stat = stats.f_oneway(*shifts)[0]
        f_perm_shifts.append(f_stat)

    print(f'Completed feature: {feature}')
    return {
        'feature': feature,
        'obs_shift': f_obs_shifts,
        'null_shift': f_perm_shifts,
    }
```

[64]:
```
def run_shift(df, n_permutations):
    # List features
    features = df.columns.to_list()
    features.remove('sb_id')
    features.remove('period')
    features.remove('shift')

    # Define the number of cores to use in parallel
    n_cores = max(1, multiprocessing.cpu_count() - 1)
    print(f'Running with {n_cores} cores.')

    # Start time
    start_time= time.time()

    # Process features in parallel
    results = Parallel(n_jobs=n_cores)(
        delayed(process_feature_shift)(df, feature, n_permutations)
        for i, feature in enumerate(features)
```

```
    )

    # Calculate execution time
    dt = time.time() - start_time
    print(f'Time to complete task: {dt:.2f} seconds.')

    # Create dataframes to store results
    observed_df = pd.DataFrame()
    null_dist_df = pd.DataFrame()

    # Iterate over result dictionary
    for result in results:
        # Get the feature
        feature = result['feature']

        # Add observed values
        observed_df[f'{feature}'] = pd.Series(result['obs_shift'])

        # Add null distribution values
        null_dist_df[f'{feature}'] = pd.Series(result['null_shift'])

    return null_dist_df, observed_df
```

[67]:

```
# Save the files
per_list =  ['p1', 'p2', 'p3']
for per in per_list:
    # Open file containing raw values for all variables
    raw = pd.read_csv(os.path.join('..', 'data', 'processed', 'combined_df.
 ↪csv'))
    raw = raw[raw['period'] == per]

    fpath_null = os.path.join('..', 'data', 'processed',␣
 ↪'f_null_dist',f'f_null_dist_shift_{per}.csv')
    fpath_obs = os.path.join('..', 'data', 'processed',␣
 ↪'f_obs',f'obs_f_vals_shift_{per}.csv')

    null_df, obs_df = run_shift(raw, 10000)
    null_df.to_csv(fpath_null, index=False)
    obs_df.to_csv(fpath_obs, index=False)
```

```
Running with 15 cores.
Time to complete task: 39.93 seconds.
Running with 15 cores.
Time to complete task: 39.58 seconds.
Running with 15 cores.
Time to complete task: 41.66 seconds.
```

**By period**

[68]:
```
def process_feature_period(df, feature, n_permutations):
    periods = [group[feature].dropna().values for _, group in df.
 ↪groupby('period')]

    # Compute observed F statistic and P-value
    f_obs_periods = stats.f_oneway(*periods)[0]

    # Create list to store null-distribution F for periods and shifts
    f_perm_periods = []

    # Permutation loop
    for _ in range(n_permutations):
        # Permutate period labels (repeated measure)
        df_perm = df.copy()
        period_perm = df.groupby('sb_id')['period'].transform(lambda sb: sb.
 ↪sample(frac=1).values)
        df_perm['period'] = period_perm

        # Compute F statistics for period
        periods = [group[feature].dropna().values for _, group in df_perm.
 ↪groupby('period')]
        f_stat = stats.f_oneway(*periods)[0]
        f_perm_periods.append(f_stat)

    print(f'Completed feature: {feature}')
    return {
        'feature': feature,
        'obs_period': f_obs_periods,
        'null_period': f_perm_periods,
    }
```

[69]:
```
def run_period(df, n_permutations):
    # List features
    features = df.columns.to_list()
    features.remove('sb_id')
    features.remove('period')
    features.remove('shift')

    # Define the number of cores to use in parallel
    n_cores = max(1, multiprocessing.cpu_count() - 1)
    print(f'Running with {n_cores} cores.')

    # Start time
    start_time= time.time()

    # Process features in parallel
```

```
    results = Parallel(n_jobs=n_cores)(
        delayed(process_feature_period)(df, feature, n_permutations)
        for i, feature in enumerate(features)
    )

    # Calculate execution time
    dt = time.time() - start_time
    print(f'Time to complete task: {dt:.2f} seconds.')

    # Create dataframes to store results
    observed_df = pd.DataFrame()
    null_dist_df = pd.DataFrame()

    # Iterate over result dictionary
    for result in results:
        # Get the feature
        feature = result['feature']

        # Add observed values
        observed_df[f'{feature}'] = pd.Series(result['obs_period'])

        # Add null distribution values
        null_dist_df[f'{feature}'] = pd.Series(result['null_period'])

    return null_dist_df, observed_df
```

[70]:
```
# Save the files
shifts =  ['all', 0, 1, 2]
for shift in shifts:
    raw = pd.read_csv(os.path.join('..', 'data', 'processed', 'combined_df.
 ↪csv'))

    if shift != 'all':
        raw = raw[raw['shift'] == shift]

    fpath_null = os.path.join('..', 'data', 'processed', 'f_null_dist',␣
 ↪f'f_null_dist_period_s{shift}.csv')
    fpath_obs = os.path.join('..', 'data', 'processed', 'f_obs',␣
 ↪f'obs_f_vals_period_s{shift}.csv')

    null_df, obs_df = run_period(raw, 10000)
    null_df.to_csv(fpath_null, index=False)
    obs_df.to_csv(fpath_obs, index=False)
```

```
Running with 15 cores.
Time to complete task: 168.12 seconds.
Running with 15 cores.
```

```
Time to complete task: 75.55 seconds.
Running with 15 cores.
Time to complete task: 106.73 seconds.
Running with 15 cores.
Time to complete task: 99.03 seconds.
```

**Calculate false positive rates (FPR) from null distributions**

[124]:
```
def get_fpr(null_dist, obs_stat, n_perm):
    """
    null_dist (dataframe containing a null distribution of f values generated␣
↪by permutations) - dataframe
    obs_stat (dataframe containing the observed f value) - dataframe
    n_perm (number of permutations previously used) - int
    """
    null_dist = np.array(null_dist)
    obs_stat = obs_stat.iloc[0]

    if np.isnan(obs_stat):
        return np.nan

    tol = max(1e-14, abs(obs_stat) * 1e-14)
    fpr = (np.count_nonzero(null_dist >= (obs_stat - tol)) + 1) / (n_perm + 1)
    return fpr
```

[125]:
```
def get_fpr_df(groups, main_factor, n_perm, list_to_save):
    """
    Groups (list containing all the groupings of the second factor) - list
    Main_factor (main factor being analyzed through permutation analysis) -␣
↪string
    n_perm (number of permutations previously used) - int
    list_to_save (indicate in which list the dataframes should be stored)

    """
    # Run pipeline for main factor, considering groupings of another
    for group in groups:
        # Open null distribution dataframe (generated by permutations)
        null_df = pd.read_csv(os.path.join(null_base,␣
↪f'f_null_dist_{main_factor}_{group}.csv'))

        # Open dataframe containing observed F values
        obs_df = pd.read_csv(os.path.join(obs_base,␣
↪f'obs_f_vals_{main_factor}_{group}.csv'))

        # List all the features
        features = null_df.columns.to_list()
```

```python
        # Create empty dataframes to store FDR values
        fpr_df = pd.DataFrame()

        fvals = []

        # Iterate over features
        for feature in features:
            # Compute the FDR
            fpr = get_fpr(null_df[feature], obs_df[feature], n_perm)

            # Save FDR-values
            fpr_df[feature] = pd.Series(fpr)

            # Retrieve fvals by order
            fvals.append(obs_df[feature].values[0])

        # Transpose the dataframe and rename new columns
        fpr_df = fpr_df.T.reset_index().rename(columns={0: 'fpr', 'index':␣
↪'feature'})

        # Identify the factor used in the analysis
        fpr_df['factor'] = main_factor

        # Identify which shifts were included in the analysis
        fpr_df['used'] = f'{group}'

        # Label metric type (in this case, a F-value)
        fpr_df['metric_type'] = 'f'

        # Store metric val
        fpr_df['metric'] = fvals

        # Save
        list_to_save.append(fpr_df)
```

[126]:
```python
# Create base path to retrieve csv files
null_base = os.path.join('..',  'data', 'processed', 'f_null_dist')
obs_base = os.path.join('..', 'data', 'processed', 'f_obs')

# Create lists to select specific groupings of secondary factor
shifts =  ['all']
periods =  ['p1', 'p2', 'p3']

# Create empty list to store dataframes
to_concat = []

# Obtain dataframes containing fpr values
```

```
get_fpr_df(groups = shifts, main_factor = 'period', n_perm = 10000,␣
 ↪list_to_save = to_concat)
get_fpr_df(groups = periods, main_factor = 'shift', n_perm = 10000,␣
 ↪list_to_save = to_concat)

# Concatenate all dataframes in the list
fpr_combined = pd.concat(to_concat, ignore_index=True)

# Save as a csv file
fpr_combined.to_csv(os.path.join('..', 'data', 'processed', 'fpr', 'f_fpr.
 ↪csv'), index=False)
```

##### Correct FPR for multiple comparisons using the BH procedure

[145]:
```
# Retrieve the dataframe contaning FPR values
ffpr = pd.read_csv(os.path.join('..', 'data', 'processed', 'fpr', 'f_fpr.csv'))
```

[146]:
```
# Since we're comparing within each secondary factor, corrections will happen␣
 ↪for each included period
levels = ['all','p1','p2','p3']

for level in levels:
    # Filter the dataframe to only include fpr values from each level being used
    used = ffpr['used'] == level

    # Adjust for multiple comparisons using the BH procedure
    ffpr.loc[used, 'fpr_bh'] = stats.false_discovery_control(ffpr.
 ↪loc[used,'fpr'], method='bh')

# Save new file containing adjusted FPR values
ffpr.to_csv(os.path.join('..', 'data', 'processed', 'fpr', 'f_fpr_adjusted.
 ↪csv'), index=False)
```

##### Convert the F-statistic to Cohen's F

[128]:
```
# Using Pingouin
def eta_sqr(df, feature, factor):
    """
    Compute eta_sqr (metric of effect size) for one-way anova (fow).
    df --> pd.DataFrame containing all the raw feature data.
    feature --> String indicating the feature being studied.
    factor --> factor to perform comparisons between levels.
    """
    # Perform fow using the Pingouin library
    fow = pg.anova(dv=feature, between=factor, data=df, detailed=True,␣
 ↪effsize='n2')
```

```
    # Return the eta_sqr (´n2´)
    return fow.round(5)['n2'][0]

def cohens_f(eta_sqr):
    """
    Compute cohens_f from eta_sqr
    eta_sqr --> np.float
    """
    return np.sqrt(eta_sqr/(1 - eta_sqr))

def effect_me(df, factor, used):
    """
    df --> pd.DataFrame - df containing all the raw feature data.
    factor --> String - factor to perform comparisons between levels.
    clock_genes --> Bool - it indicates whether clock genes should be included;␣
 ↪if false, they are excluded.
    used --> String - levels of secondary factor used in the comparison of the␣
 ↪primary factor.
    """

    # List all features within the dataframe
    features = df.columns.to_list()

    # Remove non-feature columns (or irrelevant features)
    rm_list = ['sb_id', 'period', 'shift']

    # Remove specified columns
    for item in rm_list:
        if item in features:
            features.remove(item)

    # Create list to store results
    results = []

    for feature in tqdm(features, desc='Processing features'):
        eta_squared = eta_sqr(df, feature, factor)
        results.append({
            'feature': feature,
            'metric_type': 'f',
            'factor': factor,
            'used': used,
            'eta_sqr': eta_squared,
            'cohens_f': cohens_f(eta_squared)
        })

    return pd.DataFrame(results)
```

**Calculate effect size (eta squared and Cohens' F)**

[152]:
```
# Inclusion list (list of secondary factor data included)
level_dict =  {'p1':'shift', 'p2': 'shift','p3':'shift', 'all': 'period'}

# Create a empty list to store effect size dataframes
df_list = []

for key in level_dict.keys():
    # Open file containing raw values for all features and filter
    raw = pd.read_csv(os.path.join('..', 'data', 'processed', 'combined_df.
 ↪csv'))

    if level_dict[key] == 'shift':
        # Filter data within the period block to compare shifts
        raw = raw[raw['period'] == key]
        # Append the newly generated dataframe to the df_list
        df_list.append(effect_me(raw, level_dict[key], key))

    else:
        # Append the newly generated dataframe to the df_list
        df_list.append(effect_me(raw, level_dict[key], key))

# Concatenate the list of all effect size dataframes into a single one
eff_df = pd.concat(df_list)

# Open dataframe containing adjusted FPR values
df_fpr = pd.read_csv(os.path.join('..', 'data', 'processed', 'fpr',␣
 ↪'f_fpr_adjusted.csv'))
```

```
Processing features: 100%|      | 41/41 [00:00<00:00, 250.40it/s]
Processing features: 100%|      | 41/41 [00:00<00:00, 167.46it/s]
Processing features: 100%|      | 41/41 [00:00<00:00, 170.51it/s]
Processing features: 100%|      | 41/41 [00:00<00:00, 157.29it/s]
```

[153]:
```
# Merge dataframe based on common columns
df_fpr = pd.merge(df_fpr, eff_df, on=['feature', 'factor', 'used',␣
 ↪'metric_type'], how='outer')
df_fpr.to_csv(os.path.join('..', 'data', 'processed', 'fpr',␣
 ↪'f_fpr_adjusted_effsize.csv'), index=False)
```

[155]:
```
# Save the effect size dataframe, as well.
directory = os.path.join('..','data','processed','fpr', 'effect_size')
os.makedirs(directory, exist_ok=True)
eff_df.to_csv(os.path.join(directory,'effect_sizes.csv'), index=False)
```

##### 1.4.2 Receiver Operating Characteristic (ROC) curves

Following the initial filtering step, the disriminatory ability of the selected actigrapy features is evaluated through post hoc pairwise comparisons using receiver operating characteristic (ROC) curves. Each features is treated as a univariate classifier, and its ability to distinguish between pairs of shifts and study periods is quantified using the area under the ROC curve (AUC).

Import the required functions to construct ROC curves and calculate the AUC from `sklearn.metrics`.

[2]:
```
from sklearn.metrics import roc_curve, auc
```

**Plotting functions**

[44]:
```
def roc_me(df, group_name, exclude, feature):
    """
    Args:
        df (pd.DataFrame): dataframe with raw feature data
        group_name (str): a string with the factor being used for grouping;␣
↪either shift or period
        feature (str): name of the feature for which the roc curve is being␣
↪drawn
        exclude (list): list with the order in which a group within a factor is␣
↪removed for comparisons two-by-two

    Returns:
        output (dict): a dictionary containing the fpr, tpr and auc for each␣
↪pairwise contrast
    """
    # Define dataframes for two by two comparison
    df_12 = df[~(df[group_name] == exclude[0])].copy()
    df_23 = df[~(df[group_name] == exclude[1])].copy()
    df_13 = df[~(df[group_name] == exclude[2])].copy()

    # Helper function to encode the positive class dynamically
    def encode_positive(sub_df, group_name=group_name, score_col=feature):
        # Get unique groups in the subset
        groups = sub_df[group_name].unique()
        # Compute mean score for each group
        means = {g: sub_df.loc[sub_df[group_name] == g, score_col].mean() for g␣
↪in groups}
        # Identify the group with the highest mean score
        positive_group = max(means, key=means.get)
        # Create a binary label where the positive group gets 1 and the other␣
↪gets 0
        sub_df['binary_label'] = (sub_df[group_name] == positive_group).
↪astype(int)
```

```
        return sub_df

    # Apply dynamic encoding for each pairwise comparison
    df_12 = encode_positive(df_12)
    df_23 = encode_positive(df_23)
    df_13 = encode_positive(df_13)

    # Collect true binary labels
    y_true_12 = df_12['binary_label'].values
    y_true_23 = df_23['binary_label'].values
    y_true_13 = df_13['binary_label'].values

    # Continuous variable (variable being test for the capacity to discriminate␣
↪the two groups)
    y_scores_12 = df_12[feature].values
    y_scores_23 = df_23[feature].values
    y_scores_13 = df_13[feature].values

    # Compute ROC curve
    fpr_12, tpr_12, _ = roc_curve(y_true_12, y_scores_12)
    fpr_23, tpr_23, _ = roc_curve(y_true_23, y_scores_23)
    fpr_13, tpr_13, _ = roc_curve(y_true_13, y_scores_13)

    # Compute AUC
    auc_12 = auc(fpr_12, tpr_12)
    auc_23 = auc(fpr_23, tpr_23)
    auc_13 = auc(fpr_13, tpr_13)

    # return a dataframe with fpr, tpr and auc
    output = {
        'fpr': [fpr_12, fpr_23, fpr_13],
        'tpr': [tpr_12, tpr_23, tpr_13],
        'auc': [auc_12, auc_23, auc_13]
    }
    return output
```

[43]:
```
def draw_roc(fpr,tpr,auc,feature, group_name, labels, data_included):
    """
    Args:
        fpr (list): fpr obtained from roc_me function
        tpr (list): tpr obtained from roc_me function
        auc (list): auc obtained from roc_me function
        group_name (str): a string with the factor being used for grouping;␣
↪either shift or period
        exclude (list): list containing the order in which a group within a␣
↪factor is removed for comparisons two-by-two - list
        feature (str): name of the feature for which a ROC curve is being drawn
```

```
        labels (list): labels for the two-by-two comparison legend
        data_included (str): specify which data was included to compute the␣
↪two-by-two comparison)

    Returns:
        None: it draws and saves ROC curves for each actigraphy feature, either␣
↪by shift or by period
    """
    # Create a plt figure with a 300 dpi
    plt.figure(dpi=300)

    # Plot ROC curves for the 3 comparisons
    plt.plot(fpr[0],
             tpr[0],
             label=f'{labels[0]} (AUC = {auc[0]:.2f})',
             linewidth=2,
             #color='r',
             alpha=0.8)
    plt.plot(fpr[1],
             tpr[1],
             label=f'{labels[1]} (AUC = {auc[1]:.2f})',
             linewidth=2,
             #color='yellow',
             alpha=0.8)
    plt.plot(fpr[2],
             tpr[2],
             label=f'{labels[2]} (AUC = {auc[2]:.2f})',
             linewidth=2,
             #color='teal',
             alpha=0.8)

    # Plot the chance line
    plt.plot([0,1], [0,1], color='black', alpha=0.9, linestyle="--",␣
↪label='chance', linewidth=2)

    # Label the plot
    plt.xlabel('False Positive Rate', fontsize=14)
    plt.ylabel('True Positive Rate', fontsize=14)
    plt.xticks(fontsize=12)
    plt.yticks(fontsize=12)
    plt.title(feature, fontsize=16)
    plt.legend(loc='lower right', frameon=True, fontsize=12)

    # Create the output directory if it does not already exist
    output_dir = os.path.join('..', 'data', 'processed','rocs', group_name,␣
↪f'{data_included}')
    if not os.path.exists(output_dir):
```

```
        os.makedirs(output_dir)

    # Show and save figure on the output directory
    plt.savefig(os.path.join(output_dir, f'{feature}.png'))
    plt.show()
```

[42]:
```
def roc_my_world(df, group_name, exclude, labels, data_included):
    """
    roc_my_world computes roc curves for the entire feature set ('my world').␣
↪Specifically, it calls roc_me and draw_roc
    for all features whitin a given factor (group) vector of interest.

    Args:
        df (pd.DataFrame): dataframe containing raw actigraphy feature data
        group_name (str): factor being used for grouping; either shift or␣
↪period)
        labels (list): labels for the two-by-two comparison legend
        roc_me (function): function to compute roc curve parameters
        draw_roc (function): function to draw the roc curve based on roc_me␣
↪parameters
        data_included (str): specify which data was included in the two-by-two␣
↪comparison

    Returns:
        None - it draws and saves ROC curves for each actigraphy feature,␣
↪either by shift or by period
    """
    # Drop categorical variables (sb_id, shift, period) and list continuous␣
↪variables
    feature_set = df.drop(columns=['sb_id', 'shift', 'period']).columns.
↪to_list()

    for feature in feature_set:
        # Check for and remove any nan rows within a given feature vector
        if any(df[feature].isna()):
            clean_col = df[[group_name, feature]].dropna()
            results = roc_me(clean_col, group_name, exclude, feature)
            draw_roc(results['fpr'], results['tpr'], results['auc'], feature,␣
↪group_name, labels, data_included)
        else:
            results = roc_me(df, group_name, exclude, feature)
            draw_roc(results['fpr'], results['tpr'], results['auc'], feature,␣
↪group_name, labels, data_included)

        # Adjust layout and display the figure
        plt.tight_layout()
```

```
        plt.show()
```

**Draw ROC curves by shift**

[ ]:
```
# List data to include in the analysis of shifts (P1 XOR P2 XOR P3)
data_to_include =  ['p1', 'p2', 'p3']

# Order list to exclude one shift at a time for two-by-two comparisons
excl_shift = [2,0,1]

# Labels for two-by-two shift comparisons
labels_shifts = ['S0 vs. S1', 'S1 vs. S2', 'S0 vs. S2']

# Include all the specified data in the analysis
for data in data_to_include:
    # Open file containing raw values for all variables
    raw = pd.read_csv(os.path.join('..', 'data', 'processed', 'combined_df.
 ↪csv'))

    # Filter only data to include
    raw = raw[raw['period'] == data]

    # Generate ROC curves
    roc_my_world(df=raw, group_name='shift', exclude=excl_shift,␣
 ↪labels=labels_shifts, data_included=data)
```

**Draw ROC curves by period**

[ ]:
```
# List data to include in the analysis of periods (S0 OR S1 OR S2)
data_to_include =  ['all']

# Order list to exclude one period at a time for two-by-two comparisons
excl_period = ['p3', 'p1', 'p2']

# Labels for two-by-two shift comparisons
labels_periods = ['P1 vs. P2', 'P2 vs. P3', 'P1 vs. P3']

for data in data_to_include:
    raw = pd.read_csv(os.path.join('..', 'data', 'processed', 'combined_df.
 ↪csv'))

    if data != 'all':
        raw = raw[raw['shift'] == data]

    # Generate ROC curves
```

```
roc_my_world(df=raw, group_name='period', exclude=excl_period,␣
↪labels=labels_periods, data_included=data)
```

**Determining the significance of the AUC using a Monte Carlo permutation test**

**Common functions to calculate AUC significance** Import library to perform pairwise permutations - this uses the `itertools` module instead of performing the permutations using the `sample` method as we did earlier. Either solution is appropriate, but this might be the most practical solution on the long term.

[8]:
```
import itertools
```

[9]:
```
def calculate_roc(df, group_col, feature):
    # Drop any nans in the received dataframe
    df = df[[group_col, feature]].copy()

    # Helper function to encode the positive class dynamically
    def encode_positive(sub_df, group_col=group_col, score_col=feature):
        # Get unique groups in the subset
        groups = sub_df[group_col].unique()
        # Compute mean score for each group
        means = {g: sub_df.loc[sub_df[group_col] == g, score_col].mean() for g␣
↪in groups}
        # Identify the group with the highest mean score
        positive_group = max(means, key=means.get)
        # Create a binary label where the positive group gets 1 and the other␣
↪gets 0
        sub_df['binary_label'] = (sub_df[group_col] == positive_group).
↪astype(int)
        return sub_df

    # Apply dynamic encoding for each pairwise comparison
    df = encode_positive(df)

    # Collect true binary labels
    y_true = df['binary_label'].values

    # Continuous variable (variable being test for the capacity to discriminate␣
↪the two groups)
    y_scores = df[feature].values

    # Compute ROC curve
    fpr, tpr, _ = roc_curve(y_true, y_scores)

    # Compute AUC
```

```python
    auc_value = auc(fpr, tpr)

    # return a dataframe with fpr, tpr and auc
    output = {
        'fpr': fpr,
        'tpr': tpr,
        'auc': auc_value
    }
    return output
```

**Functions to determine AUC significance (by shift)**

[10]:
```python
def process_shift(df, feature, rm_shift, n_permutations):
    # Create a copy of the original dataframe
    df_obs = df.copy().dropna(subset=[feature])

    # Remove one shift for two-by-two comparison
    df_obs = df_obs[~(df_obs['shift'] == rm_shift)]

    # Calculate observed AUC
    auc_obs_shift = calculate_roc(df_obs, 'shift', feature)['auc']

    # Create empty list to store AUC values (shift)
    auc_perm_shifts = []

    for _ in range(n_permutations):
        # Permutate shift labels
        df_perm = df.copy().dropna(subset=[feature])

        # Exclude shift label (two-by-two comparison)
        df_perm = df_perm[~(df_perm['shift'] == rm_shift)]

        # Permutate the shift labels
        df_perm['shift'] = df_perm['shift'].sample(frac=1).
↪reset_index(drop=True).values

        # Calculate permutation AUC
        auc_perm = calculate_roc(df_perm, 'shift', feature)['auc']
        auc_perm_shifts.append(auc_perm)

    # Return the observed and null distribution
    return {
        'feature': feature,
        'observed_shift': auc_obs_shift,
        'null_shift': auc_perm_shifts,
    }
```

[18]:
```
def run_shift(df, rm_shift, n_permutations):
    # List features
    features = df.columns.to_list()

    # Remove categorical variable columns
    features.remove('sb_id')
    features.remove('period')
    features.remove('shift')

    # Define the number of cores to use in parallel
    n_cores = max(1, multiprocessing.cpu_count() - 2)
    print(f'Running with {n_cores} cores.')

    # Start time
    start_time= time.time()

    # Process features in parallel
    results = Parallel(n_jobs=n_cores)(
        delayed(process_shift)(df, feature, rm_shift, n_permutations)
        for i, feature in enumerate(features)
    )

    # Calculate execution time
    dt = time.time() - start_time
    print(f'Time to complete task: {dt:.2f} seconds.')

    # Create dataframes to store results
    observed_df = pd.DataFrame()
    null_dist_df = pd.DataFrame()

    # Iterate over result dictionary
    for result in results:
        # Get the feature
        feature = result['feature']

        # Add observed values
        observed_df[f'{feature}'] = pd.Series(result['observed_shift'])

        # Add null distribution values
        null_dist_df[f'{feature}'] = pd.Series(result['null_shift'])

    return null_dist_df, observed_df
```

##### Functions to determine AUC significance (by period)

[19]:
```
def process_period(df, feature, rm_period, n_permutations):
    # Create a copy of the original dataframe
    df_obs = df.copy().dropna(subset=[feature])

    # Exclude period label (for two-by-two comparison)
    df_obs = df_obs[~(df_obs['period'] == rm_period)]

    # Get AUC (period)
    auc_obs_period = calculate_roc(df_obs, 'period', feature)['auc']

    # Create empty lists to store AUC (period)
    auc_perm_periods = []

    for _ in range(n_permutations):
        # Permutate period labels (repeated measures)
        #df_perm = df.copy().dropna(subset=[feature]).reset_index(drop=True)
        df_perm = df.copy().dropna(subset=[feature]).reset_index(drop=True)

        # Exclude one period for two-by-two comparisons
        df_perm = df_perm[~(df_perm['period'] == rm_period)]

        # Permutate period labels
        period_perm = df_perm.groupby('sb_id')['period'].transform(lambda sb:␣
↪sb.sample(frac=1).values)
        df_perm['period'] = period_perm

        # Calculate permutated AUC
        auc_perm = calculate_roc(df_perm, 'period', feature)['auc']

        # Append AUC
        auc_perm_periods.append(auc_perm)

    # Return the observed and null distribution
    return {
        'feature': feature,
        'observed_period': auc_obs_period,
        'null_period': auc_perm_periods,
    }
```

[20]:
```
def run_period(df, rm_period, n_permutations):
    # List features
    features = df.columns.to_list()

    # Remove categorical variable columns
    features.remove('sb_id')
    features.remove('period')
    features.remove('shift')
```

```
    # Define the number of cores to use in parallel
    n_cores = max(1, multiprocessing.cpu_count() - 2)
    print(f'Running with {n_cores} cores.')

    # Start time
    start_time= time.time()

    # Process features in parallel
    results = Parallel(n_jobs=n_cores)(
        delayed(process_period)(df, feature, rm_period, n_permutations)
        for i, feature in enumerate(features)
    )

    # Calculate execution time
    dt = time.time() - start_time
    print(f'Time to complete task: {dt:.2f} seconds.')

    # Create dataframes to store results
    observed_df = pd.DataFrame()
    null_dist_df = pd.DataFrame()

    # Iterate over result dictionary
    for result in results:
        # Get the feature
        feature = result['feature']

        # Add observed values
        observed_df[f'{feature}'] = pd.Series(result['observed_period'])

        # Add null distribution values
        null_dist_df[f'{feature}'] = pd.Series(result['null_period'])

    return null_dist_df, observed_df
```

##### Run AUC significance pipeline for shifts

[22]:
```
# Shift
per_list =  ['p1', 'p2', 'p3']
for per in per_list:
    # Open file containing raw values for all variables
    raw = pd.read_csv(os.path.join('..', 'data', 'processed', 'combined_df.
↪csv'))
    raw = raw[raw['period'] == per]
    raw = raw.dropna(axis=1, how='all')

    # List of exclusions (to perform two by two comparisons)
```

```
    exclusion_list_shift = [2,0,1]

    for i in exclusion_list_shift:
        idx_shift = [0,1,2]

        # Remove one shift
        idx_shift.remove(i)

        # Run parallel pipeline
        print(f'running with {per} data and with shifts {idx_shift[0]} and␣
↪{idx_shift[1]}')
        null_df, obs_df = run_shift(raw,i, 10000)

        # Save results to csv in output_dir
        os.makedirs(os.path.join('..', 'data', 'processed', 'auc_null_dist'),␣
↪exist_ok=True)
        os.makedirs(os.path.join('..', 'data', 'processed', 'auc_obs'),␣
↪exist_ok=True)
        null_df.to_csv(os.path.join('..', 'data', 'processed',␣
↪'auc_null_dist',f'auc_null_shift_{per}_rm_s{i}.csv'))
        obs_df.to_csv(os.path.join('..', 'data', 'processed',␣
↪'auc_obs',f'obs_auc_shift_{per}_rm_s{i}.csv'))
```

```
running with p1 data and with shifts 0 and 1
Running with 14 cores.
Time to complete task: 104.48 seconds.
running with p1 data and with shifts 1 and 2
Running with 14 cores.
Time to complete task: 106.49 seconds.
running with p1 data and with shifts 0 and 2
Running with 14 cores.
Time to complete task: 107.41 seconds.
running with p2 data and with shifts 0 and 1
Running with 14 cores.
Time to complete task: 118.74 seconds.
running with p2 data and with shifts 1 and 2
Running with 14 cores.
Time to complete task: 108.15 seconds.
running with p2 data and with shifts 0 and 2
Running with 14 cores.
Time to complete task: 104.67 seconds.
running with p3 data and with shifts 0 and 1
Running with 14 cores.
Time to complete task: 107.85 seconds.
running with p3 data and with shifts 1 and 2
Running with 14 cores.
Time to complete task: 108.60 seconds.
```

```
running with p3 data and with shifts 0 and 2
Running with 14 cores.
Time to complete task: 112.35 seconds.
```

**Run AUC significance pipeline for periods**

[23]:
```
# Period
shifts =  ['all']

for shift in shifts:
    raw = pd.read_csv(os.path.join('..', 'data', 'processed', 'combined_df.
↪csv'))
    if shift != 'all':
        raw = raw[raw['shift'] == shift]
    raw = raw.dropna(axis=1, how='all')

    # List of exclusions (to perform two by two comparisons)
    exclusion_list_period = ['p3','p1','p2']

    for i in exclusion_list_period:
        idx_period = ['p1','p2','p3']

        # Remove period and shift from exclusion list
        idx_period.remove(i)

        print(f'running with shifs = {shift} data and with periods␣
↪{idx_period[0]} and {idx_period[1]}')

        # Run parallel pipeline
        null_df, obs_df = run_period(raw,i, 10000)

        # Save results to csv
        null_df.to_csv(os.path.join('..', 'data', 'processed',␣
↪'auc_null_dist',f'auc_null_period_{shift}_rm_{i}.csv'))
        obs_df.to_csv(os.path.join('..', 'data', 'processed',␣
↪'auc_obs',f'obs_auc_period_{shift}_rm_{i}.csv'))
```

```
running with shifs = all data and with periods p1 and p2
Running with 14 cores.
Time to complete task: 282.70 seconds.
running with shifs = all data and with periods p2 and p3
Running with 14 cores.
Time to complete task: 271.79 seconds.
running with shifs = all data and with periods p1 and p3
Running with 14 cores.
Time to complete task: 296.84 seconds.
```

Compute FPR for AUC

[24]:
```python
# Same function as for the FDR for the F-statistic
def get_fpr(null_dist, obs_stat, n_perm):
    """
    null_dist (dataframe containing a null distribution of f values generated␣
 ↪by permutations) - dataframe
    obs_stat (dataframe containing the observed f value) - dataframe
    n_perm (number of permutations previously used) - int
    """
    null_dist = np.array(null_dist)
    obs_stat = obs_stat.iloc[0]

    if np.isnan(obs_stat):
        return np.nan

    tol = max(1e-14, abs(obs_stat) * 1e-14)
    fpr = (np.count_nonzero(null_dist >= (obs_stat - tol)) + 1) / (n_perm + 1)
    return fpr
```

[25]:
```python
def get_fpr_df(groups, exclusion_list, main_factor, n_perm, list_to_save):
    """
    Groups (list containing all the groupings of the second factor) - list
    exclusion_list (sequentially ordered list to exclude one level of the␣
 ↪factor for two-by-two comparisons) - list
    Main_factor (main factor being analyzed through permutation analysis) -␣
 ↪string
    n_perm (number of permutations previously used) - int
    list_to_save (indicate in which list the dataframes should be stored)

    """
    # Create base path to retrieve csv files
    null_base = os.path.join('..',  'data', 'processed', 'auc_null_dist')
    obs_base = os.path.join('..', 'data', 'processed', 'auc_obs')

    # Run pipeline for main factor, considering groupings of another
    for group in groups:
        for excluded in exclusion_list:
            # Open null distribution dataframe (generated by permutations)
            null_df = pd.read_csv(os.path.join(null_base,␣
 ↪f'auc_null_{main_factor}_{group}_rm_{excluded}.csv'))

            # Open dataframe containing observed F values
            obs_df = pd.read_csv(os.path.join(obs_base,␣
 ↪f'obs_auc_{main_factor}_{group}_rm_{excluded}.csv'))

            # List all the features and remove the first column (index)
            features = null_df.columns.to_list()
            del features[0]
```

```
        # Create empty dataframes to store FDR values
        fpr_df = pd.DataFrame()

        aucs = []

        # Iterate over features
        for feature in features:
            # Compute the FDR
            fpr = get_fpr(null_df[feature], obs_df[feature], n_perm)

            # Save FDR-values
            fpr_df[feature] = pd.Series(fpr)

            # Retrieve aucs by order
            aucs.append(obs_df[feature].values[0])

        # Transpose the dataframe and rename new columns
        fpr_df = fpr_df.T.reset_index().rename(columns={0: 'fpr', 'index':␣
↪'feature'})

        # Identify the factor used in the analysis
        fpr_df['factor'] = main_factor

        # Identify which levels of the secondary factor were included in␣
↪the analysis
        fpr_df['used'] = f'{group}'

        # Label metric type (in this case, a F-value)
        fpr_df['metric_type'] = 'auc'

        # Store metric val
        fpr_df['metric'] = aucs

        # Label levels within the main factor being compared
        pair = None
        match main_factor:
            case 'period':
                pair = ['p1','p2','p3']
            case 'shift':
                pair = ['s0','s1','s2']
        pair.remove(excluded)
        fpr_df['levels'] = f'{pair[0]}_{pair[1]}'

        # Save
        list_to_save.append(fpr_df)
```

[26]:
```python
# Create lists to select specific groupings of secondary factor
shifts =  ['all']
periods =  ['p1', 'p2', 'p3']

# List of exclusions (to perform 2-by-2 comparisons)
exclusions_period = ['p3','p1','p2']
exclusions_shift = ['s2','s0','s1']

# Create empty list to store dataframes
to_concat = []

# Obtain dataframes containing fpr values
get_fpr_df(groups = shifts, exclusion_list=exclusions_period, main_factor =␣
↪'period', n_perm = 10000, list_to_save = to_concat)
get_fpr_df(groups = periods, exclusion_list=exclusions_shift, main_factor =␣
↪'shift', n_perm = 10000, list_to_save = to_concat)

# Concatenate all dataframes in the list
fpr_combined = pd.concat(to_concat, ignore_index=True)

# Save as a csv file
fpr_combined.to_csv(os.path.join('..', 'data', 'processed', 'fpr',␣
↪'auc_fpr_combined.csv'), index=False)
```

#### 1.5 Figures and Summary Statistics

This section contains the code used to generate all figures presented in the main manuscript and supplementary materials. It also includes scripts for computing summary statistics, namely the mean and standard deviation of each actigraphy features, grouped by either shift or study period.

##### 1.5.1 Import the required libraries and data

1. In order to add insets to plots, such as in the case of the volcano plots, we need to import the functions `inset_axes` and `mark_inset` from `mpl_toolkits.axes_grid1.inset_locator`.

[28]:
```python
from mpl_toolkits.axes_grid1.inset_locator import inset_axes, mark_inset
```

2. Open the raw actigraphy-derived feature values to derive summary statistics.

[29]:
```python
raw = pd.read_csv(os.path.join('..', 'data', 'processed', 'combined_df.csv'))
```

3. Open the dataframe which contains all relevant statistical results on the F-statistic, including the false positive rate (FPR) estimated via permutation testing, the FDR-adjusted FPR obtained using the Benjamini-Hochberg (BH) procedure. Effect size metrics were also computed.

[30]:
```
df = pd.read_csv(os.path.join('..', 'data', 'processed', 'fpr',␣
↪'f_fpr_adjusted_effsize.csv'))
```

4. Open the dataframe that contains the information regarding AUC significance testing.

[31]:
```
dfauc = pd.read_csv(os.path.join('..', 'data', 'processed', 'fpr',␣
↪'auc_fpr_combined.csv'))
```

##### 1.5.2 Summary statistics

Calculate mean and standard deviation (either by shift or period) for each actigraphy feature.

###### By period

[ ]:
```
# Calculate descriptives by period (e.g., mean, std)
mean_raw = raw.groupby(by=['period']).describe(include=np.number).T.round(2)
```

###### By shift

[ ]:
```
# Filter only data by shift from the mission period (P2)
raw_mission = raw[raw['period'] == 'p2']

# Calculate the descriptives by shift (e.g., mean, std).
mean_raw = raw_mission.groupby(by=['shift']).describe(include=np.number).T.
↪round(2)
```

###### Statistically significant features by period (comparing shifts within each study period)

[ ]:
```
# Since we are comparing shifts, we need to select rows labeled with the shift␣
↪factor
df_shift = df[df['factor'] == 'shift']
```

[34]:
```
# Determine the number of features deemed capable of distinguishing at least␣
↪one shift during the pre-mission period (P1)
p1 = df_shift[(df_shift['used'] == 'p1') & (df_shift['fpr_bh'] < 0.05)]
print(f'In P1, the number of features capable of distinguishing at least one␣
↪shift was {len(p1)}.')
```

```
In P1, the number of features capable of distinguishing at least one shift was
0.
```

[35]:
```
# Determine the number of features deemed capable of distinguishing at least␣
↪one shift during the mission period (P2)
p2 = df_shift[(df_shift['used'] == 'p2') & (df_shift['fpr_bh'] < 0.05)]
print(f'In P2, the number of features capable of distinguishing at least one␣
↪shift was {len(p2)}.')
```

```
p2
```

In P2, the number of features capable of distinguishing at least one shift was 17.

[35]:

| | feature | factor | used | metric_type | metric | fpr | fpr_bh |
|---|---|---|---|---|---|---|---|
| 22 | L5 | shift | p2 | f | 13.922523 | 0.000300 | 0.002050 |
| 26 | L5_light | shift | p2 | f | 8.068482 | 0.002900 | 0.008492 |
| 46 | RA | shift | p2 | f | 35.262997 | 0.000100 | 0.000820 |
| 50 | SRI | shift | p2 | f | 20.230465 | 0.000100 | 0.000820 |
| 58 | activity_bouts | shift | p2 | f | 5.522662 | 0.012899 | 0.033053 |
| 62 | all_sleep_bouts | shift | p2 | f | 13.308254 | 0.001200 | 0.004472 |
| 66 | amplitude | shift | p2 | f | 7.450009 | 0.002200 | 0.006938 |
| 70 | exp_level | shift | p2 | f | 14.058354 | 0.000400 | 0.002343 |
| 78 | forger_slope | shift | p2 | f | 10.760116 | 0.000800 | 0.003280 |
| 98 | jewett_r | shift | p2 | f | 6.958469 | 0.005499 | 0.015032 |
| 110 | kRA | shift | p2 | f | 5.147488 | 0.014999 | 0.036173 |
| 114 | main_sleep_bout | shift | p2 | f | 31.671579 | 0.000100 | 0.000820 |
| 122 | minor_sleep_bouts | shift | p2 | f | 13.879122 | 0.000100 | 0.000820 |
| 130 | mlit_10lux | shift | p2 | f | 12.105244 | 0.000800 | 0.003280 |
| 138 | sleep_midpoint | shift | p2 | f | 24.722132 | 0.000100 | 0.000820 |
| 142 | tat_100lux | shift | p2 | f | 11.075624 | 0.001800 | 0.006149 |
| 146 | tat_10lux | shift | p2 | f | 11.222269 | 0.000700 | 0.003280 |

\

| | eta_sqr | cohens_f |
|---|---|---|
| 22 | 0.52692 | 1.055371 |
| 26 | 0.39227 | 0.803410 |
| 46 | 0.73829 | 1.679590 |
| 50 | 0.61809 | 1.272171 |
| 58 | 0.30643 | 0.664692 |
| 62 | 0.51566 | 1.031826 |
| 66 | 0.37343 | 0.772004 |
| 70 | 0.52934 | 1.060507 |
| 78 | 0.46260 | 0.927799 |
| 98 | 0.35761 | 0.746114 |
| 110 | 0.29168 | 0.641710 |
| 114 | 0.71701 | 1.591758 |
| 122 | 0.52614 | 1.053721 |
| 130 | 0.49198 | 0.984087 |
| 138 | 0.66418 | 1.406338 |
| 142 | 0.46979 | 0.941300 |
| 146 | 0.47307 | 0.947515 |

[36]:
```
# Determine the number of features deemed capable of distinguishing at least␣
↪one shift during the post-mission period (P3)
p3 = df_shift[(df_shift['used'] == 'p3') & (df_shift['fpr_bh'] < 0.05)]
```

```
print(f'In P3, the number of features capable of distinguishing at least one␣
↪shift was {len(p3)}.')
```

```
In P3, the number of features capable of distinguishing at least one shift was
0.
```

**Statistically significant features by shift**

[8]:
```
alls = df[(df['used'] == 'all') & (df['fpr_bh'] < 0.05) & (df['cohens_f'] > 0.
↪63)]
no_features_all = len(alls)
print(f'The number of features capable of distinguishing at least one period␣
↪was {no_features_all}.')
alls
```

```
The number of features capable of distinguishing at least one period was 21.
```

[8]:

| | feature | factor | used | metric_type | metric | fpr | fpr_bh |
|---|---|---|---|---|---|---|---|
| 8 | IS_light | period | all | f | 44.534776 | 0.0001 | 0.000164 |
| 20 | L5 | period | all | f | 22.215794 | 0.0001 | 0.000164 |
| 32 | M10 | period | all | f | 25.748272 | 0.0001 | 0.000164 |
| 36 | M10_light | period | all | f | 65.652967 | 0.0001 | 0.000164 |
| 44 | RA | period | all | f | 74.096462 | 0.0001 | 0.000164 |
| 60 | all_sleep_bouts | period | all | f | 40.746525 | 0.0001 | 0.000164 |
| 64 | amplitude | period | all | f | 28.775499 | 0.0001 | 0.000164 |
| 72 | forger_r | period | all | f | 109.013138 | 0.0001 | 0.000164 |
| 76 | forger_slope | period | all | f | 43.897175 | 0.0001 | 0.000164 |
| 80 | hannaysp_r | period | all | f | 47.758405 | 0.0001 | 0.000164 |
| 88 | hannaytp_r | period | all | f | 72.622990 | 0.0001 | 0.000164 |
| 92 | hannaytp_slope | period | all | f | 19.912030 | 0.0001 | 0.000164 |
| 96 | jewett_r | period | all | f | 100.255423 | 0.0001 | 0.000164 |
| 100 | jewett_slope | period | all | f | 30.543864 | 0.0001 | 0.000164 |
| 112 | main_sleep_bout | period | all | f | 43.804061 | 0.0001 | 0.000164 |
| 120 | minor_sleep_bouts | period | all | f | 23.439469 | 0.0001 | 0.000164 |
| 136 | sleep_midpoint | period | all | f | 18.413178 | 0.0001 | 0.000164 |
| 140 | tat_100lux | period | all | f | 140.890306 | 0.0001 | 0.000164 |
| 144 | tat_10lux | period | all | f | 34.716181 | 0.0001 | 0.000164 |
| 148 | tat_500lux | period | all | f | 105.639899 | 0.0001 | 0.000164 |
| 156 | vat_10lux | period | all | f | 40.546258 | 0.0001 | 0.000164 |

\

| | eta_sqr | cohens_f |
|---|---|---|
| 8 | 0.53634 | 1.075524 |
| 20 | 0.36590 | 0.759630 |
| 32 | 0.40076 | 0.817790 |
| 36 | 0.63035 | 1.305857 |
| 44 | 0.65807 | 1.387291 |
| 60 | 0.51417 | 1.028753 |

| | | |
|---|---|---|
| 64 | 0.42773 | 0.864539 |
| 72 | 0.74152 | 1.693745 |
| 76 | 0.53600 | 1.074789 |
| 80 | 0.55689 | 1.121060 |
| 88 | 0.65649 | 1.382434 |
| 92 | 0.34383 | 0.723875 |
| 96 | 0.72515 | 1.624299 |
| 100 | 0.44561 | 0.896540 |
| 112 | 0.53222 | 1.066657 |
| 120 | 0.37843 | 0.780275 |
| 136 | 0.32353 | 0.691565 |
| 140 | 0.78538 | 1.912955 |
| 144 | 0.47416 | 0.949589 |
| 148 | 0.73290 | 1.656477 |
| 156 | 0.51294 | 1.026224 |

##### 1.5.3 Heatmaps

Create heatmaps depicting features exhibiting near-perfect classification performance (AUC  0.99) in at least one contrast between periods or shifts.

**Pairwise shift comparisons of shift-discriminating features (ranked by AUC)** Given that features deem significant were only found in P2, we will only perform two-by-two comparisons using the significant features identified in this period.

[18]:
```
# Get a list of significant features in P2
dfs = df[df['factor'] == 'shift']
fsl = dfs[(dfs['used'] == 'p2') & (dfs['fpr_bh'] < 0.05)]['feature'].to_list()

# Filter AUC two-by-two comparison results by shift (with period being a␣
 ↪blocking factor)
dfsauc = dfauc[(dfauc['factor'] == 'shift') & (dfauc['used'] == 'p2')]
dfsauc = dfsauc[dfsauc['feature'].isin(fsl)]
```

[19]:
```
dfsaucsig = dfsauc[(dfsauc['fpr'] < 0.05)][['feature', 'levels', 'factor',␣
 ↪'metric']]
dfsaucsig = dfsaucsig.pivot(index='feature', columns='levels', values='metric')
dfsaucsig = dfsaucsig[(dfsaucsig['s0_s1'] >= 0.99) | (dfsaucsig['s0_s2'] >= 0.
 ↪99) | (dfsaucsig['s1_s2'] >= 0.99)]
x_order = ['s0_s1', 's0_s2', 's1_s2']
dfsaucsig = dfsaucsig[x_order]
dfsaucsig = dfsaucsig.loc[dfsaucsig.fillna(0).mean(axis=1).
 ↪sort_values(ascending=False).index]
dfsaucsig = dfsaucsig.T
dfsaucsig.index = ['S0 vs. S1', 'S0 vs. S2', 'S1 vs. S2']
```

[20]:
```
plt.figure(figsize=(14, 1), dpi=100)
sns.reset_defaults()
sns.heatmap(data = dfsaucsig,
            cmap="coolwarm",
            annot=True,
            annot_kws={'size': 9, 'rotation':0},
            vmin = 0.5,
            vmax = 1,
            fmt=".2f",
            linewidths=0.1
           )
#plt.title('Pairwise Shift Comparisons of Discriminating Features Ranked by␣
↪AUC', pad=15)
plt.xticks(fontsize=12)
plt.yticks(fontsize=12)
plt.xlabel('Feature', labelpad=14, fontsize=14)
plt.ylabel('Contrast', labelpad=14, fontsize=14)
plt.show()
```

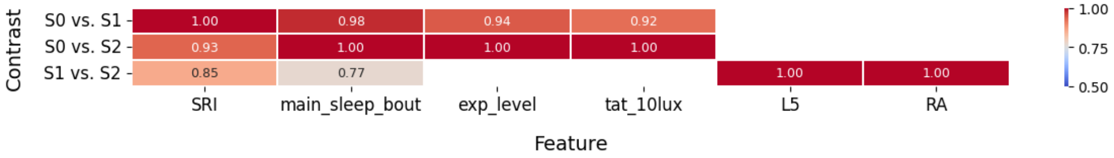


**Pairwise shift comparisons of period-discriminating features (ranked by AUC)**

[21]:
```
# Get a list of significant features in P2
dfp = df[df['factor'] == 'period']
fslp = dfp[(dfp['used'] == 'all') & (dfp['fpr_bh'] < 0.05)]['feature'].to_list()

# Filter AUC two-by-two comparison results by period (with period being a␣
↪blocking factor)
dfpauc = dfauc[(dfauc['factor'] == 'period') & (dfauc['used'] == 'all')]

# Filter features only present in the list of significantly changed features in␣
↪the ANOVA
dfpauc = dfpauc[dfpauc['feature'].isin(fslp)]
```

[22]:
```
dfpaucsig = dfpauc[(dfpauc['fpr'] < 0.05)][['feature', 'levels', 'factor',␣
↪'metric']]
dfpaucsig = dfpaucsig.pivot(index='feature', columns='levels', values='metric')
dfpaucsig = dfpaucsig[(dfpaucsig['p1_p2'] >= 0.99) | (dfpaucsig['p1_p3'] >= 0.
↪99) | (dfpaucsig['p1_p3'] >= 0.99)]
dfpaucsig
```

```
dfpaucsig = dfpaucsig[['p1_p2', 'p2_p3', 'p1_p3']]
dfpaucsig = dfpaucsig.loc[dfpaucsig.fillna(0).median(axis=1).
    ↪sort_values(ascending=False).index]
dfpaucsig = dfpaucsig.T
dfpaucsig.index = ['P1 vs. P2', 'P2 vs. P3', 'P1 vs. P3']
```

[23]:
```
plt.figure(figsize=(14, 1), dpi=100)

g = sns.heatmap(data = dfpaucsig,
            cmap="coolwarm",
            annot=True,
            annot_kws={'size': 10, 'rotation':0},
            vmin = 0.5,
            vmax = 1,
            fmt=".2f",
            linewidths=0.1
           )

plt.xticks(fontsize=12)
plt.yticks(fontsize=12)
plt.xlabel('Feature', labelpad=14, fontsize=14)
plt.ylabel('Contrast', labelpad=14, fontsize=14)
plt.show()
#plt.title('Pairwise Shift Comparisons of Discriminating Features Ranked by␣
    ↪AUC', pad=15)
```

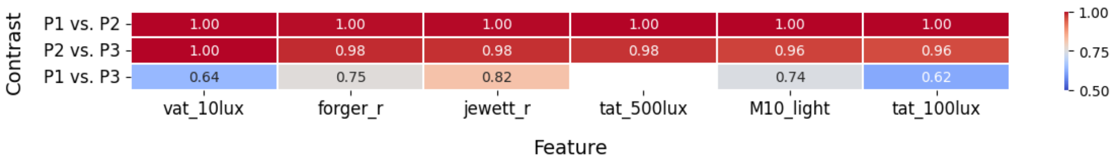


##### Top 10 shift-discriminating features (two-by-two comparison)

[44]:
```
# Top 10 features by comparison
dfsaucsig = dfsauc[(dfsauc['fpr'] < 0.05)][['feature', 'levels', 'factor',␣
    ↪'metric']]

comps = ['s0_s1', 's0_s2', 's1_s2']
outdict = {}

for comp in comps:
    outdict[comp] = dfsaucsig.loc[dfsaucsig[dfsaucsig['levels'] == comp].
    ↪sort_values(by='metric', ascending=False).index]['feature'].to_list()[:10]
```

[45]:
```
outdict['s0_s1']
```

[45]:
```
['SRI',
 'main_sleep_bout',
 'L5_light',
 'exp_level',
 'amplitude',
 'tat_10lux',
 'kRA',
 'mlit_10lux',
 'tat_100lux',
 'all_sleep_bouts']
```

[46]:
```
outdict['s0_s2']
```

[46]:
```
['exp_level',
 'tat_10lux',
 'main_sleep_bout',
 'minor_sleep_bouts',
 'activity_bouts',
 'sleep_midpoint',
 'tat_100lux',
 'SRI',
 'amplitude',
 'kRA']
```

[47]:
```
outdict['s1_s2']
```

[47]:
```
['L5',
 'RA',
 'sleep_midpoint',
 'mlit_10lux',
 'L5_light',
 'minor_sleep_bouts',
 'forger_slope',
 'SRI',
 'jewett_r',
 'main_sleep_bout']
```

##### 1.5.4 Jittered violin plots for Cohen's $f$ and AUC distributions

Larger effect sizes were found to be associated with the distinctions between study periods. To visualize this, we construct jittered violin plots of Cohen's f distributions across features, by shift and period:

[24]:
```
# Open dataframe containing f-stats
df = pd.read_csv(os.path.join('..', 'data', 'processed', 'fpr',␣
↪'f_fpr_adjusted_effsize.csv'))

# Violin plot
sns.reset_defaults()
plt.figure(dpi=100)
sns.violinplot(data=df, x='factor', y='cohens_f', hue='factor', alpha=0.4,␣
↪palette='Blues')

# Overlay points
sns.stripplot(data=df, x='factor', y='cohens_f', color='black', alpha=0.8,␣
↪jitter=True, size=4)

#plt.title('Distribution of Cohen\'s $f$ across Factors', fontsize=16)
plt.ylabel('Cohen\'s $f$', fontsize=14)
plt.xlabel('Factor', fontsize=14)
caps = ['Period', 'Shift']
plt.xticks(ticks=range(len(caps)), labels=caps, fontsize=12)
plt.yticks(fontsize=12)
plt.ylim(bottom=0)
plt.tight_layout()
plt.show()
```

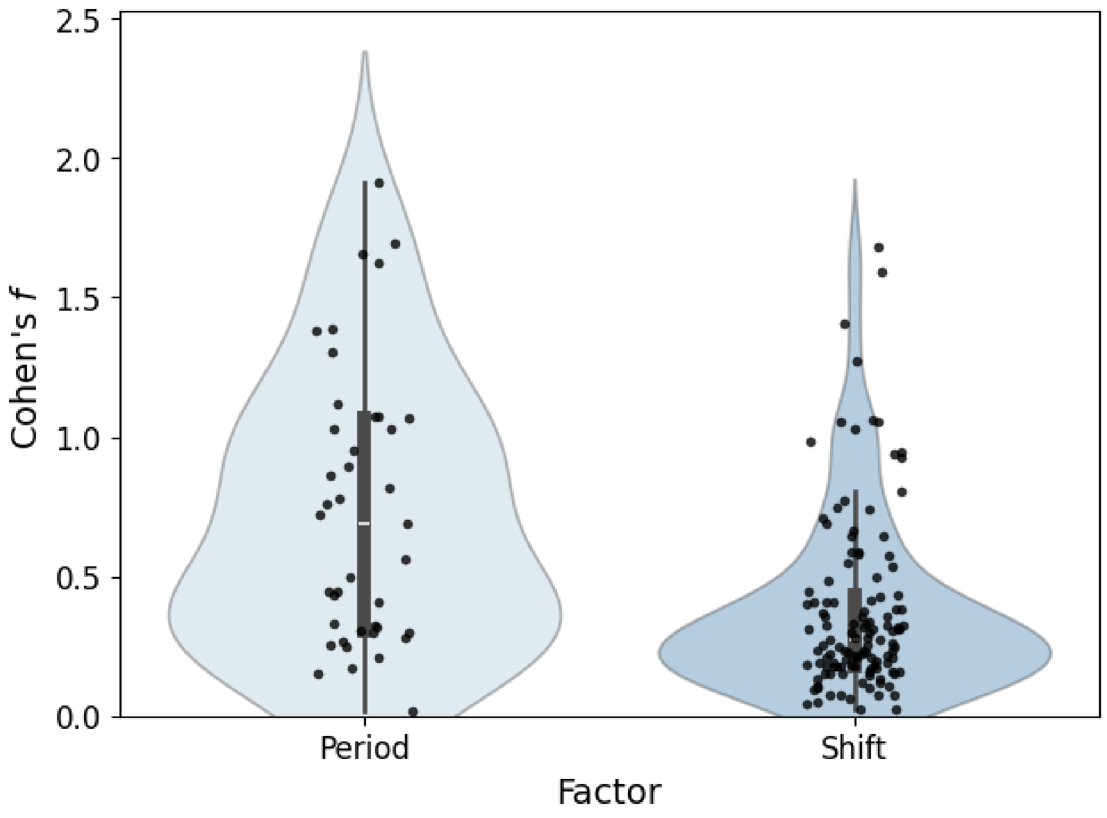

To assess the likelihood of observing a result as extreme as ours under the null hypothesis, we use the non-parametric Mann–Whitney U test (MWU), given that the assumption of normality was not met.

[25]:
```
# Check assumption of normality
pg.normality(data=df, dv='cohens_f', group='factor', method='shapiro', alpha=0.
↪05)
```

[25]:
```
               W          pval  normal
factor
period  0.924712  9.678857e-03   False
shift   0.798385  1.047984e-11   False
```

[26]:
```
# Perform Mann-Whitney U test
pg.mwu(x=df[df['factor'] == 'period']['cohens_f'],
       y=df[df['factor'] == 'shift']['cohens_f'],
       alternative='two-sided')
```

[26]:
```
      U-val alternative     p-val      RBC     CLES
MWU  3747.0   two-sided  0.000003  0.48602  0.74301
```

Next, generate a jittered violin plot with the three contrasts, and perform the Kruskall-Wallis test, with the post-hoc results mentioned in the main manuscript.

[33]:
```
# Open dataframe containing f-stats
df = pd.read_csv(os.path.join('..', 'data', 'processed', 'fpr',␣
↪'auc_fpr_combined.csv'))
df = df[(df['factor'] == 'period') & (df['fpr'] < 0.05)]

# Use classic matplotlib style
sns.reset_defaults()

# Violin plot
plt.figure(dpi=100)
sns.violinplot(data=df, x='levels', y='metric', hue='levels', alpha=0.4,␣
↪palette='Blues')
sns.stripplot(data=df, x='levels', y='metric', color='black', alpha=0.8,␣
↪jitter=True, size=5)

# Capital letters for the contrast
caps = ['P1 vs. P2', 'P2 vs. P3', 'P1 vs. P3']

#plt.title('$AUC$ Magnitude Across Study Periods', fontsize=16)
plt.ylabel('$AUC$', fontsize=14)
plt.ylim([0.5, 1.0])
```

```
plt.xlabel('Contrast', fontsize=14)
plt.xticks(ticks=range(len(caps)), labels=caps, fontsize=12)
plt.yticks(fontsize=12)
plt.tight_layout()
plt.show()
```

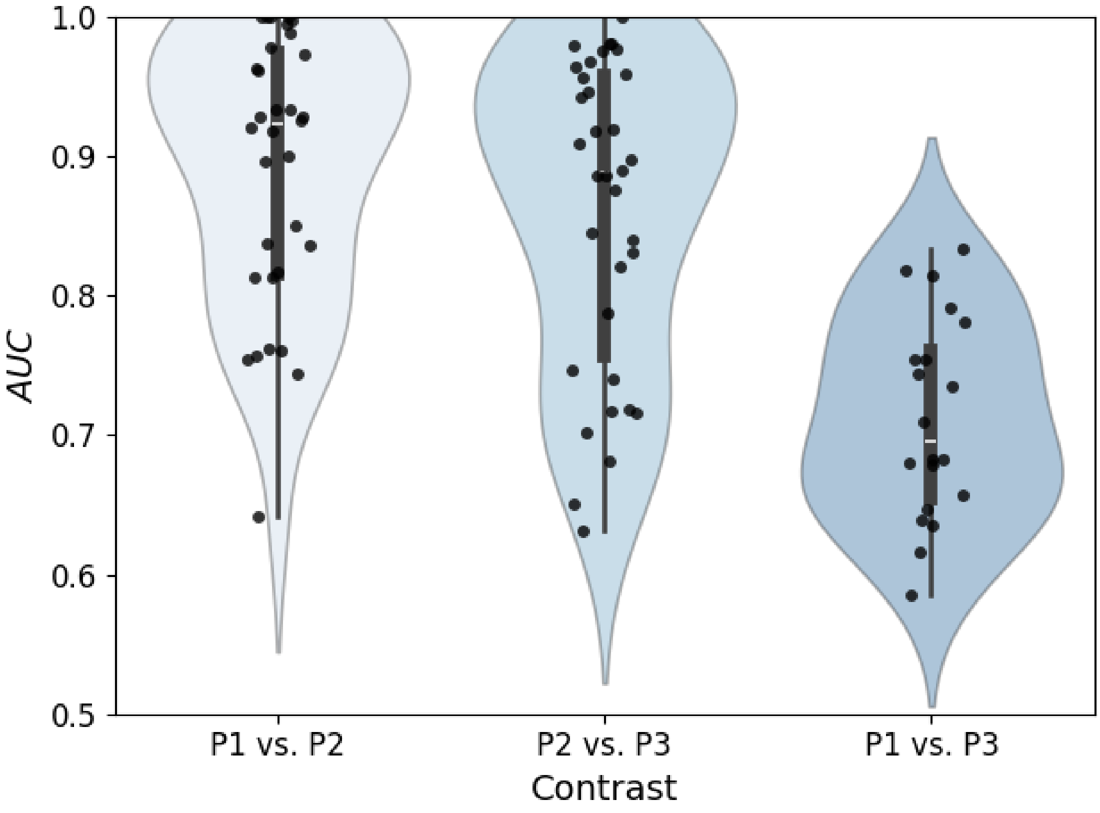


[34]:
```
# Import additional library to perform posthoc_dunn test
import scikit_posthocs as sp
```

[35]:
```
# Let's determine significance
pg.kruskal(data=df, dv='metric', between='levels', detailed=True)
```

[35]:
```
         Source  ddof1          H         p-unc
Kruskal  levels      2  28.036236  8.165987e-07
```

[40]:
```
# Run post-hoc test (Dunn's test)
p1_p2 = df[df['levels'] == 'p1_p2']['metric']
p2_p3 = df[df['levels'] == 'p2_p3']['metric']
p1_p3 = df[df['levels'] == 'p1_p3']['metric']

data = [p1_p2, p2_p3, p1_p3]
```

```
sp.posthoc_dunn(data, p_adjust = 'fdr_bh')
```

[40]:

| | 1 | 2 | 3 |
|---|---|---|---|
| 1 | 1.000000e+00 | 0.22260 | 7.534749e-07 |
| 2 | 2.226003e-01 | 1.00000 | 4.982712e-05 |
| 3 | 7.534749e-07 | 0.00005 | 1.000000e+00 |

##### 1.5.5 Cluster maps

Since we computed a considerable amount of features based on actigraphy expression, we wonder whether the features would correlate. To do this, we calculated a non-pametric (Spermann correlation) correlation matrix and built a cluster map.

[20]:
```
# Open file containing actigraphy and clock gene features
raw = pd.read_csv(os.path.join('..', 'data', 'processed', 'combined_df.csv'))
```

[21]:
```
# Remove non-numeric columns
rm = ['sb_id', 'period', 'shift']
raw = raw.drop(columns=rm)

# Drop rows containing NaNs
raw = raw.dropna(axis='rows', how='all')
```

[33]:
```
# Draw the clustermap
#sns.set(font_scale=0.6)
sns.reset_defaults()

#spearman
plt.figure(dpi=200)
sns.clustermap(raw.corr(method='spearman'),
               method='average',
               vmin=-1,
               vmax=1,
               dendrogram_ratio=(0.2, 0.2),
               figsize=(14, 14),
               cmap='coolwarm',
               linewidth = 0,
               tree_kws={'linewidths':2})
plt.show()
```

<Figure size 1280x960 with 0 Axes>

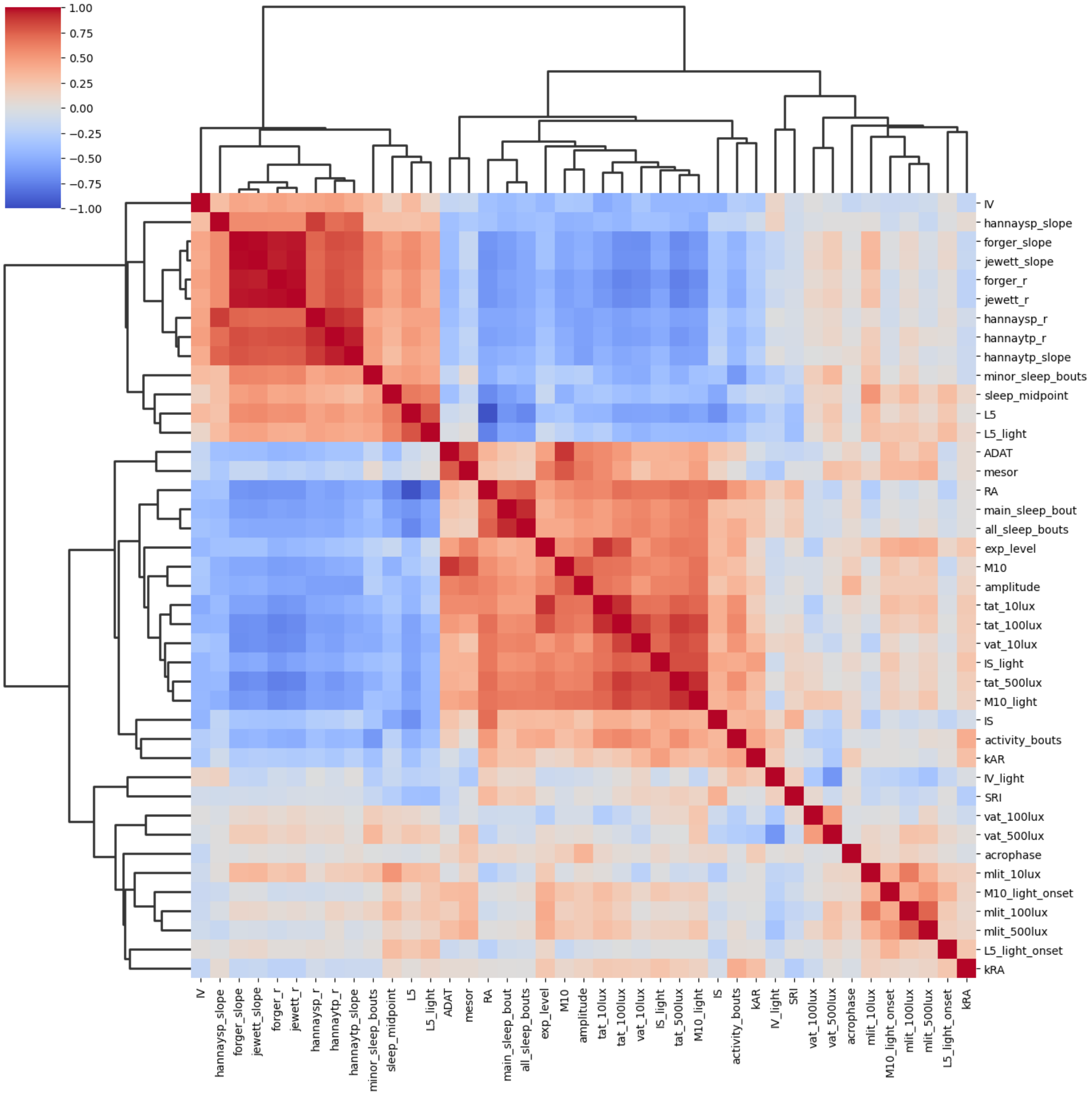


##### 1.5.6 $-log_{10}(fdr) \times Cohen'sf$ scatter plots

A scatter plot is generated to display the magnitude of effect size against statistical significance for features discriminating between shifts and periods. This visualization facilitates the identification of features with both strong effects and low likelihood of arising by chance.

###### Shift-discriminating features (P2)

[19]:
```
# Filter comparisons between shifts from the mission period (p2)
dffp = df.loc[(df['factor'] == 'shift') & (df['metric_type'] == 'f') &
↪(df['used'] == 'p2')].copy()
```

```
# Fixed color pallete for significance
sig_palette = {'FDR < 0.05': 'orange', 'FDR   0.05': 'steelblue'}

# Create a significance labeling for hue
dffp['significance'] = np.where(dffp['fpr_bh'] < 0.05, 'FDR < 0.05', 'FDR   0.
↪05')

# Create a top 10 ranking based on effect size
dffp['cohens_f_rank'] = dffp['cohens_f'].rank()
dffp['top_5'] = np.where(dffp.index.isin(dffp['cohens_f_rank'].nlargest(5).
↪index), 1,0)

# Main plot
fig, ax = plt.subplots(dpi=100)

# Draw a scatterplot, with fixed axis
sns.scatterplot(data = dffp,
                x='cohens_f',
                y= -np.log10(dffp['fpr_bh']),
                hue='significance',
                palette=sig_palette,
                ax=ax,
                alpha=.5,
                s=80)

# Inset (zoomed-in) plot
axins = inset_axes(ax, width="50%", height="45%", loc="lower right",␣
↪bbox_to_anchor=(-0.2, 0.1, 1.6, 1.6), bbox_transform=ax.transAxes)

# Plot the same data in inset
sns.scatterplot(
    data=dffp,
    x='cohens_f',
    y=-np.log10(dffp['fpr_bh']),
    hue='significance',
    alpha=0.5,
    palette=sig_palette,
    ax=axins,
    legend=False,
    s=80
)

# Set limits for zoomed area
axins.set_xlim(1.0, 1.85)
axins.set_ylim(2.55, 3.5)

# Remove axis labels and ticks from inset
```

```
axins.set_xlabel('')
axins.set_ylabel('')
axins.set_xticks([])
axins.set_yticks([])

# Add labels just to top 5 in inset
for i, (_, row) in enumerate(dffp[dffp['top_5'] == 1].iterrows()):
    jitter = (i % 6 - 2) * 0.05
    axins.text(
        row['cohens_f'],
        -np.log10(row['fpr_bh']),
        row['feature'],
        fontsize=12,
        rotation=45
    )

# Mark the zoomed-in area on the main plot
mark_inset(ax, axins, loc1=2, loc2=1, fc="none", ec="0.5")

ax.legend(title='', fontsize=12)
ax.set_xlim([0, 2])
ax.set_ylim([0, 3.5])
ax.set_xlabel('Cohens\'s $f$', fontsize=14)
ax.set_ylabel(r'$-\log_{10}(FDR)$', fontsize=14)
#ax.set_title('Shifts (P2)', fontsize=16)
ax.tick_params(axis='both', labelsize=12)
plt.show()
```

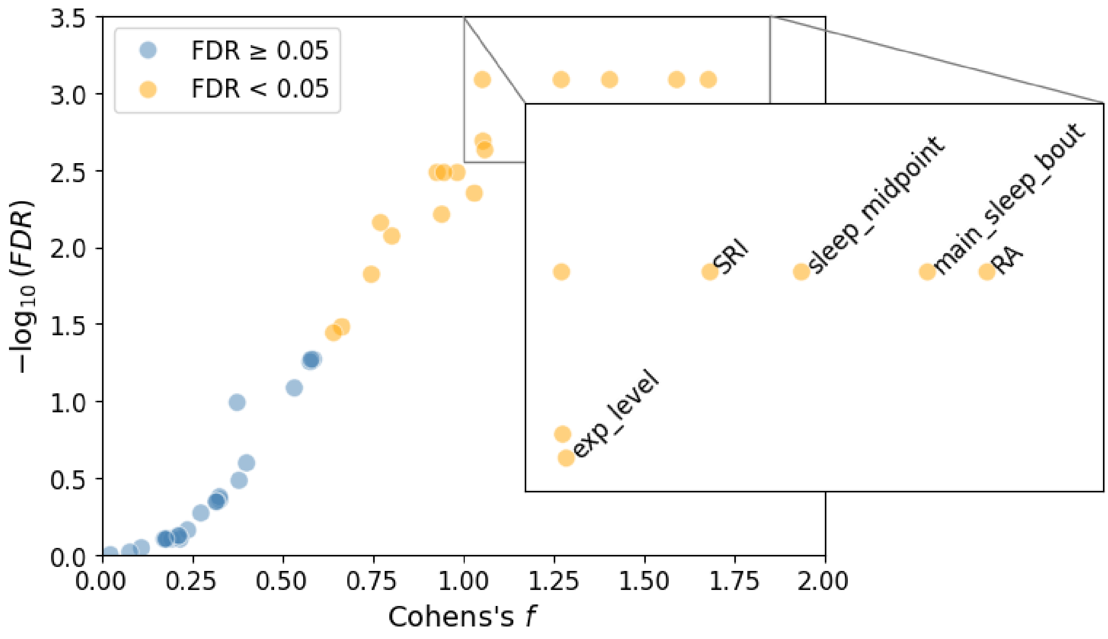

##### Period-discriminating features

[15]:
```
# Import function to create inset axis and to mark it
from mpl_toolkits.axes_grid1.inset_locator import inset_axes, mark_inset

# Fixed color pallete for significance
sig_palette = {'FDR < 0.05': 'orange', 'FDR  0.05': 'steelblue'}

# Open dataframe containing f-stats
df = pd.read_csv(os.path.join('..', 'data', 'processed', 'fpr',␣
 ↪'f_fpr_adjusted_effsize.csv'))

# Filter and compute new columns
dffp = df.loc[(df['factor'] == 'period') & (df['metric_type'] == 'f') &␣
 ↪(df['used'] == 'all')].copy()
dffp['significance'] = np.where(dffp['fpr_bh'] < 0.05, 'FDR < 0.05', 'FDR  0.
 ↪05')
dffp['cohens_f_rank'] = dffp['cohens_f'].rank()
dffp['top_5'] = np.where(dffp.index.isin(dffp['cohens_f_rank'].nlargest(6).
 ↪index), 1, 0)

# Main plot
fig, ax = plt.subplots(dpi=100)
sns.scatterplot(
    data=dffp,
    x='cohens_f',
    y=-np.log10(dffp['fpr_bh']),
    hue='significance',
    alpha=0.5,
    palette=sig_palette,
    ax=ax,
    s=80
)

ax.set_xlim([0, 2])
ax.set_ylim([0, 4])
ax.set_xlabel('Cohens\'s $f$', fontsize=14)
ax.set_ylabel(r'$-\log_{10}(FDR)$', fontsize=14)
#ax.set_title('Period (all shifts)', fontsize=16)

# Inset (zoomed-in) plot
#axins = inset_axes(ax, width="70%", height="40%", loc="center right")
axins = inset_axes(ax, width="70%", height="40%", loc="lower right",␣
 ↪bbox_to_anchor=(0.1, 0.2, 1, 1.1), bbox_transform=ax.transAxes)
```

```python
# Plot the same data in inset
sns.scatterplot(
    data=dffp,
    x='cohens_f',
    y=-np.log10(dffp['fpr_bh']),
    hue='significance',
    alpha=0.5,
    palette=sig_palette,
    ax=axins,
    legend=False,
    s=80
)

# Set limits for zoomed area
axins.set_xlim(1.35, 1.95)
axins.set_ylim(3.68+0.02, 3.94+0.05)

# Remove axis labels and ticks from inset
axins.set_xlabel('')
axins.set_ylabel('')
axins.set_xticks([])
axins.set_yticks([])

# Add labels just to top 6 in inset
for i, (_, row) in enumerate(dffp[dffp['top_5'] == 1].iterrows()):
    if row['feature'] == 'RA':
        axins.text(
            row['cohens_f'] - 0.015,
            -np.log10(row['fpr_bh']) + 0.02,
            row['feature'],
            fontsize=12,
            rotation=90
        )
    elif row['feature'] == 'hannaytp_r':
        axins.text(
            row['cohens_f'] + 0.015,
            -np.log10(row['fpr_bh']) + 0.02,
            row['feature'],
            fontsize=12,
            rotation=90
        )
    else:
        axins.text(
            row['cohens_f'],
            -np.log10(row['fpr_bh']) + 0.02,
            row['feature'],
            fontsize=12,
```

```
            rotation=90
        )

# Mark the zoomed-in area on the main plot
mark_inset(ax, axins, loc1=2, loc2=1, fc="none", ec="0.5")

# Omit legend of main axis legend and adjust font size
ax.tick_params(axis='both', labelsize=12)
ax.legend(title='', fontsize=12)
plt.show()
```

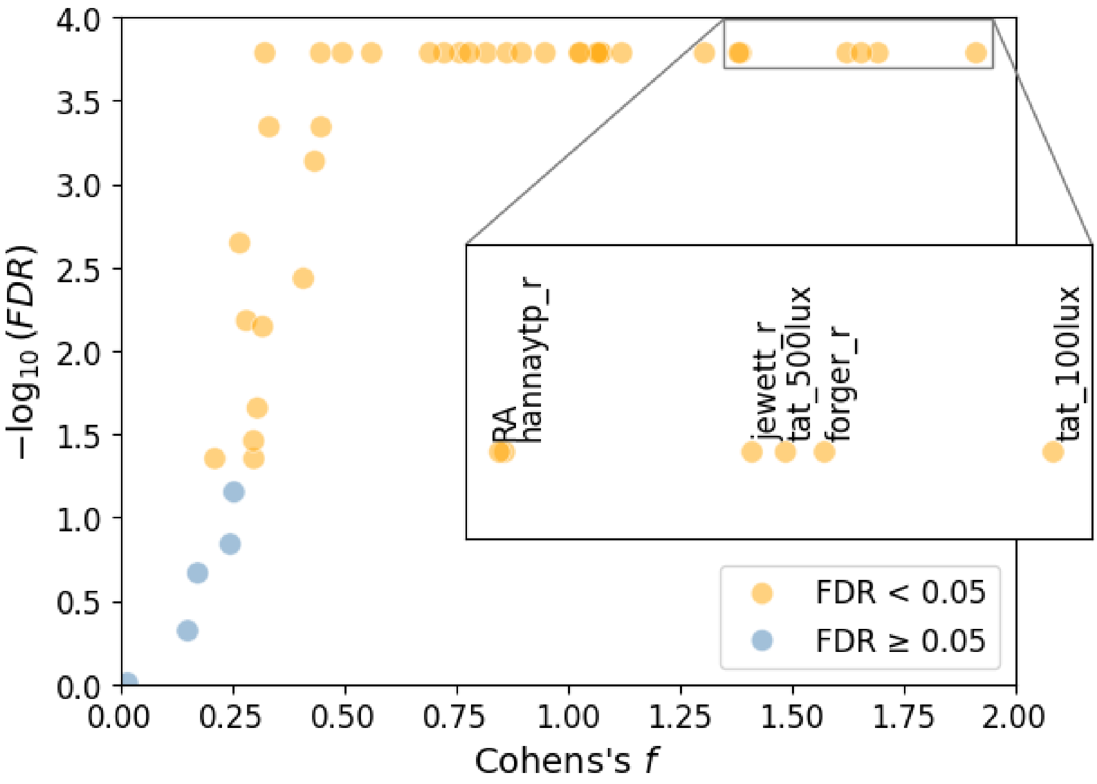


##### 1.5.7 Violin plots (by period and shift)

Draw violin plots for individual actigraphy features. For clarity, we format the $y$-axis tick labels using mathematical notation via `mtick` from `matplotlib.ticker`. We also define a `play_violin` function to easily compute and draw violin plots for each actigraphy feature in the study.

[5]:
```
import matplotlib.ticker as mtick
```

[10]:
```
def play_violin(raw,
                feature,
                unit_label,
                main_factor,
                sec_factor,
                sec_level,
                ylimits=None,
                up_ylim=None,
                low_ylim=None,
                zero_line=False,
                scientific=True,
                conversion_factor=1,
                dpi = 100,
               ):
    """
    Generate a violin plot for a selected actigraphy feature, allowing␣
↪comparisons across levels
    of the primary factor within a specific level of a secondary, blocking␣
↪factor.

    Args:
        raw (pd.DataFrame): DataFrame containing the actigraphy features.
        feature (str): Name of the feature to plot (e.g., "ADAT").
        unit_label (str): Label for the y-axis indicating the feature's unit (e.
↪g., "°C", "lux").
        main_factor (str): Primary factor to compare (e.g., "shift" or␣
↪"period").
        sec_factor (str): Secondary factor used to subset the data (e.g.,␣
↪"shift" or "period").
        sec_level (str or int): The level of the secondary factor to include in␣
↪the comparison.
        ylimits (tuple, optional): Tuple specifying the full y-axis limits␣
↪(min, max), if applicable.
        up_ylim (float, optional): Upper limit for the y-axis; overrides␣
↪`ylimits` if provided.
        low_ylim (float, optional): Lower limit for the y-axis; overrides␣
↪`ylimits` if provided.
        zero_line (bool, optional): Whether to draw a horizontal line at y=0␣
↪(default is False).
        scientific (bool, optional): Whether to format y-axis ticks using␣
↪scientific notation (default is True).
        conversion_factor (float, optional): Factor by which to multiply the␣
↪feature values
                                              (e.g., use 100 to convert␣
↪proportions to percentages).

```

```
    Returns:
        None: Displays a violin plot
    """
    # Check if there is a filtering condition on the secondary factor
    if sec_level != 'all':
        raw = raw[raw[sec_factor] == sec_level]

    # Adjust the number of dots per inch in the figure
    plt.figure(dpi=dpi)

    # Generate a violin plot
    sns.violinplot(data=raw,
                   x=main_factor,
                   y=raw[feature]*conversion_factor,
                   hue=main_factor,
                   palette='Blues',
                   alpha=0.4,
                   linecolor='black',
                   inner_kws={'color': 'black'},
                   legend=False)

    # Plot individual datapoints
    sns.stripplot(data=raw,
                  x=main_factor,
                  y=raw[feature] * conversion_factor,
                  alpha=1,
                  jitter=True,
                  size=6,
                  color='black',
                  legend=False)

    if ylimits != None:
        plt.ylim(ylimits[0], ylimits[1])

    if up_ylim != None:
        plt.ylim(top=up_ylim)

    if low_ylim != None:
        plt.ylim(bottom=low_ylim)

    if zero_line:
        plt.axhline(y=0, color='salmon', linestyle='dashed')

    # Adjust ytick numbers to scientific notation
    if scientific:
        formatter = mtick.ScalarFormatter(useMathText=True)
        formatter.set_scientific(True)
```

```
        formatter.set_powerlimits((-1, 1))
        plt.gca().yaxis.set_major_formatter(formatter)

    if main_factor == 'period':
        caps = ['P1', 'P2', 'P3']
        plt.xticks(ticks=range(len(caps)), labels=caps, fontsize=12)

    # Title
    plt.title(f"{feature}", fontsize=16)

    # Set ylabel and increase the fontsize
    plt.ylabel(ylabel, fontsize=14)

    # Increase the fontsize of the yticks
    plt.yticks(fontsize=12)

    # Set the xlabel and increase the fontsize
    if main_factor == 'period':
        plt.xlabel('Period', fontsize=14)
    if main_factor == 'shift':
        plt.xlabel('Shift', fontsize=14)

    # Increase the fontsize of the yticks
    plt.xticks(fontsize=12)
    plt.yticks(fontsize=12)

    plt.show()
    plt.close()
```

Draw illustrative violin plot for `jewett_r` and `SRI`:

[11]:
```
raw = pd.read_csv(os.path.join('..', 'data', 'processed', 'combined_df.csv'))
ylabel = 'Pearson correlation coefficient'
play_violin(raw, 'jewett_r', ylabel, 'period', 'shift', 'all', ylimits=(-1,1),␣
 ↪zero_line=True)
```

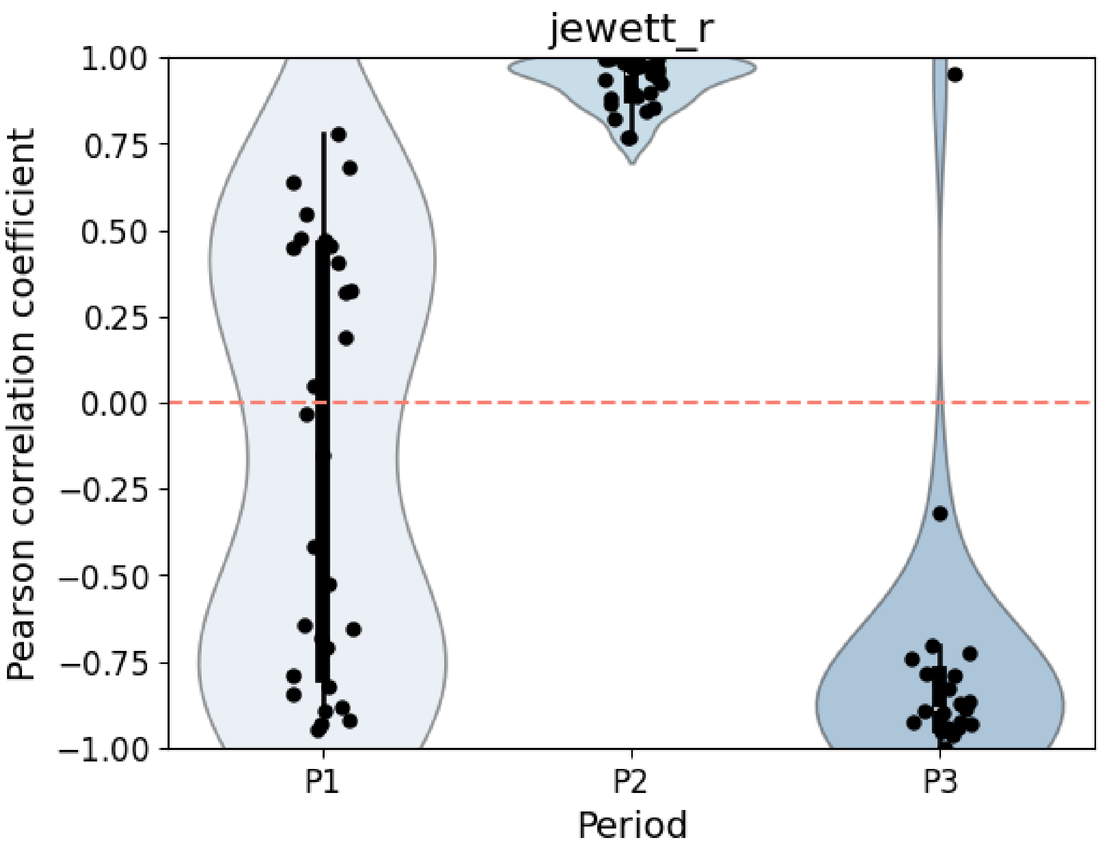


[12]:
```
raw = pd.read_csv(os.path.join('..', 'data', 'processed', 'combined_df.csv'))
ylabel = 'Probability (%)'
play_violin(raw, 'SRI', ylabel, 'shift', 'period', 'p2', scientific=False,␣
 ↪ylimits=(0,100))
```

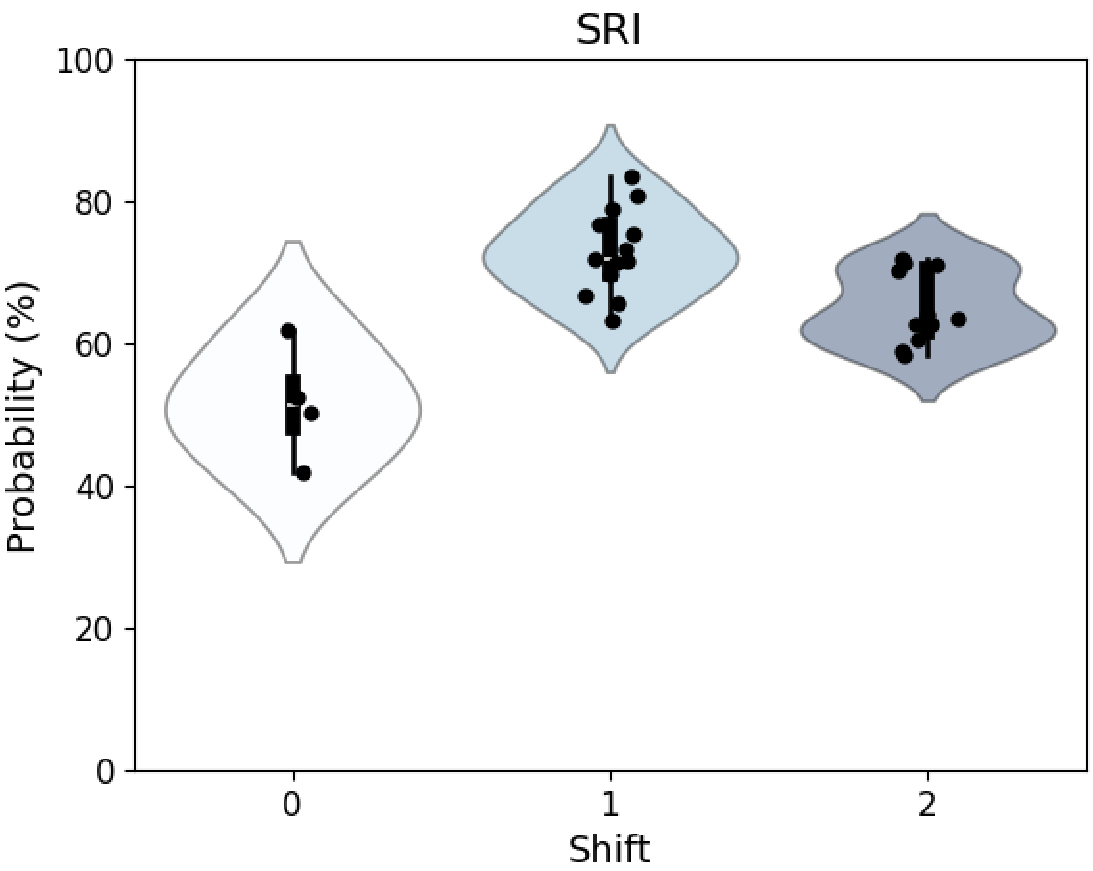
SRI
Probability (%)
100
80
60
40
20
0
0
1
2
Shift
